# GSDME unlocks astrocyte-driven neurotoxicity in Alzheimer’s Disease

**DOI:** 10.1101/2025.08.28.672784

**Authors:** Xueman Xie, Chenhui Ji, Jinmei Xu, Xiaoyang Lu, Guigui Guo, Xiaomin Wu, Wenjing Liu, Yaozhen Chen, Yuanxing Zhang, Jintao Wang, Jianping Li, Xingbin Hu, Shouwen Chen, Gang Wang, Qin Liu

## Abstract

Astrocytic calcium dysregulation and reactivity precede Aβ deposition in amyloid-β deposition in Alzheimer’s disease (AD) but the neurotoxic mechanisms remain unclear. We show that GSDME acts as a switch, linking MAM-mediated calcium release to astrocyte-driven neurotoxicity. Specifically, Aβ-activated microglial signals activate astrocytic GSDME, releasing its N-terminal fragment, which targets MAMs and triggers ER calcium efflux. This induces biphasic CaMKIIα phosphorylation, initially boosting NRF2 defenses, then activating NF-κB-driven inflammation, shifting astrocytes from protective to toxic states. GSDME activation also drives astrocyte-derived exosomes (ADEs) to carry neurotoxic tau, proinflammatory miRNAs, and toxic lipids, propagating toxicity. GSDME deletion in AD mice reduces Aβ burden, restores NF-κB/NRF2 balance, reprograms astrocytes and ADEs to protective states, and rescues cognition. Multi-omics profiling of serum ADEs from AD patients reveals a disease-specific signature with central neurotoxicity and peripheral immune regulation. These findings position GSDME as a promising dual diagnostic and therapeutic target for early AD invention.

## INTRODUCTION

Alzheimer’s disease (AD), the predominant form of dementia, is classically defined by amyloid-β (Aβ) plaques and tau neurofibrillary tangles.^1-4^ However, longitudinal studies reveal that 20-30% of Aβ-positive individuals maintain cognitive stability over decadal follow-ups,^5,6^ underscoring the necessity to identify co-factors that convert Aβ pathology into neurodegeneration. Emerging preclinical evidence reveals a critical temporal pattern, astrocyte reactivity-manifested by pro-inflammatory activation and impaired glutamate uptake precedes detectable Aβ deposition and tau hyperphosphorylation.^7-10^ This early astrocytic dysfunction is further supported by human biomarker studies, where elevated plasma GFAP, a reactivity marker, predicts subsequent tau propagation in presymptomatic stages.^9,11^ Collectively, these findings position astrocyte reactivity as a potential initiator of AD pathogenesis. Given the limited efficacy of Aβ-targeting therapies in late-stage disease,^12,13^ identifying the molecular triggers and spatiotemporal progression of astrocyte reactivity may offer a strategic window for early interventions to halt synaptic degeneration before irreversible damage occurs.^14^

Astrocytic calcium dysregulation emerges as a pivotal early event in AD pathogenesis.^7,10,15^ Under physiological conditions, calcium oscillations in astrocytes maintain synaptic efficiency and neurovascular coupling through regulated gliotransmitter release.^16^ In early AD, however, these cells develop pathological calcium hyperactivity, which occurs prior to Aβ deposition and drives synaptic hyperexcitability.^1,17^ This aberrant signaling disrupts neurovascular coordination by altering prostaglandin release and exacerbates neuroinflammation via uncontrolled cytokine secretion.^18,19^ Crucially, restoring calcium homeostasis reverses cognitive deficits in preclinical models,^20^ while spatial transcriptomics identifies calcium-dependent inflammatory pathways in plaque-associated astrocytes from human AD brains.^21,22^ Despite these advancements, a significant knowledge gap remains regarding the mechanisms by which early calcium signaling anomalies trigger the transition from homeostatic astrocytes to a chronically reactive state.

Emerging evidence underscores the dualistic role of astrocyte reactivity in neurodegeneration, governed by distinct activation states.^19,23^ Early classifications proposed a binary paradigm that neurotoxic (A1) astrocytes induced by microglial inflammatory signals, such as IL-1α and TNF-α, via JAK-STAT/NF-κB pathways, and neuroprotective (A2) astrocytes activated in ischemic contexts.^7,19^ Recently, single-cell and spatial transcriptomics redefine this dichotomy, identifying disease-associated astrocyte (DAA) subpopulations marked by inflammatory gene signatures, notably exhibiting elevated expression of C3, alongside phagocytic dysfunction.^24^ These DAAs engage in bidirectional crosstalk with microglia through cytokines, including IL-6, TGF-β,^25^ and recruit peripheral immune cells via chemokine gradients such as CXCL10,^26^ thereby establishing self-reinforcing neuroinflammatory niches that accelerate synaptic loss. Crucially, spatial mapping reveals DAAs forming peri-plaque "hotspots" that expand with disease progression.^22^ However, the fundamental mechanism by which localized astrocytic inflammatory foci propagate across brain networks remains elusive.

While gasdermin (GSDM)-gated pyroptosis has been reported to engaged in microglial and neuronal dysfunction in neurodegeneration^27^, such as ALS^28^ and MS^29^, however, its role in reactive astrocytes and AD-related pathologies remains largely unknown. Here, we identify a non-canonical GSDME activation pathway that transforms astrocytes into neurodegenerative effectors via calcium-dependent mechanisms. In response to Aβ-activated microglial conditional culture medium, GSDME cleavage targets its N-terminal fragment to mitochondria-associated ER membranes (MAMs), inducing ER calcium efflux and activating the CaMKIIα-NF-κB axis to drive reactive astrocyte-induced neurotoxicity. Notably, astrocytic neurotoxicity can be propagated through astrocyte-derived exosomes (ADEs) enriched with pathogenic cargo. In AD patient brains and 5×FAD mice, GSDME activation was predominantly observed in reactive astrocytes, while genetic GSDME ablation reduces Aβ deposition, restores NF-κB/NRF2 balance, and converts astrocytes and ADEs to neuroprotective phenotypes. Multi-omics of serum ADEs from AD patients reveals a neurotoxic signature, AD-ADEs activate microglia, induce neurotoxic astrocytes, increase neuronal death, and also reduce platelet/neutrophil counts in mice. Our findings collectively identify GSDME as a dual regulator of astrocyte reactivity and ADE neurotoxicity, proposing therapeutic targeting of GSDME alongside serum ADE monitoring for real-time disease tracking.

## RESULTS

### GSDME exhibits spatial and subcluster-specific heterogeneous expression in human and mouse astrocytes

To investigate the potential involvement of GSDM family members in the central CNS, we analyzed publicly available RNA sequencing data from human^30^ and mouse^31^ brain tissues. Notably, GSDME transcript levels were markedly higher in the brain compared to other GSDM family members (Figure S1A and S1B). Further examination of single-cell RNA-seq data^32^ revealed pronounced GSDME expression in the *Slc1a3*^high^ astrocyte subpopulation (Figure S1C), suggesting a cell-type-specific role in astrocytes. Given the established heterogeneity of astrocytes, we next evaluated GSDME expression patterns across astrocyte subtypes using high-resolution single-cell transcriptomic data.^32,33^ Our analysis showed that AST1 and AST4 were the main subtypes with high GSDME expression under resting conditions (Figure S1D). These subtypes are characterized by high *GFAP* and *Slc1a3* expression, consistent with prior whole-brain single-cell studies^32^ (Figure S1C). Intriguingly, GSDME expression correlated strongly with *GFAP*, but less with other astrocytic markers, including *Slc1a3*, *Aldh1l1*, or *Slc1a2* (Figure S1E), underscoring varied GSDME expression among astrocyte subtypes.

Based on the spatial position data concerning the *GFAP*^high^ AST1 and AST4 astrocyte subtypes,^33^ we hypothesized that GSDME expression in astrocytes might similarly favor the hippocampal region. To evaluate this hypothesis, we extracted astrocytes from both the cerebral cortex and the hippocampus for immunoblot analysis. Our results demonstrated that GSDME protein levels were higher in astrocytes isolated from the hippocampus compared to those from the cerebral cortex (Figure S1F and S1G). To verify the astrocyte-specific enrichment of GSDME in the hippocampus, we isolated astrocytes, microglia, and neurons from this region and performed immunoblotting. GSDME was detected exclusively in GFAP-positive astrocyte fractions, while GSDMD was absent (Figure S1H and S1I). Consistently, mouse hippocampal co-staining showed strong GSDME-GFAP colocalization, with no overlap with NeuN or microglial markers (Figure S1J and S1K). Human hippocampal co-staining confirmed GSDME-GFAP colocalization (Figure S1L and S1M), validating single-cell RNA-seq findings.

### Pathological GSDME activation in reactive astrocytes is shared by both AD patients and 5×FAD mouse

To elucidate the pathogenic role of astrocytic GSDME in AD, we conducted an integrative multi-omics analysis of human postmortem brain samples, complemented by functional validation in 5×FAD mouse models. Single-nucleus RNA sequencing comparing AD and other neurodegenerative patients to controls demonstrated a significantly increase in *GSDME* transcript levels (Figure 1A), particularly within reactive astrocyte clusters^34-40^ (Figure S1N-S1Q). As GSDME activation is functionally essential,^41^ we analyzed hippocampal lysates from both AD patients and controls by immunoblotting. The results showed elevated GSDME-NT fragments and increased caspase-3 activation in AD samples, correlating with decreased synaptic expression compared to controls (Figure 1B and 1C).To substantiate the role of astrocytic GSDME activation in AD we conducted co-immunostaining of human tissue sections using C3, a marker for neurotoxic astrocytes, along with GFAP and GSDME-NT. The results indicated a higher proportion of GSDME-NT-positive neurotoxic reactive astrocytes in AD brain samples (Figure 1D and 1E). These observations imply that GSDME activation in reactive astrocytes may contribute to AD pathogenesis.

**Figure 1.**
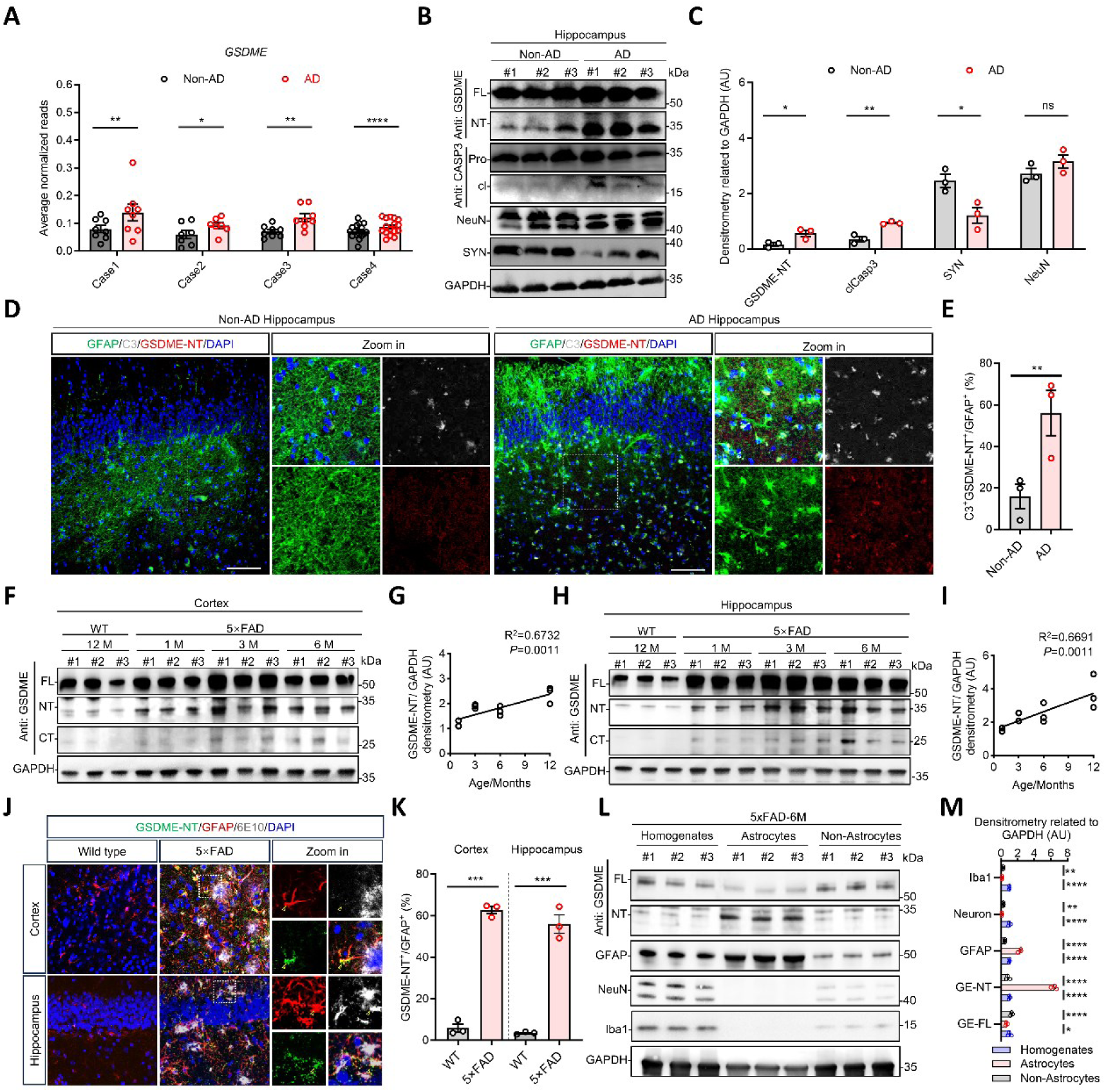
Pathological GSDME activation in reactive astrocytes is preserved across Alzheimer’s disease patients and 5×FAD mouse models. (A) Comparison of GSDME expression in astrocytic clusters between disease (Dg) and healthy (Hg) groups was conducted based on open single-nucleus RNA sequencing (snRNA-seq) datasets of AD (Case 1^34^, *n*_Hg_ = 8, *n*_Dg_ = 8; Case 2^35^, *n*_Hg_ = 7, *n*_Dg_ = 27; Case 3^36^, *n*_Hg_ = 8, *n*_Dg_ = 8; Case 4^38^, *n*_Hg_ = 16, *n*_Dg_ = 16). (B and C) Immunoblot analysis of the activation of GSDME, caspase-3, as well as the expression of NeuN and Synaptophysin (SYN) in hippocampus tissue lysates collected from AD patients and healthy controls (hippocampus, *n*_Hg_ = 3, *n*_Dg_ = 3), measured by Image J based on gray scale. (D and E) Representative IF images of hippocampus region from AD patients and healthy controls (hippocampus, *n*_Hg_ = 3, *n*_Dg_ = 3) co-stained with anti-GSDME-NT, anti-GFAP and anti-C3 antibody. Scale bars, 100 μm. Percentage of GSDME-NT and C3-double positive cells within GFAP-positive cell populations was quantified using Image J software. Each point means two sections. (F-I) Immunoblot analysis of GSDME activation in cortex (F and G) and hippocampus (G and H) tissue lysates collected from 5×FAD and wild type (WT) mice. Regression analysis of GSDME-NT expression dynamics during aging in 5×FAD mice. GSDME-NT levels were quantified by ImageJ (*n* = 3 for each group). (J and K) Representative IF images of cortex and hippocampus region from 5×FAD (*n* = 3) and WT mice (*n* = 3) co-stained with anti-GSDME-NT, anti-GFAP and anti-6E10 antibody (J). Scale bars, 30 μm. Percentage of GSDME-NT within GFAP-positive cell populations (K) was quantified by Image J. (L-M) Immunoblot analysis of GSDME-NT, GFAP, as well as the expression of NeuN and Iba1 in cortex tissue homogenates, astrocyte-enriched fractions, and non-astrocytic fractions isolated from 6-month-old 5×FAD mice, measured by Image J (*n* = 3 for each fraction). Statistical significance determined using a two-tailed Student’s *t* test (C, E, G, I, K and M), or two-sided Wilcoxon signed-rank test (A). All values are represented as mean ± SEM, \**p* < 0.05; \*\**p* < 0.01; \*\*\**p* < 0.001; \*\*\*\**p* < 0.0001. See also Figure S1 and Table S1.

To further validate the *in vivo* significance of GSDME activation in AD pathogenesis, we examined cortical and hippocampal tissues from 5×FAD mice at various aging timepoints. Immunoblotting revealed a significant linear correlation between the intensity of the GSDME cleavage fragment band and mouse age, suggesting progressive GSDME activation in parallel with AD pathology (Figure 1F-1I). Immunofluorescence (IF) analysis in 6-month-old 5×FAD mice showed an increase in GSDME-NT-positive astrocytes surrounding Aβ plaques, compared to age-matched controls, with pronounced colocalization between GSDME-NT and C3 (Figure 1J and 1K). Notably, adult mouse brain cell fractionation assays confirmed that GSDME cleavage products were predominantly localized within the astrocytic compartment (Figure 1L and 1M). Collectively, these findings demonstrate that pathological activation of GSDME in reactive astrocytes is a common feature observed in both AD patients and the 5×FAD mouse model.

### Non-pyroptotic GSDME activation is required for reactive astrocyte-mediated neurotoxicity

Building upon the observed pathological correlation between GSDME activation and AD progression, we subsequently explored its functional role in Aβ-induced glial crosstalk, a recognized contributor to neurotoxicity in AD pathogenesis.^15,42^ To emulate disease-relevant interactions between microglia and astrocytes, we established an *in vitro* model system. In this system, BV2 microglial cells were primed with pathogenic Aβ_1-42_ oligomers to produce activated microglia-conditioned medium (AMCM), and primary astrocytes were subsequently exposed to this AMCM to observe phenotypic transformation. Temporal transcriptomic analysis demonstrated that prolonged exposure to AMCM induced a progressive transformation in astrocytes, characterized by a stable neurotoxic transcriptional signature at 24, 48, 72 and 96 h post-stimulation (hps) compared to microglia-conditioned medium (MCM) with PBS treatment. This neurotoxic shift was characterized by three distinct molecular features: (1) Upregulation of neurotoxic mediators, such as *C3*, and *Srgn*, consistent with their known roles in reactive astrogliosis and neuroinflammatory responses;^19^ (2) Dysregulation of protective pathways, evidenced by significant downregulation of *S100A10*, *Sphk1* and *Emp1*, thereby compromising their neuroprotective functions; and (3) Deficits in neurotrophic support, as indicated by reduced levels of *Fgf1*, *Shh* and *Gpc4*, critically impairing synaptic maintenance and neuronal survival^43^ (Figure S2A and S2B). These molecular changes collectively establish AMCM-induced astrocytic transformation as a key mediator of AD-related neurodegeneration, with concurrent activation of detrimental pathways and suppression of protective mechanisms.

To elucidate the functional role of GSDME activation in reactive astrocytes, we investigated whether the caspase-3-GSDME signaling pathway (Figure 2A), which executes pyroptosis through GSDME cleavage,^41^ was operational in AMCM-treated astrocytes. Temporal analysis following AMCM treatment demonstrated sustained caspase-3 activation, leading to robust GSDME cleavage and production of GSDME-NT fragments (Figure 2B and S2C). Contrary to the anticipated outcomes associated with pyroptotic cell death, neither LDH release assays (Figure 2C and S2D) nor PI staining (Figure 2D and S2E) detected compromised membrane integrity in AMCM-treated groups. These findings contrast with the significant cell death increases observed in GSDME-NT-EGFP overexpression controls. To further validate these observations, we performed IF staining (Figure 2E, 2F and S2F-2H) and scanning electron microscopy (SEM) on AMCM-treated astrocytes (Figure 2G and 2H). Quantitative analysis revealed a marked increase in GSDME-NT-positive cells under AMCM conditions, while the cellular morphology remained indistinguishable from controls, exhibiting no blebbing and intact membranes, and demonstrated significantly lower PI-positive cell counts compared to GSDME-NT-EGFP controls (Figure 2E-2H). These convergent data imply that while AMCM activates GSDME cleavage in neurotoxic reactive astrocytes, this activation does not induce classical pyroptotic features such as membrane pore formation or cell blebbing typically associated with pyroptotic cell death.

**Figure 2.**
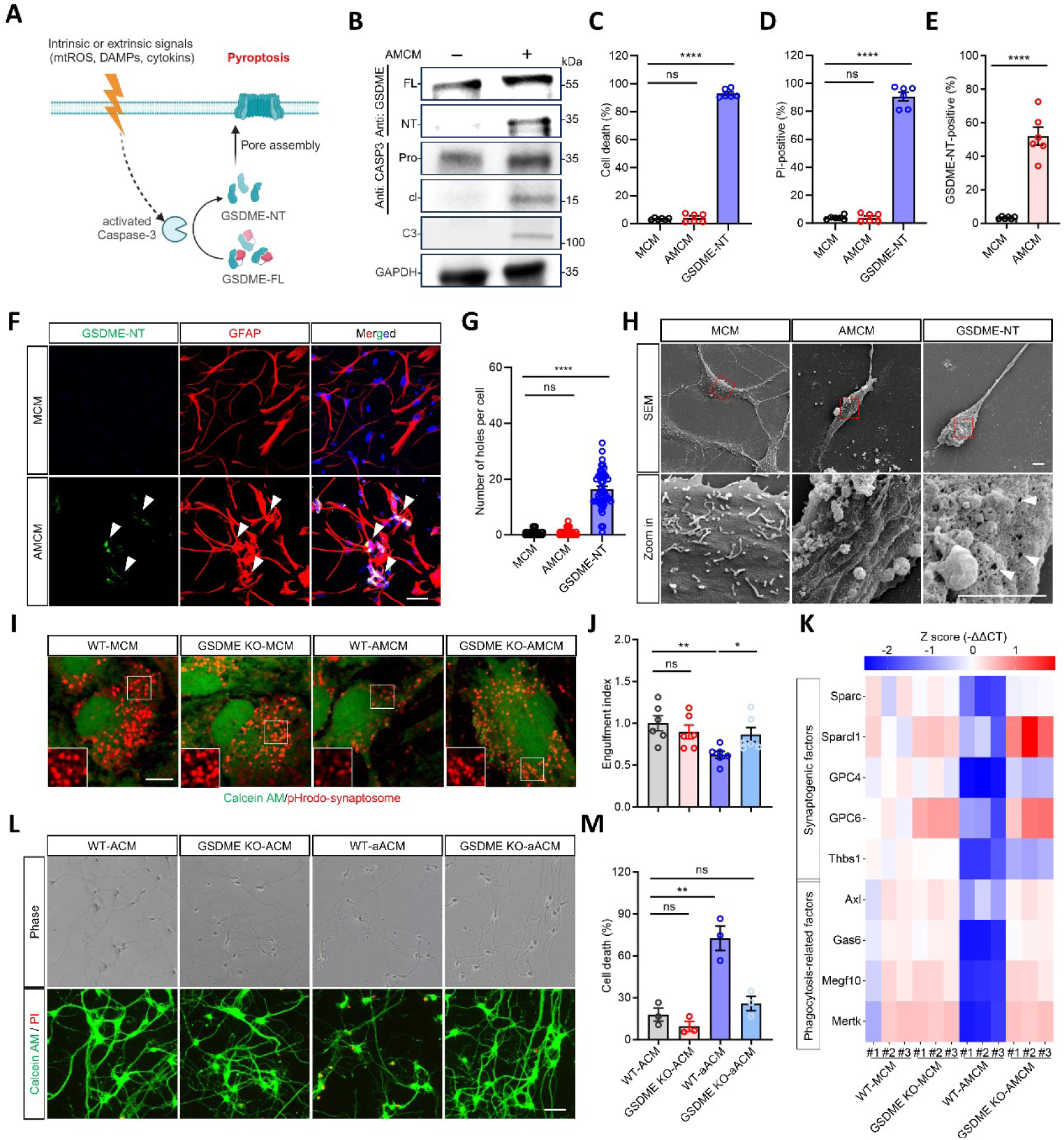
GSDME activation enhances reactive astrocyte-mediated neurotoxicity independent of pyroptosis. (A) Schematic of GSDME cleavage mediated by activated caspase-3 to induce cell membrane rupture and pyroptosis upon intrinsic or extrinsic stimulus (biorender.com). (B) Immunoblotting of caspase-3, GSDME activation and C3 secretion of primary astrocytes upon stimulation by MCM or AMCM with Aβ_42_ oligomers plus LPS at 48 h. (C and D) Percentage of primary astrocytic cell death was quantified by LDH release (C) or PI staining (D) following the astrocytes treated with AMCM or by overexpression of GSDME-NT-EGFP (*n* = 6 replicates per group). (E and F) Percentage of GSDME-NT-positive astrocytes in (C) was calculated with Image J (E) and representative IF images were shown in (F). GSDME-NT (anti GSDME-NT; green) in primary astrocytes (anti-GFAP; red) 48 h post MCM or AMCM treatment. Scale bars, 20 μm. Arrows indicate the GSDME-NT^+^ astrocytes (*n* = 6 replicates per group). (G and H) Quantification of holes (G) on the primary astrocytic cell membrane via SEM at 48 h post MCM or AMCM treatment. Astrocytes overexpressing GSDME-NT were used as positive controls. Representative SEM images are shown in (H). Scale bars, 3 μm. Arrows indicate the holes. (I and J) Representative images of WT or GSDME KO (knock-out) primary astrocytes treated with MCM or AMCM as indicated, showing the engulfment of pHrodo conjugated synaptosomes (I) and the extent of engulfing pHrodo-conjugated synaptosomes was calculated with Image J (J). The inserts show the engulfed particles (red). Scale bars, 10 μm. (K) Relative mRNA expression of phagocytosis-related proteins (*Axl*, *Gas6*, *Megf10*, *Mertk*), synaptogenic factors (*Sparc*, *Sparcl1*, *GPC4*, *GPC6*, *Thbs1*), quantified by qRT-PCR (*n* = 3 replicates per group). (L and M) Representative live images of primary cortical neuronal death induced by ACM, analyzed with the Calcein AM/PI staining (L) and the ratio of neuronal death was calculated by Image J (M) (*n* = 3 replicates per group). Scale bars, 20 *μ*m. Statistical significance determined using a two-tailed Student’ s *t* test. All values are represented as mean ± SEM, \**p* < 0.05; \*\**p* < 0.01; \*\*\*\**p* < 0.0001. See also Figure S2.

Recent studies indicate that neurotoxic reactive astrocytes obtain aberrant functions, including impaired phagocytosis and disrupted induction of synaptogenesis.^19^ To assess the role of GSDME in these abnormalities, we stimulated both wild-type and GSDME KO astrocytes with MCM or AMCM for 24 h, evaluating phagocytic activity by quantifying synaptosome uptake. The results demonstrated that AMCM-treated GSDME KO astrocytes exhibited increased phagocytosis of pHrodo-labeled synaptosomes compared to AMCM-treated WT astrocytes, aligning with MCM-treated WT astrocytes (Figure 2I and 2J). This corresponds with upregulated mRNA expression of phagocytic receptors,^19^ *Megf10, Mertk, Axl,* along with the bridging molecule *Gas6* in AMCM-exposed GSDME KO astrocytes compared to AMCM-treated WT astrocytes (Figure 2K). Additionally, genes encoding synaptogenic factors, *Thbs1, Gpc6, Sparcl1* were upregulated in AMCM-treated GSDME KO astrocytes versus WT (Figure 2K). This suggests that the knockout of GSDME, to some extent, alleviates the abnormalities in astrocytic function following AMCM treatment. To assess neurotoxicity, we administered astrocyte-conditioned medium (ACM) harvested from MCM-treated WT, MCM-treated GSDME KO, AMCM-treated WT, and AMCM-treated GSDME KO astrocytes to cultures of mouse primary cortical neurons. Our results indicated that, in comparison to the control group exposed to ACM derived from AMCM-treated WT astrocytes, ACM from AMCM-treated GSDME KO astrocytes significantly reduced neuronal deaths (Figure 2L and 2M). These findings suggest that GSDME deficiency have the potential to attenuate neuronal death caused by neurotoxic reactive astrocytes, thereby highlighting a possible therapeutic target for neuroprotective strategies.

### MAM-localized GSDME-NT drives ER-to-cytosol Ca²⁺ efflux to trigger neurotoxic reactive astrogliosis

Given that GSDME was activated in reactive astrocytes without inducing pyroptosis, we sought to identify the specific intracellular membranes targeted by the released GSDME-NT. To determine the distribution of GSDME-NT across the intracellular membranes of reactive astrocytes, we employed a standard protocol^44^ for isolating fractions of mitochondria, endoplasmic reticulum (ER), and mitochondria-associated ER membranes (MAMs), constituting the primary intracellular membrane systems in mammalian cells (Figure 3A). To assess the quality of subcellular fractionation, immunoblotting was utilized to analyze marker proteins specific to different cellular compartments. The results revealed that VDAC1, a mitochondrial marker, was predominantly detected in the pure mitochondria fraction and was absent from the cytosol fraction. Likewise, calnexin, acting as a marker for both ER and MAMs, exhibited significant levels in both ER and MAM fractions. Consistent with these findings, ACSL4, previously identified as an MAM-resident protein,^45^ was enriched in the MAM fraction. Conversely, β-actin, a cytosolic marker, was not detected in mitochondria, MAMs, or ER fractions (Figure 3A). Collectively, these findings suggest a high quality of subcellular fractionation harvest.

**Figure 3.**
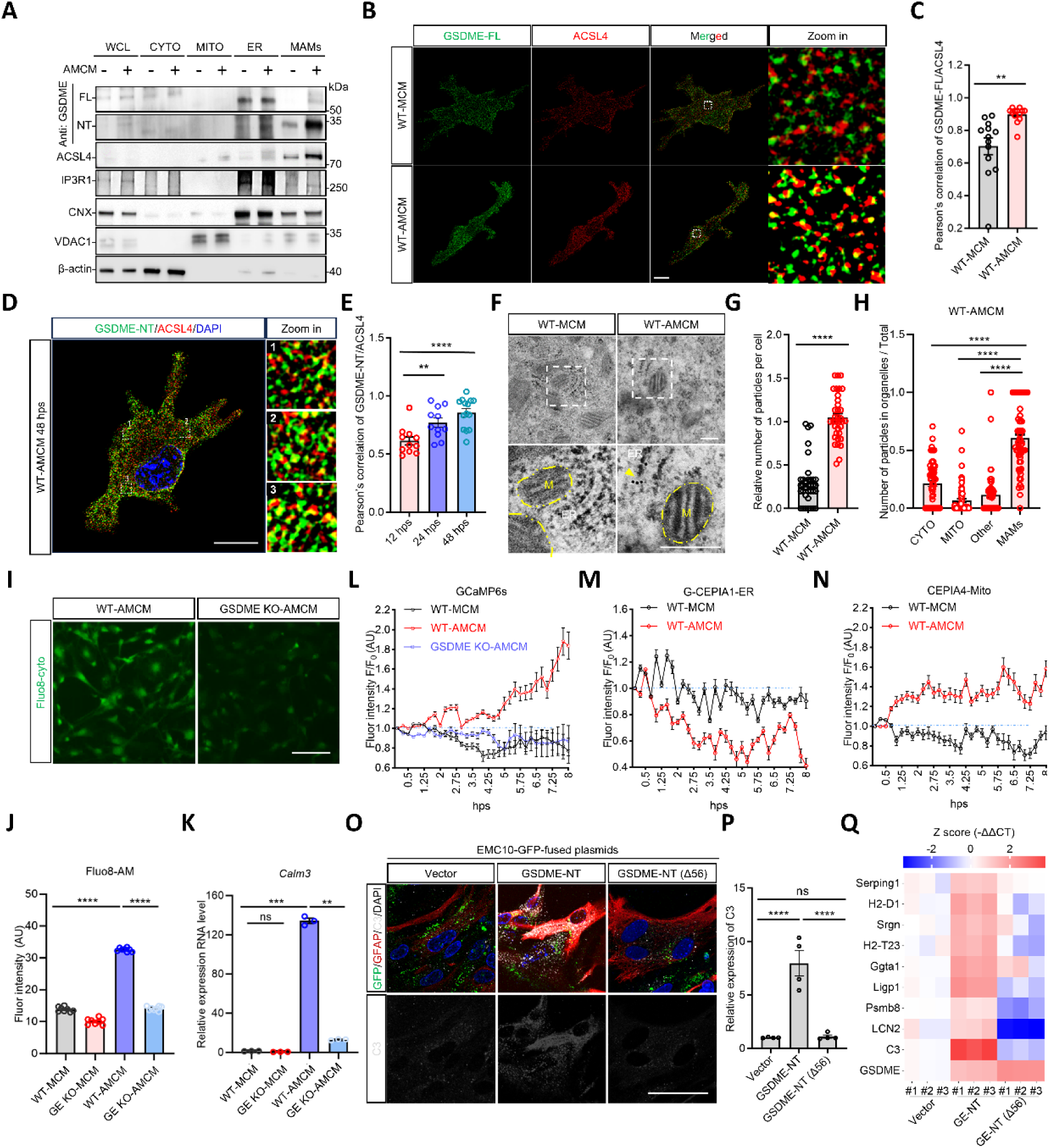
MAM-Localized GSDME-NT induces neurotoxic reactive astrogliosis via ER-to-Cytosol Ca²⁺ Efflux. (A) Immunoblotting of subcellular fractionation from WT-MCM or WT-AMCM astrocytes demonstrates the subcellular distribution of GSDME-FL and GSDME-NT. Each fraction was confirmed by immunoblotting with antibodies to ACSL4, IP3R1 (ER-resident protein), CNX, VDAC1 and β-actin. WCL, whole cell lysates. CYTO, cytosol. MITO, mitochondria. ER, endoplasmic reticulum. MAMs, mitochondria associated endoplasmic reticulum membranes. (B and C) SIM images stained for GSDME-FL and ACSL-4 (B) at 48 hps. Quantification of Pearson’ s correlation between GSDME-FL and ACSL-4 (C). Enlarged images of the boxed area are shown (*n* = 2 replicates, at least 8 cells were calculated each group per replicate). Scale bars, 5 μm. (D and E) SIM images stained for GSDME-NT and ACSL-4 at 48 hps (D). Quantification of Pearson’s correlation between GSDME-NT and ACSL-4 (E). Enlarged images of the boxed area are shown (*n* = 2 replicates, at least 8 cells were calculated each group per replicate). Scale bars, 10 μm. (F-H) TEM images stained with GSDME-NT-immunogold (F). Yellow dashed lines outline mitochondria, arrows indicate the GSDME-NT-immunogold. Scale bars, 200 nm. Quantification of gold particles in WT-MCM astrocytes versus WT-AMCM astrocytes was shown in (G). The relative subcellular distribution of gold particles in WT-AMCM astrocytes, including cytosol, mitochondria, MAMs were further quantified in (H) (*n* = 3 replicates, 10-20 cells each group per replicate). (I-K) Representative IF image stained with Fluo8-AM was shown in (I). Quantification of Fluo8 AM fluorescence intensity to indicate the cytosolic Ca^2+^ was shown in (J) (*n* = 8 replicates per group). Scale bars, 50 μm. Relative mRNA expression of endogenous cytosolic Ca^2+^ sensor, Calm3, quantified by qRT-PCR was shown in (K) (*n* = 3 replicates per group). (L-N) Quantification of relative fluorescence intensity over time in WT-MCM, WT-AMCM or GSDME KO-AMCM astrocytes by the genetically encoded Ca^2+^ indicators, GCaMP6s (L) for the cytosol, CEP1AE-ER (M) for the ER, and CEPIA4-Mito (N) for the mitochondria. (O-Q) Representative IF image stained with C3 (grey) in primary mouse astrocytes (anti-GFAP; red) was shown in (O), quantified by qRT-PCR was shown in (P) (*n* = 4 replicates per group). Relative mRNA expression of reactive astrocyte marker (*Serping1*, *H2-D1*, *Srgn*, *H2-T23*, *Ggta1*, *Ligp1*, *Psmb8*, *LCN2*), quantified by qRT-PCR (*n* = 3 replicates per group). Statistical significance determined using a two-tailed Student’ s *t* test. All values are represented as mean ± SEM, \*\**p* < 0.01; \*\*\**p* < 0.001; \*\*\*\**p* < 0.0001. See also Figure S3 and Movie S1-S3.

Upon administering AMCM, we observed a notable elevation in GSDME-NT fragments in the whole cell lysates of WT-AMCM astrocytes compared to MCM-treated controls (Figure 3A). This finding implies the activation of GSDME in reactive astrocytes. Further delving the activation of GSDME in various subcellular fractions, we discovered a significant enrichment of GSDME-NT fragments specifically in the MAM fraction, as opposed to the mitochondria fraction (Figure 3A). IF staining followed by structured illumination microscopy (SIM) imaging revealing a significant elevation in the colocalization of GSDME-FL with ACSL4 in WT-AMCM astrocytes compared to WT-MCM astrocytes (Figure 3B and 3C). Additional SIM imaging showcased a high degree of colocalization between GSDME-NT and ACSL4 (Figure 3D and 3E) and hinted at the targeting of GSDME-NT to MAMs. Furthermore, transmission electron microscopy (TEM) observations confirmed frequent GSDME-NT labeling at ER-mitochondria contact sites (Figure 3F). Stereological analysis further revealed the distinct enrichment of GSDME-NT at MAMs (Figure 3G and 3H). These results suggest that GSDME, activated by AMCM, is translocated to MAMs in reactive astrocytes.

Ca^2+^ transfer is a primary function attributed to ER-mitochondrial contact sites.^46^ Given that GSDME-NT targets and forms pores on the membrane, we investigated whether the translocation of GSDME-NT to MAMs facilitates Ca^2+^ transfer. To test this hypothesis, we utilized Fluo 8-AM, a fluorescent cytoplasmic Ca^2+^ indicator, to monitor changes in cytoplasmic Ca^2+^ concentration in reactive astrocytes. Our findings reveal that incubation with AMCM significantly increases cytoplasmic Ca^2+^ concentration in these cells (Figure 3I and 3J). However, this elevation was markedly impeded in the absence of GSDME (Figure 3I and 3J). Additionally, we quantified the mRNA expression of *Calm3*, an inherent calcium-binding protein capable of sensing cytosolic Ca^2+^ levels,^47^ as an indicator of cytoplasmic Ca^2+^ concentration changes in astrocytes. Consistently, we noted a reduction in *Calm3* transcription in GSDME KO-AMCM astrocytes compared to their WT-AMCM counterparts (Figure 3K). These results suggest that GSDME-NT targeting MAMs contributes to the elevation of cytoplasmic Ca^2+^ concentration in reactive astrocytes.

Next, we expressed genetically encoded Ca^2+^ indicators specifically targeting cytosol (GCaMP6s), ER (CEPIA1-ER), and mitochondria (CEPIA4-Mito) to monitor Ca^2+^ dynamics^48,49^ (Figure S3A-S3E). Notably, the depletion of extracellular Ca^2+^ did not change the intracellular Ca^2+^ elevation through GCaMP6s fluorescence intensity observation in astrocytes treated with AMCM (Figure 3L). This suggests that the cytosolic Ca^2+^ rise primarily originates from GSDME pores located at the endosomal membrane, rather than the plasma membrane. Meanwhile, the fluorescence intensity enhancement of GCaMP6s in WT-AMCM astrocytes was suppressed in the absence of GSDME (Figure 3L). These results reinforce the hypothesis that GSDME-NT targeting MAMs contributes to the cytoplasmic Ca^2+^ concentration elevation. To pinpoint the source of this cytoplasmic Ca^2+^ surge, we dissected Ca^2+^ dynamics within mitochondria and ER, the principal Ca^2+^ reservoirs in astrocytes. The analysis reveals that AMCM triggers a decrease in ER Ca^2+^ dynamics, as evidenced by CEPIA1-ER fluorescence (Figure 3M), while simultaneously inducing an increase in mitochondrial Ca^2+^ dynamics, demonstrated by CEPIA4-Mito fluorescence (Figure 3N), in reactive astrocytes compared to WT-MCM astrocytes. Therefore, we believe that GSDME-NT localizes to MAMs, facilitating the release of Ca^2+^ from ER into cytosol.

To mechanistically interrogate the causal linkage between GSDME-NT-driven ER calcium mobilization and neurotoxic astrocyte transformation, we designed two molecular probes, namely full-length EMC10-GSDME-NT-EGFP incorporating the ER-localizing EMC10 domain^50^ and its N-terminal 56-amino acid^41^ truncated counterpart EMC10-GSDME-NT(Δ56)-EGFP. Lentiviral delivery of EMC10-GSDME-NT-EGFP in primary astrocytes preserved viability equivalent to EMC10-EGFP controls, consistent with prior observations of pyroptosis-independent GSDME activation (Figure S3F). IF quantification demonstrated a notable elevation in C3 expression in cells expressing EMC10-GSDME-NT, a finding further substantiated by qPCR, which indicated a coordinated transcriptional upregulation of genes specific to neurotoxicity (Figure 3O-3P). In contrast, the Δ56 truncation mutant not only inhibited C3 induction but also repressed the expression of neurotoxic-specific genes (Figure 3O-3P).These evidences collectively suggest that MAM-anchored GSDME-NT mediates compartment-specific calcium shutting from ER to cytosol, which functions as a molecular switch initiating the neurotoxic astrogliosis program.

### Phase-Specific CaMKⅡα phosphorylation by GSDME governs the neuroprotective-to-neurotoxic transition

To elucidate how GSDME-NT induces MAM-dependent calcium dysregulation and drives astrocytes to a neurotoxic state, we combined transcriptomic profiling with functional validation. Unexpected, bulk RNA sequencing demonstrated that GSDME knockout suppressed transcription of neurotoxic genes such as *C3* and *Lcn2*, while upregulating protective markers including *S100A10*, *Emp1*, and *Sphk1* (Figure 4A). These data indicate that GSDME is essential for reprogramming astrocytes toward a neuroprotective phenotype. KEGG pathway enrichment analysis identified calcium signaling as the most significantly affected pathway (Figure 4B), aligning with GSDME’s role in modulating calcium dynamics at the ER-mitochondria interface. Further differential expression analysis within the calcium signaling category highlighted *Camk2a* (the gene encoding CaMKIIα) as a central regulator of the GSDME-driven neurotoxic transition (Figure 4C). Functionally validation via siRNA-mediated knockdown of *Cam*k2a demonstrated that suppression of CaMKIIα abrogates the induction of protective gene (Figure 4D), confirming CaMKIIα as a key effector in the GSDME-calcium signaling axis.

**Figure 4.**
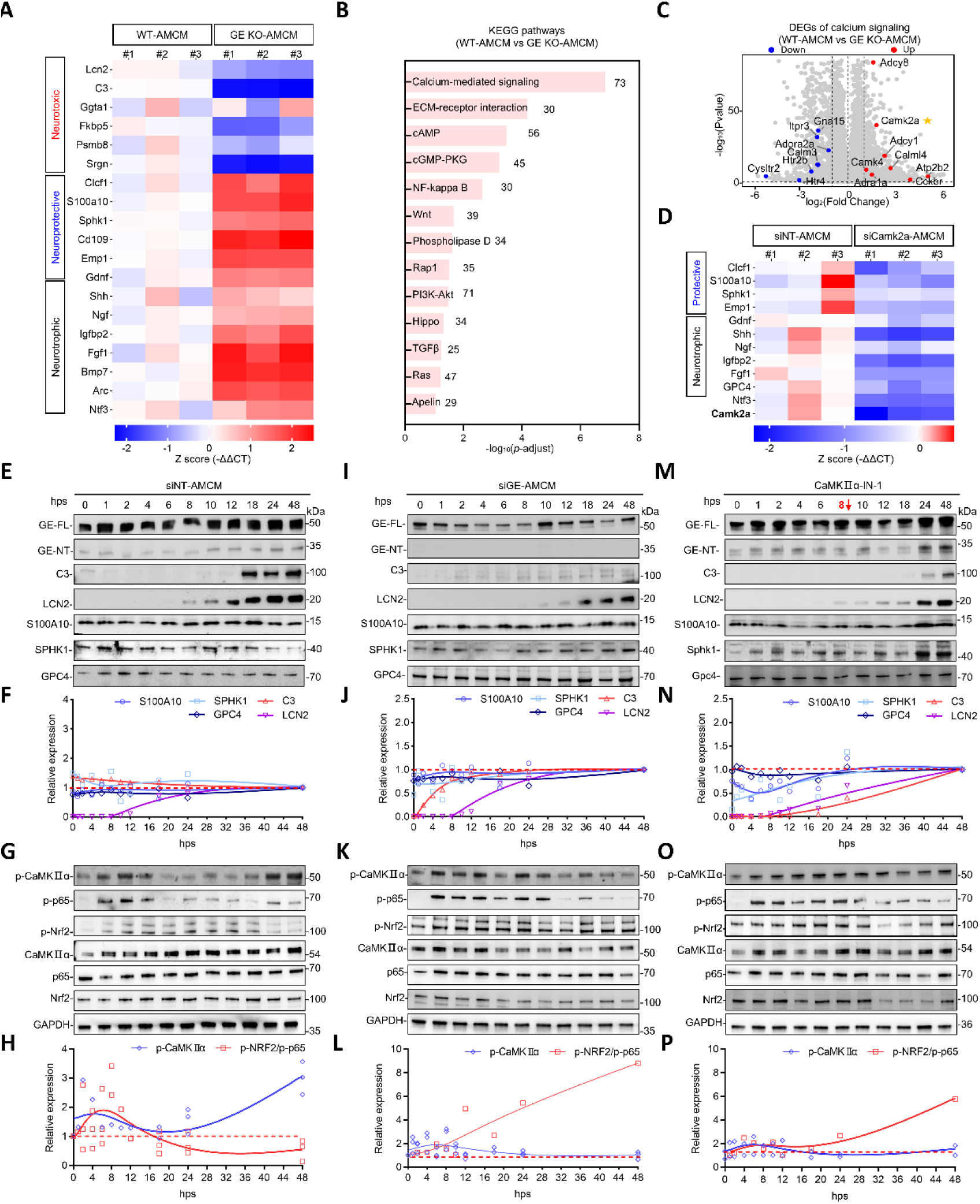
Phase-Specific CaMKIIα phosphorylation orchestrates GSDME-driven neuroprotection-to-neurotoxicity switch. (A) Hierarchical clustering heatmap of RNA-seq profiles illustrating distinct expression patterns of neurotoxic, neuroprotective, and neurotrophic genes in WT versus GSDME-knockout (GE KO) primary astrocytes following AMCM stimulation at 24 h. (B) KEGG pathway enrichment analysis of differentially expressed genes (DEGs) identified between GSDME KO and WT astrocytes. Pathways are ranked by enrichment scores (−log_10_(*p*-value)). (C) Volcano plot of calcium-mediated signaling genes in GSDME KO vs WT astrocytes. Significantly upregulated (red, |log_2_FC| > 1, FDR < 0.05) and downregulated (blue) genes are annotated. *Camk2α* (asterisk) is highlighted as a top DEG. (D) Heatmap analysis of neuroprotective and neurotrophic gene expression by qPCR in astrocytes transfected with CaMKIIα-targeting siRNA (siCamk2α) or non-targeting scrambled siRNA (siNC), followed by 24-h AMCM stimulation. Data are normalized to *Gapdh* and presented as fold change relative to siNC. (E-G) Time-dependent immunoblot analysis of GSDME-FL and GSDME-NT, neurotoxic (C3, LCN2), neuroprotective (S100A10, SPHK1), and neurotrophic (GPC4) markers in astrocytes transfected with non-targeting scrambled siRNA (siNT) following AMCM treatment for 48 h (E). Quantification of grayscale values (ImageJ) and temporal trends (GraphPad Prism fit spline analysis) are shown in (F). Phosphorylation kinetics (G) of CaMKIIα, p65, and NRF2 in (E). Quantification of grayscale values (ImageJ) and temporal trends (GraphPad Prism fit spline analysis) are shown in (H). (I-L) Time-dependent immunoblot of GSDME-FL/NT, neurotoxic/neuroprotective markers, and GPC4 (I and J) and phosphorylation kinetics (L and M) in astrocytes transfected with siRNA targeted GSDME (siGE) under identical conditions in (E). Quantification and trend fitting are shown in (J and L). (M-P) Time-dependent immunoblot of GSDME-FL/NT, neurotoxic/neuroprotective markers, and GPC4 (M and N) and phosphorylation kinetics (O and P) in astrocytes treated with CaMKⅡα-IN-1at 8 hps with AMCM. Quantification and trend fitting are shown in (O and Q). Statistical significance determined using a two-tailed Student’s *t* test. All values are represented as mean ± SEM with at least 3 biological replicates. Each point represents one replicate. See also Figure S4.

Time-resolved immunoblot analysis revealed that astrocytes adopt a neuroprotective phenotype within 12 hps, as indicated by increased expression of S100A10, SPHK1, and GPC4, along with suppressed levels of C3 and LCN2 (Figure 4E and 4F). Strikingly, after 12 hps, astrocytes transition to a neurotoxic state: C3 and LCN2 are robustly upregulated, whereas protective markers S100A10, GPC4, and SPHK1 decline (Figure 4E and 4F). This biphasic pattern demonstrates that induction of a neuroprotective phase precedes toxic conversion, consistent with previous observation.^51^ Concurrently, CaMKⅡα phosphorylation (p-CaMKⅡα) exhibits a dynamic biphasic profile (Figure 4G and 4H), implicating posttranslational regulation of astrocyte state. Detailed analysis of NF-κB and NRF2 signaling revealed that, during the early phase (0-12 hps), phosphorylated NRF2 (p-NRF2) predominates while phosphorylated NF-κB subunit p65 (p-p65) remains low. In contrast, the late phase (12-48 hps) shows diminished p-NRF2 and increased p-p65 (Figure 4G and 4H). Importantly, the ratio of p-NRF2 to p-p65 correlates positively with p-CaMKIIα activity (Figure 4H), suggesting that CaMKⅡα dynamically balances NF-κB and NRF2 signaling to dictate astrocyte fate.

To validate the spatiotemporal regulation of CaMKⅡα phosphorylation by GSDME, we conducted time-resolved immunoblotting in astrocytes with GSDME silenced. During the early phase (0-12 hps), GSDME silencing did not significantly alter protective marker expression, likely reflecting minimal GSDME activation at this stage (Figure 4I and 4J). However, in the late phase (12-48 hps), GSDME knockdown prevented induction of neurotoxic markers and maintained protective marker expression, effectively blocking the transition to a neurotoxic phenotype (Figure 4I and 4J). Correspondingly, GSDME silencing attenuated late-phase CaMKⅡα phosphorylation and preserved the p-NRF2/p-p65 ratio (Figure 4K and 4L). These findings suggest that early-phase CaMKⅡα phosphorylation is required for neuroprotection, whereas excessive phosphorylation at later stages disrupts the NF-κB/NRF2 balance and promotes toxicity.

Pharmacological modulation corroborated the phase-specific requirement for CaMKⅡα in astrocyte fate transition. Continuous inhibition at the beginning of stimulus with CaMKIIα-IN-1^52^ failed to prevent the neurotoxic conversion (Data not shown). In contrast, a sequential treatment recapitulated the protective effects observed with GSDME ablation. This regimen elevated S100A10, GPC4, and SPHK1 and reduced C3 and LCN2 levels (Figure 4M and 4N). Mechanistically, early-phase activation of CaMKⅡα enhanced the p-NRF2/p-p65 ratio, while subsequent inhibition prevented late-phase CaMKIIα hyperphosphorylation and suppressed neurotoxic marker induction (Figure 4O and 4P). Together, these findings underscore the necessity of precise temporal regulation of CaMKⅡα activity: early activation potentiates NRF2-mediated neuroprotection, whereas unchecked hyperphosphorylation in later phases shifts the balance toward NF-κB-driven neurotoxicity. Overall, GSDME functions a molecular switch that dynamically modulates CaMKIIα phosphorylation to maintain NF-κB/NRF2 signaling equilibrium and govern the transition between neuroprotective and neurotoxic states in astrocytes.

### GSDME reprograms astrocyte-derived exosomes to propagate neurotoxic signals

Given that GSDME activation elevates cytosolic Ca²⁺ levels in reactive astrocytes, we examined whether altered calcium signaling regulates exosome production in these cells. Using a standardized exosome isolation protocol, we obtained astrocyte-derived exosomes (ADEs) from ACM and aACM, respectively (Figure 5A-5C). TEM imaging confirmed intact vesicular structures (Figure S5A), while nanoparticle tracking analysis (NTA) revealed size distributions between 30-200 nm, consistent with canonical exosome dimensions (Figure S5B). Notably, quantitative comparison showed equivalent average diameters between ACM-ADEs (ADEs) and aACM-ADEs (aADEs) (Figure S5C), while the particle counts of aADEs were much higher than those of ADEs (Figure S5D) Functional validation through GSDME knockout revealed markedly reduced aADEs production in GSDME-deficient astrocytes cultured in MCM (Figure 5B), suggesting GSDME as a critical regulator of ADEs biogenesis in reactive astrocytes.

**Figure 5.**
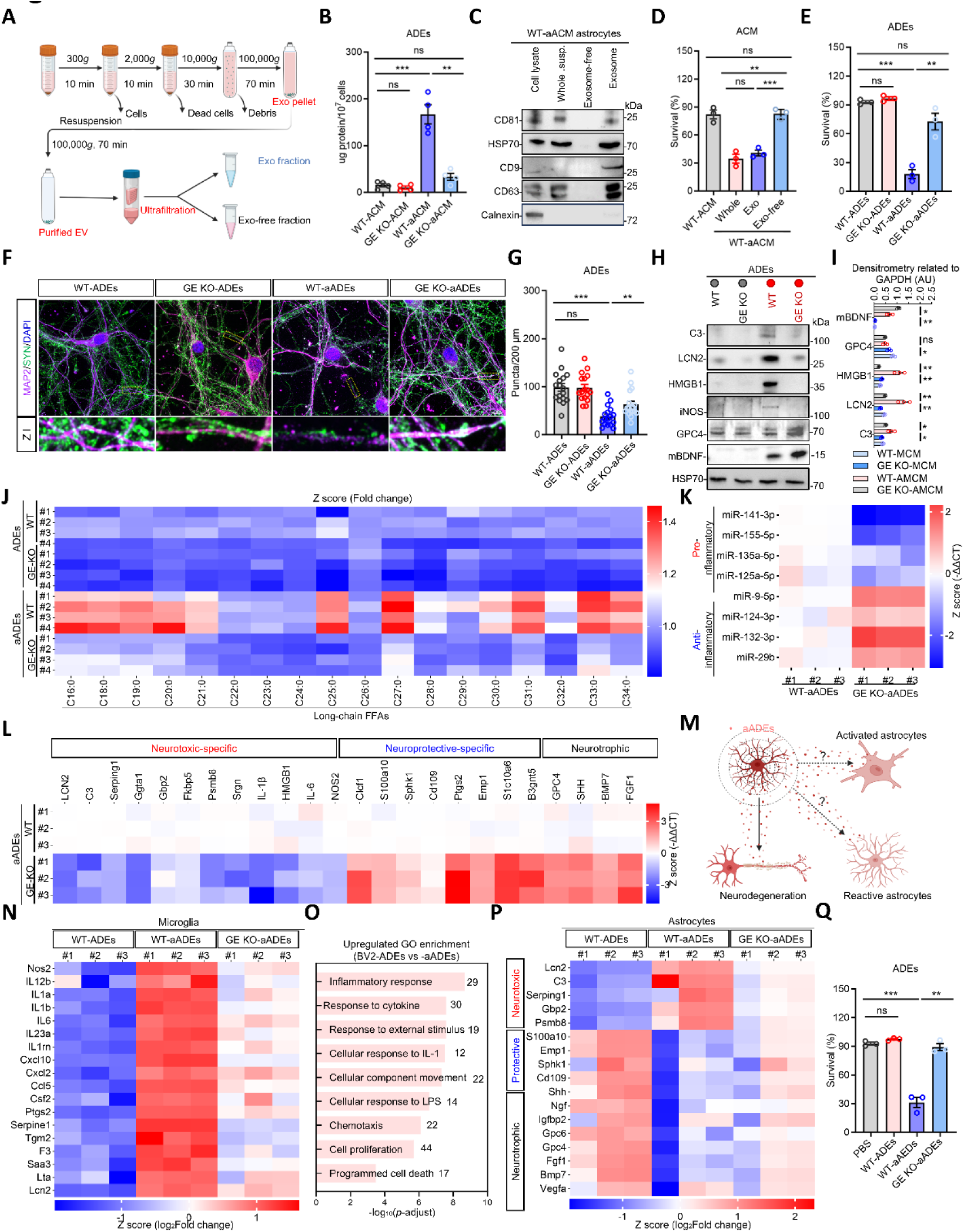
GSDME regulates the production of astrocytic neurotoxic exosomes. (A-C) Diagram of exosome (Exo) or Exo-free fraction isolation based on a standard protocol (A). Quantification of exosomes isolated from WT-ACM, GSDME KO-ACM, WT-aACM and GSDME KO-aACM astrocytes by Bradford protein assay (B) (*n* = 4 replicates per group). Each fraction was confirmed by immunoblotting with antibodies CD81, HSP70, CD9 and CD63, Calnexin (C). (D) Quantification of the neuronal cell survival rate of mouse cortical neurons 24 h after the treatment with WT-ACM, aACM (Whole), aACM-isolated exosomes (Exo) and Exo-free aACM (Exo-free) (*n* = 3 replicates per group). (E) Quantification of the neuronal cell survival rate of mouse cortical neurons 24 h after the treatment of ACM-isolated exosomes from WT-ACM, GSDME KO-ACM, WT-aACM and GSDME KO-aACM astrocytes supernatant (*n* = 3 replicates per group). (F and G) Representative IF images of neurons, as treated in (E), stained for both MAP2 and SYN (F). Enlarged images of the boxed area are shown (*n* = 2 replicates, at least 10 cells were calculated each group per replicate). The number of puncta per 200 μm as in (D) is calculated(G). Scale bars, 10 μm. (H and I) Immunoblotting of equal amounts of exosomes with antibodies, C3, LCN2, HMGB1, iNOS, GPC4 and BDNF, controlled by HSP70 (H), measured by ImageJ based on gray scale (I). (J) Quantification of saturated lipids in equal amounts of exosomes, as treated in (E), by liquid chromatography-mass spectrometry (LC-MS). FFA, free fatty acid (*n* = 4 replicates per group). (K) Quantification of pro- and anti-inflammatory miRNAs in exosomes, as treated in (E), by RT-qPCR (*n* = 3 replicates per group). (L) Relative expression of neurotoxic, neuroprotective and neurotrophic genes in exosomes, as treated in (E), by RT-qPCR (*n* = 3 replicates per group). (M) Schematic model of ADE-mediated neuroinflammatory propagation across neuronal-glial networks. (N and O) Heatmap depicting relative expression of pro-inflammatory genes in BV2 cells treated with different exosomes by transcriptional sequencing (*n* = 3 replicates per group) (N). Analysis of upregulated transcriptome signal pathway by GO pathway between BV2 cells treated with aADEs vs ADEs (O). (P) Relative expression of neurotoxic, neuroprotective and neurotrophic genes in primary astrocytes treated with different exosomes by RT-qPCR (*n* = 3 replicates per group). (Q) Quantification of neuronal death induce by supernatant from WT-ADEs, WT-aADEs, and GSDME KO-aADEs-treated astrocytes. Statistical significance determined using a two-tailed Student’s *t* test. All values are represented as mean ± SEM, \**p* < 0.05; \*\**p* < 0.01; \*\*\**p* < 0.001. See also Figure S5.

Exosomes have emerged as pivotal mediators of neuroinflammation and cognitive decline,^53-55^ prompting investigation into their role in aACM-mediated neurotoxicity. Fractionation of neurotoxic aACM revealed that its exosomal component recapitulated the full neurotoxicity of equivalent aACM volumes (Figure 5D), with effects scaling dose-dependently (Figure S5E). Removal of aADEs from aACM abolished its neurotoxic activity, restoring neuronal viability to ACM levels (Figure 5D), confirming exosomes as primary neurotoxic effectors. To interrogate the role of GSDME, we isolated ADEs from WT-ACM, GSDME KO-ACM, WT-aACM, and GSDME KO-aACM, and confirmed equivalent neuronal uptake across all groups (Figure 5G and S5F). While WT-aADEs reduced neuronal survival compared to WT-ADEs counterparts (Figure 5E), GSDME ablation completely rescued this neurotoxicity. Synaptic analysis through MAP2/SYP immunostaining further demonstrated that WT-aADEs disrupted synaptogenesis more severely than WT-ADEs, an effect entirely reversed by GSDME deficiency (Figure 5F and 5G). These findings indicate GSDME-controlled ADEs as indispensable mediators of pathological neurotoxicity, directly linking GSDME activation in reactive astrocytes to synaptic pathology and neuronal demise.

Exosomes serve as critical carriers of neurotoxic cargoes, including misfolded proteins and inflammatory mediators, directly contributing to the pathogenesis of neuronal injury.^55^ To systematically dissect the neurotoxic molecular components in aADEs, we performed multi-dimensional molecular profiling analyses. Immunoblotting further confirmed differential loading of neurotoxic proteins, including C3, LCN2, HMGB1, iNOS and neuroprotective proteins, GPC4 and BDNF, in WT-aADEs (Figure 5H and 5I). Lipidomic profiling by LC-MS revealed significantly elevated levels of neurotoxic long-chain saturated lipids^56^ in WT-aADEs compared to WT-ADEs (Figure 5J). Concurrently, miRNA profiling demonstrated marked upregulation of neurotoxic miRNAs,^55,57^ such as miR-141-3p, miR-155-5p, miR-135a-5p, miR-125a-5p, and downregulation of neuroprotective miRNAs, such as miR-9-5p, miR-124-3p, miR-132-3p, miR-29b in WT-aADEs (Figure 5K). Notably, transcriptomic analysis of ADEs revealed that WT-aADEs recapitulated the transcriptional signature of their parental cells, characterized by activation of neurotoxic transcripts and suppression of neuroprotective transcripts (Figure S5H). This finding suggests that ADEs may function as biological sentinels reflecting astrocyte reactivity states. Importantly, GSDME deficiency not only inhibited the incorporation of neurotoxic proteins, lipids and miRNAs or mRNAs, but also enhanced the packaging efficiency of neuroprotective miRNAs, mRNAs and selectively enriched neurotrophic factors (Figure 5H-5L). Collectively, these findings reveal that GSDME ablation profoundly reprogrammed the molecular composition of aADEs converting these exosomes from neurotoxic effectors to neuroprotective signaling entities.

Next, we investigated whether the aADEs mediate propagating neuroinflammation. We isolated WT-ADEs, WT-aADEs and GSDME-KO-aADEs and assessed their effects on BV2 microglia and primary astrocytes (Figure 5M). Compared to WT-ADEs, WT-aADEs upregulated pro-inflammatory genes (Figure 5N) and signals in BV2 cells (Figure 5O), while inducing neurotoxic transcripts (Figure 5P) and neurotoxicity (Figure 5Q) of primary astrocytes, suggesting their role as neuroinflammatory propagators. Notably, GSDME ablation suppressed pro-inflammatory gene expression in microglia (Figure 5N) and inhibited neurotoxic phenotype conversion (Figure 5P and 5Q) in astrocytes. These findings position GSDME as a key regulator of aADEs-mediated neuroinflammatory cascades, operating through coordinated control of cytotoxic cargo sorting and intercellular signaling amplification.

### Blocking astrocytic GSDME activation alleviates pathology in 5×FAD mice

Given our prior observation of pathological GSDME activation in astrocytes from both AD patients and 5×FAD transgenic mice (Figure 1), we hypothesized that astrocyte-specific knockdown of GSDME might attenuate Aβ-induced neurodegeneration in 5×FAD mice. To achieve targeted knockdown in astrocytes throughout the developing brain, we employed a PHP.eB-serotype AAV vector, which is capable of crossing the blood–brain barrier to express GSDME-specific shRNA (AAV-shGE) or a non-targeting control shRNA (AAV-shNT) under the GFAP promoter.^58,59^ Neonatal (postnatal day 1–3) mice received stereotaxic intracerebroventricular injections, and all subsequent analyses were performed at 6 months of age (Figure 6A). Whole-brain mapping of viral expression confirmed robust distribution of both constructs and high colocalization with GFAP-positive astrocytes (Figure 6B and 6C), validating our approach for cell-type-specific intervention. Quantification by immunoblotting revealed that total GSDME protein levels were significantly reduced in the hippocampi of AAV-shGE–treated 5×FAD mice compared to the AAV-shNT controls (Figure 6D and 6E). Importantly, the abundance of the cleaved GSDME N-terminal fragment (GSDME-NT) was likewise diminished (Figure 6D and 6E), suggesting effective inhibition of GSDME activation in astrocytes.

**Figure 6.**
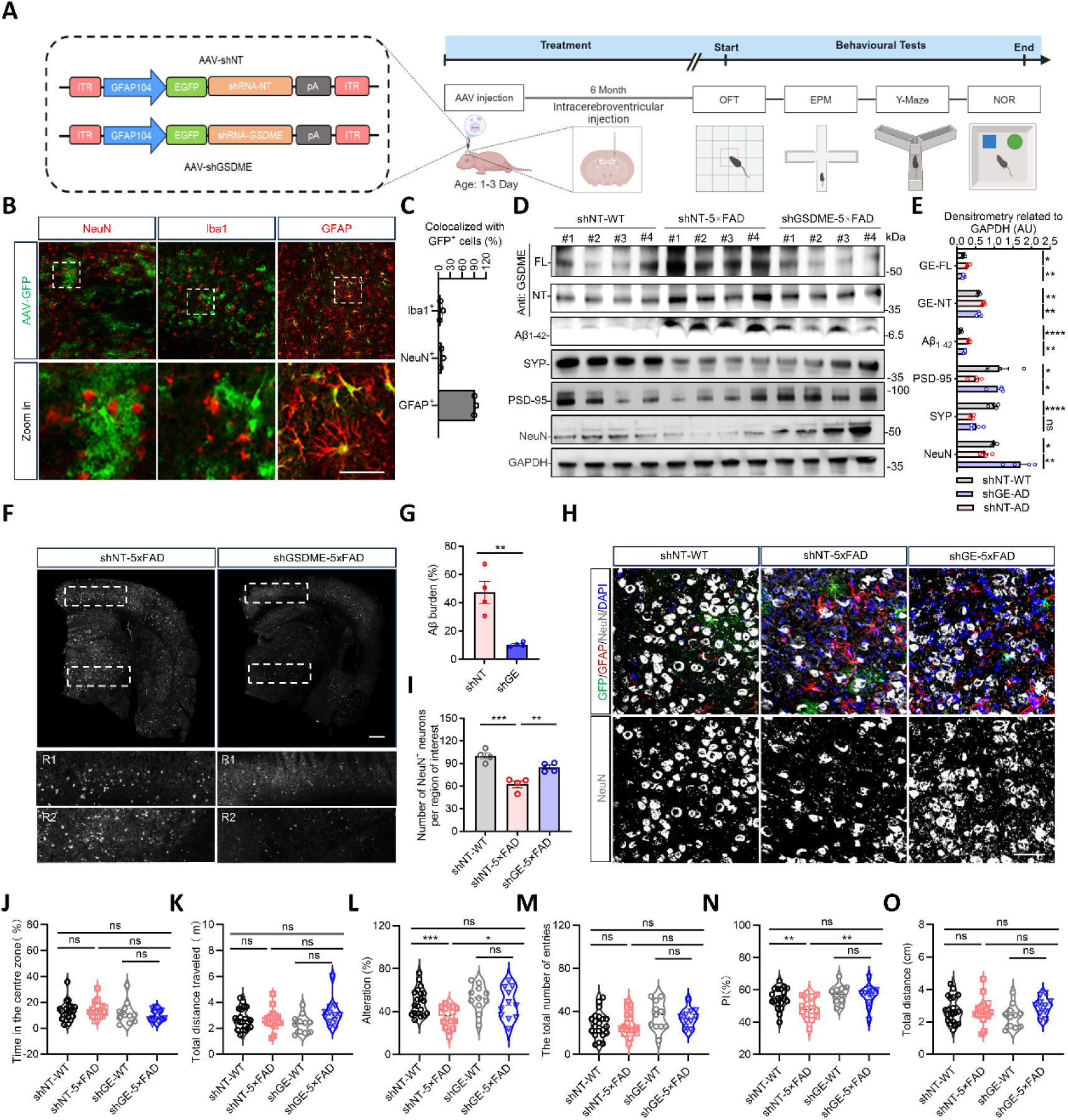
Inhibition of Astrocytic GSDME Activation Attenuates Pathological Progression in 5×FAD Mice. (A) Experimental timeline and group design. Neonatal WT and 5×FAD mice were injected with AAV.PHP.eB-hGFAPS-GFP expressing either a scrambled control shRNA (shNT) or GSDME-targeting shRNA (shGSDME, shGE). Behavioral and pathological assessments were performed at indicated time points. (*n* > 10 mice per group). (B and C) Representative confocal images showing GFP^+^ cells (green) colocalized with NeuN, Iba1 or GFAP positive cells (Red) in the cortex of shNT-WT mice (B). Quantification of the ratio of GFAP⁺GFP⁺, Iba1⁺GFP⁺, and NeuN⁺GFP⁺ double positive cells within total GFP⁺cells in the cortex (*n* = 3 replicates per group) (C). Scale bar, 50 μm. (D and E) Representative immunoblots of cortical tissues from AAV-treated mice showing protein expression levels of GSDME-FL, GSDME-N, Aβ_1-42_, synaptic markers (SYP and PSD-95), neuronal nuclei marker (NeuN), with GAPDH as loading control (D). Quantitative analysis of protein bands by ImageJ: GSDME isoforms (FL/NT), Aβ_1-42_, synaptic proteins (SYP/PSD-95), and NeuN. Data normalized to GAPDH with background subtraction (*n* = 4 mice per group) (E). (D and E) Immunoblot analysis of the activation of GSDME, Aβ_1-42_, SYP, PSD-95 and NeuN in cortical tissue lysates collected from AAV-treated mice (*n* = 4 replicates per group) (D), (E) measured by Image J based on gray scale. (F and G) Representative images of QM-FN-SO3 staining of Aβ plaques in coronal whole-brain sections of shGE and shNT-5×FAD mice (F) and quantification of plaque burden (G). Scale bar, 1000 μm (*n* = 4 replicates per group). (H and I) Representative IF images of sections collected from shGE and shNT-5×FAD mice co-stained with anti-NeuN, anti-GFAP antibodies (H). Number of NeuN-positive cells was quantified using Image J (I).Scale bar, 1000 μm (*n* = 3 replicates per group). (J and K) Open-field test for assessing anxiety-like behavior and locomotor activity. Times in the centre zone (J) and tolal distance traveled (K) (*n* > 10 mice per group). (L and M) Y maze test on the same groups of mice. Spontaneous alternation rate, calculated using the standard method, represent working memory capacity and cognitive flexibility (L).Total entries into all three arms reflect baseline locomotor activity (M) (*n* > 10 mice per group). (N and O) Novel object recognition (NOR) test on the same groups of mice. Object recognition index (ORI), reflecting short-term memory performance (N).Total distance traveled during the test session, indicating baseline locomotor activity (O) (*n* > 10 mice per group). Statistical significance determined using a two-tailed Student’s *t* test. All values are represented as mean ± SEM, \**p* < 0.05; \*\**p* < 0.01; \*\*\**p* < 0.001; \*\*\*\**p* < 0.0001.

To evaluate therapeutic efficacy, we performed neuropathological assessments and behavioral testing six months after injection. Immunoblot analysis showed that GSDME silencing not only reduced Aβ_1-42_ levels but also prevented the downregulation of synaptic markers (Figure 6D and 6E). Consistent with these findings, QM-FN-SO₃ staining, which specifically targets Aβ plaques, revealed a marked reduction in plaque burden in AAV-shGE-treated mice (Figure 6F and 6G), suggesting that astrocyte-specific GSDME silencing suppresses Aβ pathology. NeuN immunostaining further revealed preservation of layer V cortical pyramidal neurons in the AAV-shGE group (Figure 6H and 6I), implying a neuroprotective effect mediated by astrocytic GSDME knockdown.

In the open-field test, no significant differences in locomotor activity were observed among treatment groups (Figure 6J and 6K), ruling out motor dysfunction as a confounding variable for cognitive readouts. In contrast, AAV-shGE-treated 5×FAD mice exhibited significantly improved spatial memory in the Y-maze spontaneous alternation test (Figure 6L and 6M) and displayed greater discrimination in the novel object recognition assay (Figure 6N and 6O). These data collectively demonstrate that astrocyte-specific inhibition of GSDME activation rescues both neuropathology and cognitive deficits in the 5×FAD model, highlighting GSDME as a promising therapeutic target for AD.

### GSDME ablation orchestrates dual reprogramming of astrocyte and exosome reactivity to halt neurotoxic signaling *in vivo*

To examine the potential modulation of astrocyte state transitions by astrocyte-specific GSDME knockdown, we analyzed brain tissues from 5×FAD mice through immunoblotting and transcriptomic sequencing. Immunoblotting revealed that GSDME ablation suppressed neurotoxic factor C3 expression, while upregulating neuroprotective genes, such as S100A10, SHH, GPC4, and BDNF, accompanied by reduced Iba1, and GFAP levels (Figure 7A and 7B), indicating attenuated neuroinflammatory signaling. Mechanistically, astrocytic GSDME knockout restored aberrantly low p-CaMKIIα phosphorylation to wild-type levels (Figure 7C and 7D), a molecular signature linked to improved synaptic plasticity markers^60^ and elevated p-NRF2/p-p65 (Figure 7C and 7D), suggesting its contribution to neuronal protection. Transcriptomics and GSEA further demonstrated suppression of pro-inflammatory mediators such as *Lcn2*, *Nlrc5*, and *Irf7*, alongside upregulation of neuroprotective factors, such as *Sphk1* and *B3GNT5*, as well as neurotrophic factors, *BMP6* and *BMP7* (Figure 7E and 7F). Pathway analysis revealed inhibition of type I interferon and oxidative stress signaling, coupled with activation of neurotransmitter biosynthesis and glial-derived neurotrophic factor synthesis (Figure 7F). qPCR also confirmed downregulated *C3* and *Lcn2*, and upregulated *EMP1* and *Sphk1* mRNA levels in hippocampus (Figure 7G). IF validation confirmed reduced proportions of C3-positive toxic astrocytes (Figure 7H and 7I), increased S100A10 (Figure 7J and 7K) or BDNF-positive (Figure 7L and 7M) protective astrocytes, and diminished microglial infiltration (Figure 7N and 7O), establishing GSDME deletion as a suppressor of astrocyte neurotoxic phenotypic switching *in vivo*.

**Figure 7.**
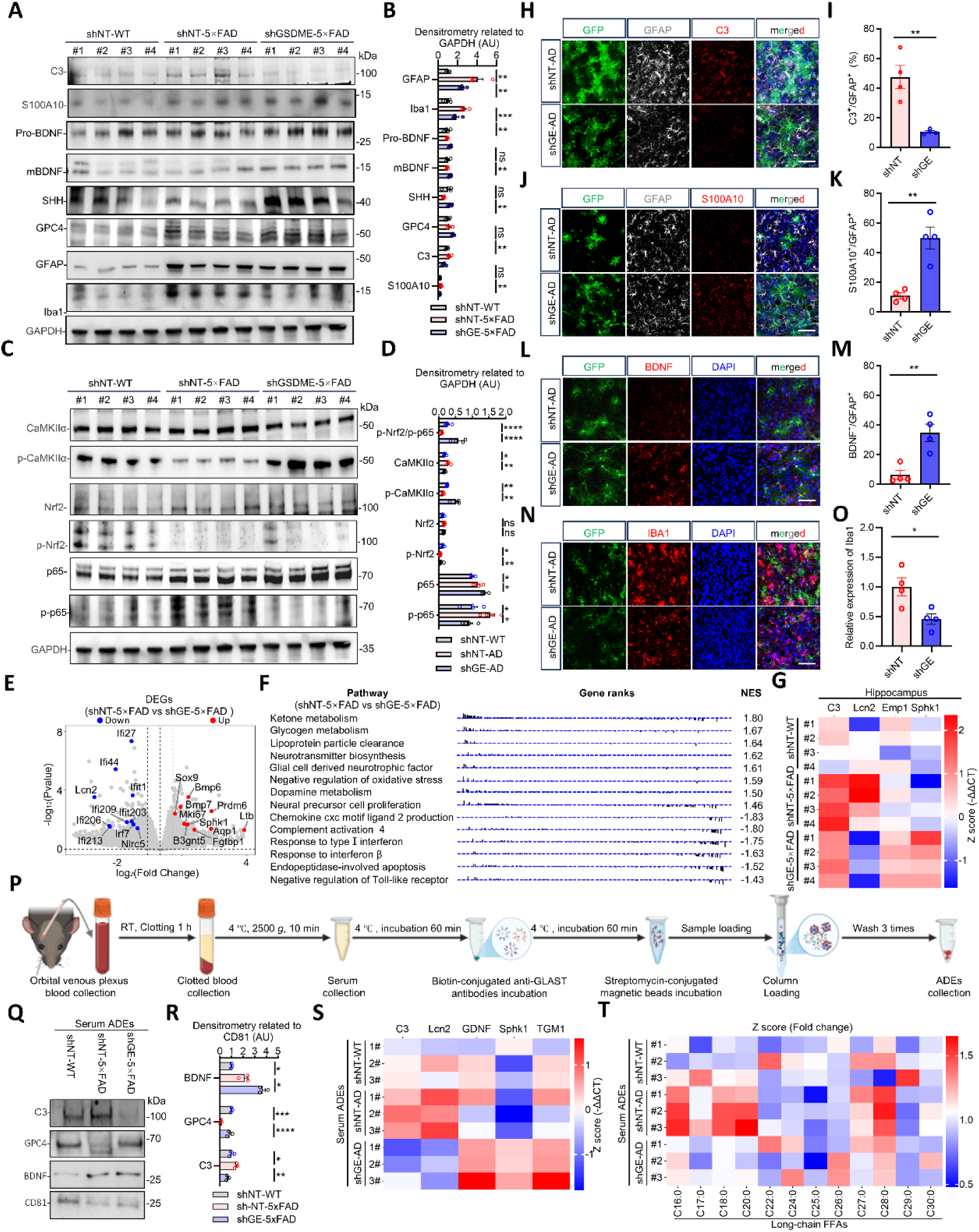
GSDME silencing coordinates biphasic reprogramming of astrocyte-exosome crosstalk to suppress neurotoxic cascade*s in vivo*. (A and B) Immunoblot analysis of C3, S100A10, BDNF, SHH, GPC4, GFAP and Iba1 in cortex tissue lysates collected from shGE and shNT-5×FAD mice (A), (B) measured by Image J based on gray scale (*n* = 3 replicates per group). (C and D) Immunoblot analysis of CaMKIIα, p-CaMKIIα, Nrf2, p-Nrf2, p65 and p-p65 in cortex tissue lysates collected from shGE and shNT 5×FAD mice (C), (D)measured by Image J based on gray scale (*n* = 3 replicates per group). (E) Representative RNA-seq volcano plot of cortical differentially expressed genes (DEGs) between shGE-5×FAD and shNT-5×FAD groups; Red and blue dots indicate significantly upregulated and downregulated genes respectively (|log2FC| > 0.585, FDR < 0.05). Vertical dashed lines mark fold-change thresholds. (F) Cortical pathway enrichment analysis of shGE-5×FAD vs shNT-5×FAD groups by GSEA; Significantly altered pathways are displayed according to normalized enrichment score (NES) absolute values. Horizontal dashed line indicates significance threshold. (G) Hippocampal qPCR validation of dysregulated genes; Heatmap showing relative mRNA expression levels of *C3*, *Lcn2*, *Emp1*, an*d Sphk1* in shGE-5×FAD, shNT-5× FAD, and shNT-WT groups (*n* = 4). Data normalized to *Gapdh* and expressed as log_2_-fold changes relative to shNT-WT controls (2^^-ΔΔCT^ method). (H and I) Representative IF image of sections collected from shGE- and shNT-5×FAD mice cortex co-stained with anti-GFAP, anti-C3 antibodies (H). Percentage of C3-positive cells within GFAP^+^ cell populations were quantified by Image J (n *=* 4 replicates per group) (I). (J and K) Representative IF image of sections collected from shGE- and shNT-5×FAD mice cortex co-stained with anti-GFAP, anti-S100A10 antibodies (J). Percentage of S100A10-positive cells within GFAP^+^ cell populations were quantified by Image J (*n* = 4 replicates per group) (K). (L and M) Representative IF image of sections collected from shGE- and shNT-5× FAD mice cortex co-stained with anti-BDNF antibody (L). Relative expression level of Iba1 was quantified by Image J (*n* = 4 replicates per group) (M). (N and O) Representative IF image of sections collected from shGE- and shNT-5×FAD mice cortex co-stained with anti-Iba1 antibody (N). Relative expression level of Iba1 was quantified by Image J (*n* = 4 replicates per group) (O). (P) Schematic workflow of astrocyte-derived exosomes (ADEs) isolation from mouse serum via orbital venous plexus blood collection. (Q and R) Immunoblot analysis of C3, GPC4, BDNF and CD81 in C3 in ADEs collected from shNT-WT, shGE- and shNT-5×FAD mice (Q), (R) measured by Image J based on gray scale (*n* = 3 replicates per group). (S) Transcriptomic heatmap of ADE-derived differentially expressed genes (DEGs) across shNT-WT, shGE and shNT-5×FAD groups (*n* = 3 replicates per group). (T) Lipidomic heatmap of ADE-losded long-chain fatty acid profiles from the same experimental groups (*n* = 3 replicates per group). Statistical significance determined using a two-tailed Student’s *t* test. All values are represented as mean ± SEM, \**p* < 0.05; \*\**p* < 0.01; \*\*\**p* < 0.001. See also **Figure S6**.

To investigate whether GSDME regulates the reactive state of ADEs *in vivo*, we isolated ADEs from mouse serum for functional validation (Figure 7P and S6). Immunoblot analysis revealed that astrocytic GSDME knockdown suppressed C3 loading in ADEs, while promoting the enrichment of neurotrophic factors, GPC4 and BDNF (Figure 7Q and 7R). The qPCR results demonstrated that astrocytic GSDME ablation significantly reduced transcriptional levels of *C3* and *Lcn2* and upregulated the neuroprotective factor *Gdnf, Tgm, Sphk1* (Figure 7S). Lipidomic analysis further confirmed that GSDME deletion attenuated the production of toxic lipids (Figure 7T). Collectively, astrocyte-specific GSDME knockdown dually suppresses neuroinflammation and restores protective phenotypes intracellularly, while blocking exosomal toxins transfer and boosting reparative signals extracellularly, systemically resetting CNS and peripheral pathology. These findings establish GSDME as a molecular switch that regulates astrocyte-bystander cell communication through ADEs to limit disease progression.

### Multi-omics analysis of human serum astrocyte-derived exosomes uncovers central-to-peripheral immune-regulatory signaling

To determine the clinical roles of ADEs in mediating neuroinflammatory signaling in AD, we performed a comprehensive multi-omics analysis of serum-derived ADEs from AD patients and age-matched healthy controls (Figure 8A). Our transcriptomic analysis revealed that ADEs from AD patients exhibit a distinct neurotoxic molecular signature, characterized by the significant upregulation of neurotoxic factors, including *C3*, *Lcn2*, and *Hmgb1*, alongside a marked downregulation of protective factors like *S100A10*, *Sphk1*, and *Tgm1*, as well as neurotrophic factors such as *Fgf1*, *Nptx1*, and *Ntf3* (Figure 8B). This gene expression pattern closely mirrors the profile of neurotoxic reactive astrocytes observed *in vitro*. Further analysis using the KEGG pathway framework confirmed that ADEs from AD patients are enriched in pro-inflammatory pathways, such as TNF, mTOR, and PI3K-AKT signaling (Figure 8C), while concurrently showing suppression of protective pathways, including JAK-STAT and neurotrophin signaling (Figure 8D). Mass spectrometry analysis corroborated these findings by identifying elevated levels of neurotoxic proteins like LCN2 and B2M, alongside reduced levels of neurotrophic factors, such as THBS1(Figure 8E-8G). Gene Set Enrichment Analysis (GSEA) highlighted significant enrichment in pro-inflammatory pathways, including NF-κB and IL-17 signaling (Figure 8H and 8I). Notably, immunoblotting and single-molecule array quantification indicated elevated levels of LCN2 (Figure 8J), p-Tau181 (Figure 8K), and p-Tau217 (Figure 8L) in AD-ADEs.^61^ Moreover, full-length GSDME protein was detected exclusively in AD-ADEs for the first time (Figure 8M). Lipidomics analysis further showed an enrichment of long-chain saturated fatty acids, such as C16:0 and C18:0, in AD-ADEs (Figure 8N). Taken together, these cross-omics data outline a neurotoxic molecular signature in AD-ADEs, characterized by the synergistic presence of pathogenic proteins, pro-inflammatory signaling, and disrupted lipid metabolism, which may contribute to the propagation of neurodegeneration via neurotoxic and inflammatory amplification.

**Figure 8.**
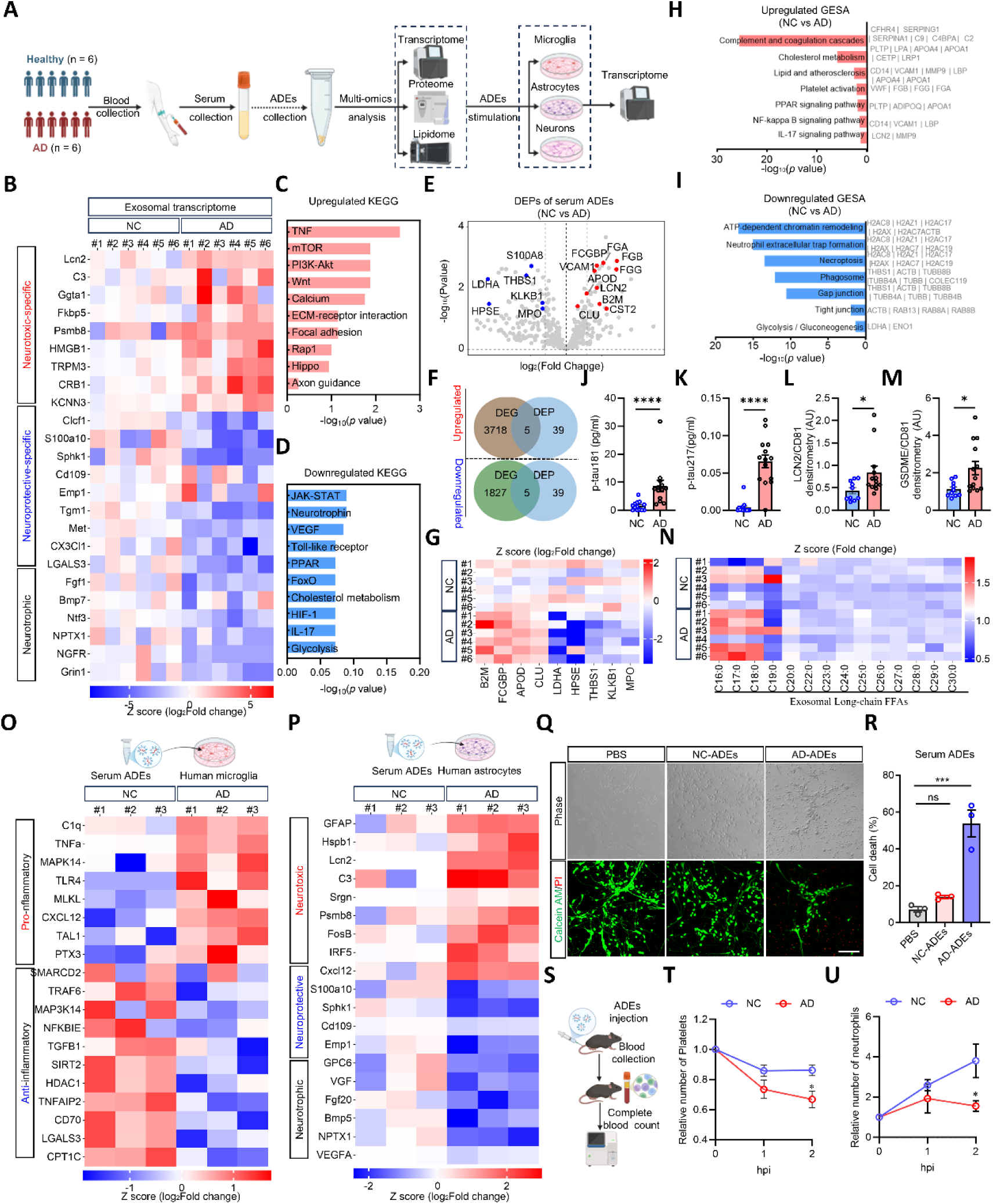
Detecting the roles of AD-ADEs in the modulation of Alzheimer’s pathogenesis. (A) Schematic diagram of multi-omics analysis on serum-derived ADEs. (B) Transcriptome heatmap showing a unique neurotoxic molecular genes profile of AD-ADEs (*n* = 6 replicates per group). (C and D) Analysis of upregulated (C) and downregulated (D) DEGs by KEGG pathway between AD and non-AD (NC) control ADEs. (E) Volcano plot showing elevated neurotoxic proteins and reduced neurotrophic factors from AD-ADEs compared to NC-ADEs (*n* = 6 replicates per group). (F and G) Mass spectrometry reveals co-upregulation and co-downregulation of differentially expressed genes (DEGs) and proteins (DEPs) between NC and AD. (H and I) Gene Set Enrichment Analysis (GESA) showing enrichment in pro-inflammatory pathways of AD-ADEs. (J-M) Immunoblotting and simoa quantification showing elevated LCN2 (J), p-Tau181 (K), p-Tau217 (L) and GSDME-FL (M) in AD-ADEs (*n* = 13 replicates per group). (N) Lipidomic analysis showing enrichment of long-chain saturated fatty acids in AD-ADEs (*n* = 6 replicates per group). (O and P) Transcriptome heatmap of human glial cells simulated by serum AD-ADEs compared to NC-ADEs showing activated of microglia (O) and activated astrocytes (P) *in vitro*. (Q and R) Representative images of primary neuron stimulated with AD-ADEs (Q) and neuronal mortality (R) were detected with Calcein-AM and PI stain, quantified by Image J (*n* = 6 replicates per group) (S) Schematic diagram of the peripheral pro-inflammation role of AD-ADEs in a mouse model. (T and U) The count of blood cells in mice treated with AD-ADEs showing decreased platelets (T) and increased neutrophils (U) 2 h post tail vein injected. Statistical significance determined using a two-tailed Student’s *t* test. All values are represented as mean ± SEM, \**p* < 0.05; \*\**p* < 0.01; \*\*\**p* < 0.001.

To functionally validate the impact of AD-ADEs on neuroinflammatory cascades, we performed a series of in vitro and in vivo assays. *In vitro* stimulation of human microglial cell lines with AD-ADEs induced the transcriptional upregulation of pro-inflammatory mediators^62^, including *C1q*, *Tnfα*, and *Cxcl12*, alongside the suppression of anti-inflammatory regulators such as *NFKBIE*, *Sirt2*, and *HDAC1*, suggesting the activation of microglia (Figure 8O). In parallel, AD-ADEs triggered a neurotoxic reactive state in human primary astrocytes, illustrating their ability to transmit neuroinflammatory signals across glial populations (Figure 8P). Moreover, AD-ADEs significantly increased neuronal mortality compared to control exosomes (NC-ADEs), confirming their direct neurotoxic effects (Figure 8Q and 8R). Considering the peripheral origin of serum ADEs, we also investigated their systemic immunomodulatory effects. Intravenous administration of AD-ADEs in mice resulted in a specific reduction in platelet and neutrophil counts (Figure 8S-8U), highlighting their potential to modulate peripheral immune responses. Collectively, our findings suggest that AD-ADEs orchestrate a dual pathological mechanism, propagating neuroinflammation and neurotoxicity within the central nervous system, while exacerbating systemic pathology through peripheral immune dysregulation.

## DISCUSSION

Our study redefines GSDME as a calcium-dependent molecular rheostat that orchestrates astrocyte and ADE state transitions through spatiotemporal regulation of ER calcium flux at MAMs. Unlike its canonical roles in anti-infection and tumor immunity^63^, GSDME-NTs localize to MAMs, where they induce ER calcium leakage without plasma membrane pore formation. This subcellular activity drives biphasic CaMKIIα phosphorylation dynamics that early-phase activation enhances NRF2-mediated defenses (neuroprotective state), while sustained phosphorylation paradoxically amplifies NF-κB-driven neuroinflammation (neurotoxic state). These findings align with the prevailing consensus that astrocytes can adopt diverse substates in response to various pathological stimuli, extending beyond the simplistic A1/A2 or neurotoxic/neuroprotective framework.^19^ Notably, genetic ablation of GSDME stabilizes the neuroprotective phenotype, as evidenced by reduced expression of neurotoxic markers, such as C3 and LCN2, elevated expression of neurotoxic markers, such as EMP1, SPHK1 and GPC4 levels, while rescuing neuronal viability and cognitive deficits in 5×FAD mice. This mechanistic cascade positions GSDME as a druggable regulator of astrocyte plasticity, which provides a mechanistic foundation for precision therapeutics targeting glial state transitions in AD.

Recent studies have demonstrated that GSDM-NTs target not only the plasma membranes but also diverse organelle membranes.^64-66^ Here, our work demonstrates that GSDME-NT is preferentially localized at MAMs in neurotoxic astrocytes. MAMs, lipid rafts in the endoplasmic reticulum near mitochondria, are crucial for cell health by regulating intracellular Ca^2+^ transfer.^67,68^ They regulate Ca^2+^ release from the ER via IP3Rs and its entry into mitochondria through VDACs, while also balancing cytosolic Ca^2+^ with the MCU complex for uptake and NCLX for release.^68^ Although dysfunctions in MAMs are linked to various pathological conditions and neurodegenerative diseases,^69,70^ their precise role in AD pathogenesis remains inadequately understood. The targeted relocation of GSDME-NTs to MAMs results in ER calcium leakage, as opposed to forming pores in the plasma membrane. Thus, we hypothesize that GSDME-NTs permeabilize MAMs in neurotoxic astrocytes, thereby facilitating the transfer of Ca^2+^ from the ER to both the cytosol and mitochondria. This specific localization mechanism elucidates why the activation of GSDME as a cytosolic calcium channel does not trigger pyroptosis but instead promotes transcriptomic alterations by modulating calcium signaling pathways. This subcellular targeting may constitute a common characteristic within the GSDM family, offering a novel framework for understanding the diversity of their pathological roles.

Astrocytic calcium dysregulation, a hallmark of early AD, leads to synaptic issues and cognitive decline through reactive astrogliosis.^17,71^Although normalizing Ca²⁺ transients can reverse cognitive deficits in AD mice,^72,73^ the specific mechanisms connecting altered calcium signaling to astrocyte-induced neurotoxicity are not yet understood. Here, we elucidate a temporal rheostat mechanism wherein calcium-dependent phosphorylation dynamics of CaMKIIα govern the transition from neuroprotective to neurotoxic through phase-specific transcriptional reprogramming. During the early phase, CaMKIIα phosphorylation enhances NRF2-mediated antioxidant defenses, thereby promoting neuroprotection. Conversely, prolonged phosphorylation paradoxically intensifies NF-κB-driven neuroinflammation, converting astrocytes into a neurotoxic state. This biphasic regulation challenges the traditional view of CaMKIIα as a solely neuroprotective kinase and underscores the importance of timing in therapeutic interventions.^60^ GSDME is identified as a crucial temporal checkpoint, its activation triggers ER Ca²⁺ leakage, initiating NF-κB signaling. Interestingly, GSDME knockdown selectively inhibits late-phase CaMKIIα phosphorylation, thereby preventing NF-κB hyperactivation and subsequent neurotoxicity. These findings align with broader calcium signaling dynamics, where transient vs. sustained calcium signals differentially regulate synaptic plasticity and cellular homeostasis. Consistently, pharmacological testing with antagonists shows a time-dependent dual effect that early CaMKIIα activation is protective, whereas late inactivation hampers toxicity, indicating metabolic feedback with distinct neuroprotective outcomes. Taken together, our work reconceptualizes astrogliosis as a calcium-regulated process orchestrated by the dual functions of CaMKIIα. By modulating its activity in a phase-specific manner, we propose a “late inhibition” strategy aimed at mitigating pathogenic astrogliosis through targeted p-CaMKIIα. This methodology integrates AD pathomechanisms with therapeutic development, presenting a precision neuroprotection framework for early intervention in neurodegenerative disorders.

A growing body of evidence positions exosomes as critical mediators of neuropathological progression, with ADE serving as potent coordinators of multicellular neurodegeneration.^74-77^ Unlike prior models attributing astrocyte-mediated neuronal damage to direct cytokine, toxic lipid release or oxidative stress,^56,78,79^ our findings reveal that ADEs function as context-sensitive pathogenic platforms whose cargo dynamically adapts to neurotoxic microenvironments. Multi-omics profiling demonstrates that ADEs from reactive astrocytes exhibit enrichment in neurotoxic lipids, pro-apoptotic miRNAs, and pathogenic tau isoforms relative to homeostatic astrocytes, while co-encapsulating mRNA cargo that coordinately disrupts synaptic architecture, astrocyte to pro-inflammatory microglial activation, and astrocyte-to-astrocyte reactive phenotype propagation. This suggests a tripartite mechanism unprecedented in classical single-pathway neurotoxicity models. Central to this paradigm shift is the identification of GSDME as a molecular switch regulating ADE pathogenicity. Genetic ablation of GSDME prevents reactive astrocyte transition from neuroprotective to neurotoxic state, while reprogramming ADE cargo toward neuroprotective mediators, increasing BDNF,^80^ GPC4 levels^81^ and enhancing anti-inflammatory miRNAs. These findings redefine GSDME as a master regulator of astrocyte exosome biology, further extending its canonical role in pyroptosis to include ADE-mediated intercellular communication, and position ADEs as critical mediators of neuroinflammation propagation.

While early intervention remains pivotal for AD therapeutics,^82^ current limited glial markers, including serum GFAP, lack resolution to decode the functional heterogeneity of reactive astrocytes driving AD progression.^11,24^ Here, we pioneer a multi-dimensional serum ADE biomarker platform that integrates transcriptomic, lipidomic, and proteomic profiling to transcend static tissue paradigms. This strategy identifies an AD-specific molecular fingerprint, including coordinated depletion of neuroprotective transcripts, amplification of pro-degenerative effectors, and lipidomic dysregulation marked by C16:0, C18:0 elevation. Unlike conventional methods, this cross-omics framework captures dynamic astrocyte state transitions through serum ADE-mediated intercellular networks, enabling presymptomatic stratification with mechanistic granularity. By bridging real-time molecular pathology and therapeutic targeting, ADE multi-omics repositions biomarker discovery as a mechanism-to-therapeutics pipeline, where early diagnosis directly informs astrocyte-modulating precision strategies.

In summary, our work redefines astrocyte activation as a calcium-regulated continuum. The discovery of GSDME-NTs localized at MAM and its dual regulation of cell-autonomous signaling via CaMKIIα and non-cell-autonomous communication through ADEs provides a unified framework for understanding glial-driven neurodegeneration. This paradigm shift integrates calcium dynamics, transcriptional reprogramming, and intercellular communication, offering novel insights into AD pathogenesis. Future studies should explore the conservation of GSDME-MAM interactions in human AD progression and investigate the therapeutic potential of combinatorial strategies targeting both calcium signaling and ADE-mediated inflammation.

### Limitations of the study

Although our study establishes GSDME as a critical regulator of astrocyte state transitions, several limitations warrant further investigation. First, the molecular mechanisms driving GSDME-NT’s preferential localization to MAMs remain incompletely defined; whether specific lipid modifications or chaperones mediate this subcellular targeting requires systematic interrogation. Second, the therapeutic validation is restricted to Aβ pathology models, lacking examination in Tau-inclusive models like 3×Tg, which may limit generalizability to AD with mixed pathologies. Third, our ADE multi-omics approach, while identifying disease-specific signatures, does not resolve heterogeneity among ADE subpopulations, which may obscure functionally distinct cargo subsets. Single-vesicle profiling and spatial transcriptomics may address this limitation. Fourth, the long-term consequences of GSDME ablation on physiological processes remain unexplored, raising potential safety concerns for therapeutic targeting. Finally, human tissue validation of GSDME associations is limited to postmortem AD samples; future studies should employ PET-MRI imaging to correlate GSDME activity with real-time neuroinflammation in living patients. Addressing these gaps will refine our understanding of the spatiotemporal regulation of GSDME and accelerate translation of astrocyte-targeted therapies.

## Acknowledgments

We thank Dr. M. Xu from the Center for Brain-Inspired Intelligence at the Chinese Academy of Sciences for the comments. We thank Dr. J. Xiang and Dr. Y.Z. Wang from the Air Force Medical University for the helpful advices. We thank the analysis & testing laboratory for life sciences and medicine of the Air Force Medical University for their assistance with electron microscopy imaging, as approved by Y. Zhao, and structured illumination microscopy imaging, as approved by D.L. Shi. We thank the Chinese Brain Bank Center for providing the human brain samples.

## Funding

Funding was obtained from the National Key Research and Development Plan (No.2022YFC3400103 to Q.L. and X.B.H.), the ECUST-OPM Open Fund (No. 20220701 to Q.L.).

## Author contributions

L. and S.W. C., X.B. H. conceived the project; Q. L., S.W. C., X.M. X designed the experiments and wrote the manuscript. Q. L., S.W. C., X.B. H., X.M. X, J.M. X., X.Q. L., X.Y. L., X.M. W. and Y.X. Z. revised the manuscript. S.W. C., J.M. X., X.Q. L., X.Y. L. and X.M. W., S.X. L. and Y.X. G. performed the experiments with the assistance from Y.Z. C., G.G. G.; and Q. L., S.W. C. and X.M. X, J.M. X., X.Q. L., X.Y. L., X.M. W. analyzed the data and created the figures. All authors discussed the results and commented on the manuscript.

## Competing interests

The authors declare that they have no competing interests.

## STAR Methods

Detailed methods are provided in the online version and include the following:

● KEY RESOURCES TABLE
● RESOURCE AVAILABILITY
  ○ Lead contact
  ○ Materials availability
  ○ Data and code availability
● EXPERIMENTAL MODEL AND SUBJECT DETAILS
  ○ Animals
  ○ Cell culture and treatment
● METHOD DETAILS
  ○ Single-cell transcriptome analysis using open snRNA-seq data
  ○ RNA extraction and RNA-seq analysis
  ○ Isolation of astrocytic exosomes
  ○ Nanoparticle tracking analysis (NTA)
  ○ Synaptosome engulfment assay
  ○ Neurotoxicity assay
  ○ Scanning electron microscope (SEM)
  ○ Transmission electron microscopy (TEM)
  ○ Immuno-electron microscope (IEM)
  ○ Western blotting
  ○ Immunostaining
  ○ Characterization of subcellular Ca^2+^
  ○ Saturated lipid analysis
  ○ Behavioral test
  ○ siRNA transfection
  ○ Lentivirus preparation and infection
  ○ Peripheral effect of ADVs *in vivo*

## Supplementary information

**Movie 1.** Detection of cytosolic Ca^2+^ dynamics in WT-MCM, WT-AMCM and GSDME KO-AMCM astrocytes following the stimulation within 8 h, related to Figure 3

**Movie 2.** Detection of ER Ca^2+^ dynamics in WT-MCM and WT-AMCM astrocytes following the stimulation within 8 h, related to Figure 3

**Movie 3.** Detection of mitochondrial Ca^2+^ dynamics in WT-MCM and WT-AMCM astrocytes following the stimulation within 8 h, related to Figure 3

## KEY RESOURCES TABLE

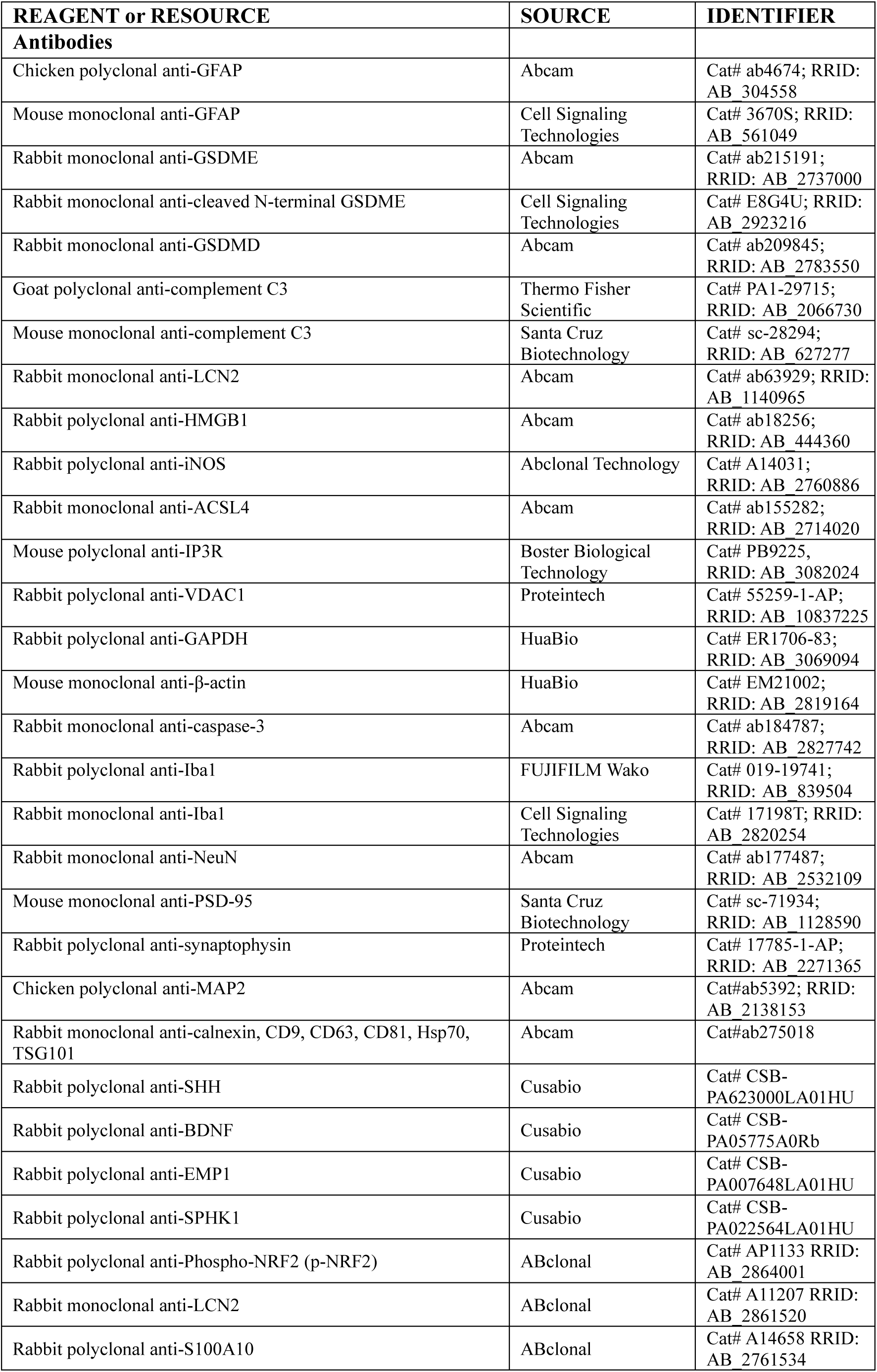
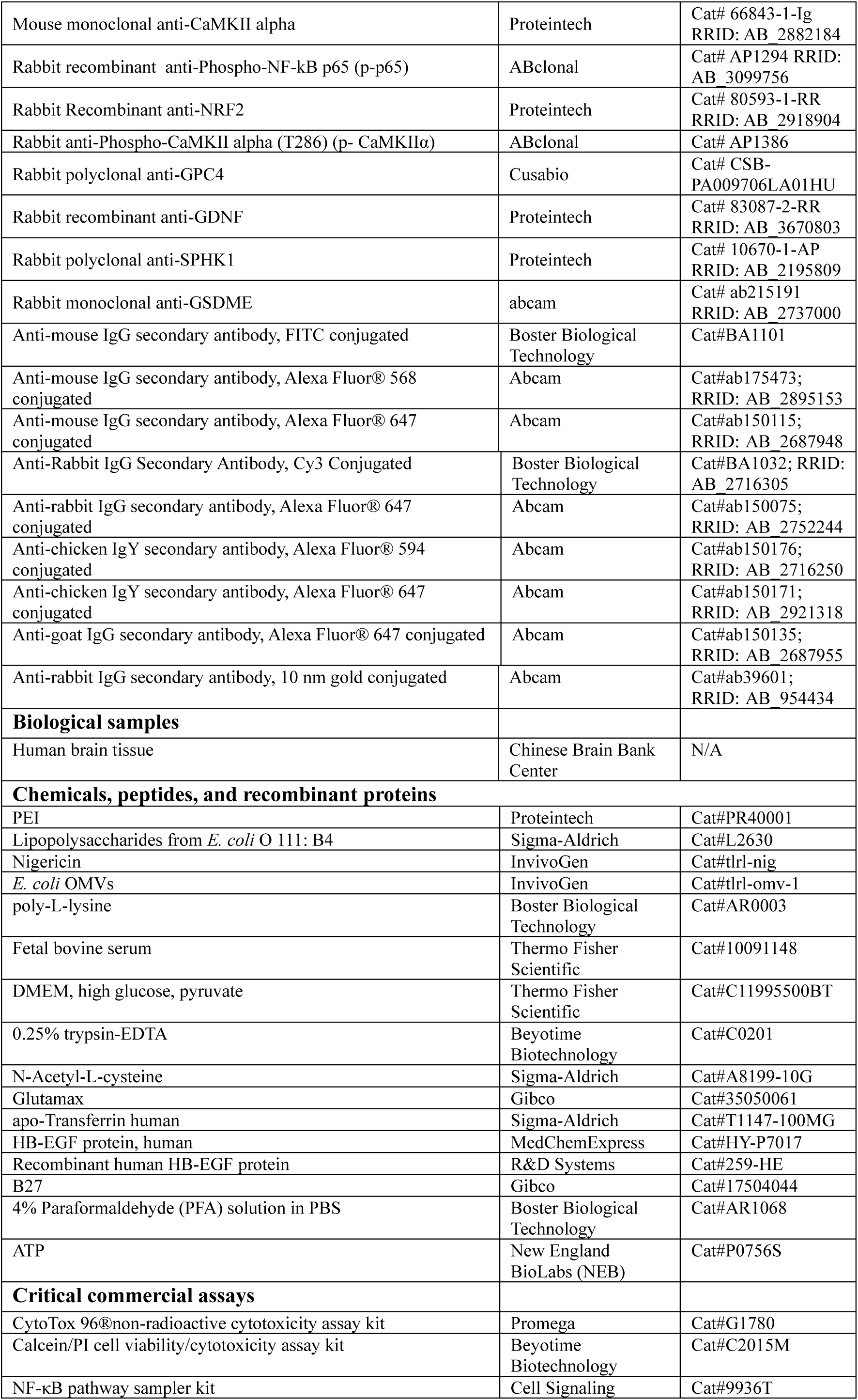
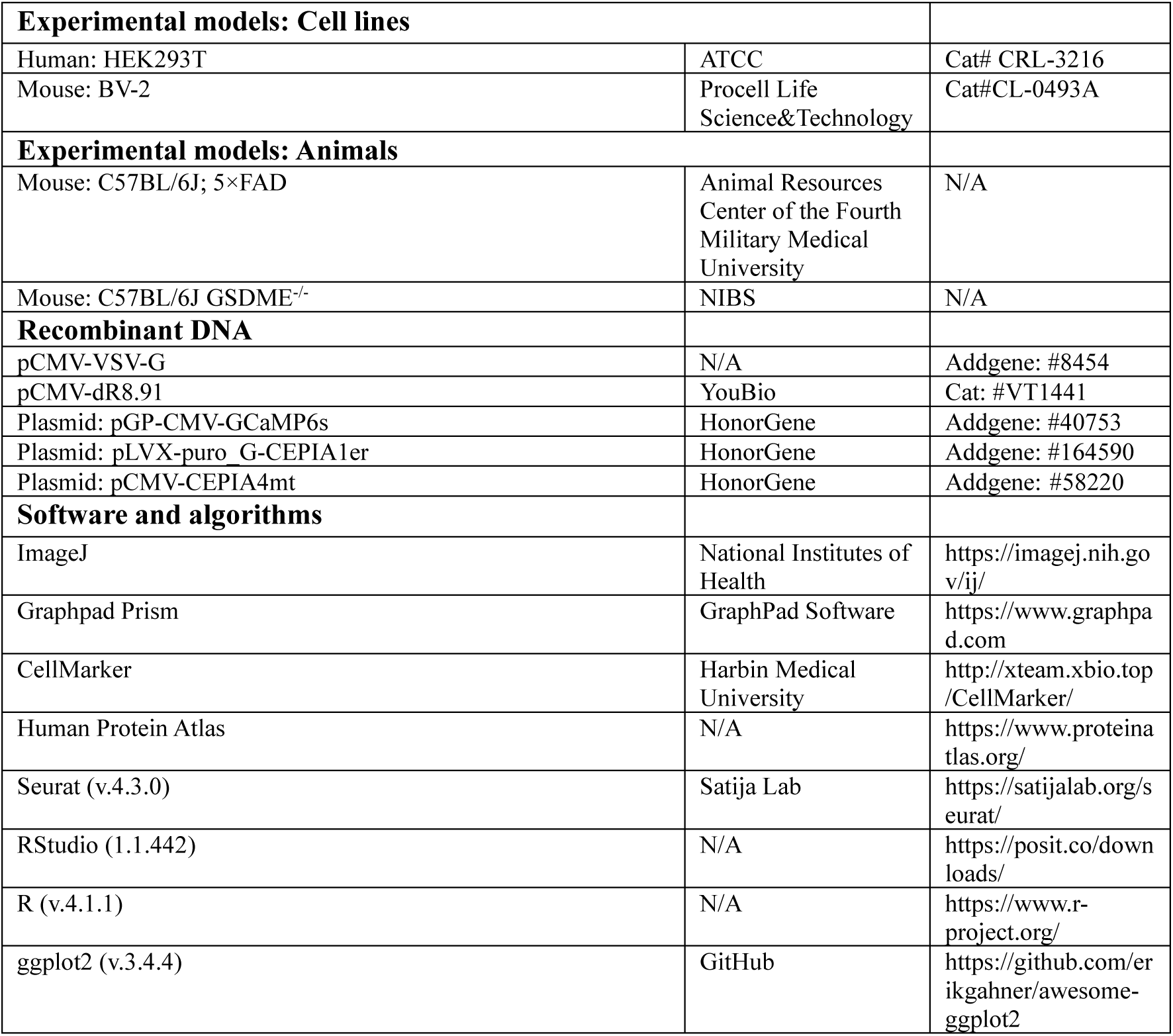

## RESOURCE AVAILABILITY

### Lead contact

Further inquiries and requests for resources and reagents should be directed to the lead contact, Doctor Qin Liu, and will be fulfilled upon reasonable request.

### Materials availability

This study did not generate new unique reagents.

### Data and code availability

● Data reported in this paper will be shared by the lead contact upon request.
● All scripts used in this manuscript are available from the lead contact upon reasonable request.
● Any additional information required to reanalyze data reported in this paper is available from the lead contact upon request.

## EXPERIMENTAL MODEL AND SUBJECT DETAILS

### Animals

All experimental procedures involving mice were conducted under a protocol approved by the Fourth Military Medical University Institutional Animal Care and Use Committee, adhering to the guidelines outlined by the Animal Welfare Act and relevant Chinese regulations concerning animal experimentation. Wild-type C57BL/6 mice and 5×FAD transgenic mice (C57BL/6 background) were supplied by the Animal Resources Center of the Fourth Military Medical University (Xi’an, China), while GSDME^-/-^transgenic mice (C57BL/6) were kindly provided by Professor Feng Shao. All strains were maintained under specific pathogen-free conditions by breeding with C57BL/6 mice, with animals housed in controlled environments (temperature: 22 ± 1°C, humidity: 50 ± 10%, 12-h light/dark cycle) and provided ad libitum access to food and water. To eliminate bias, animals were randomly assigned identification numbers and subsequently evaluated in a blinded manner throughout the experimental procedures.

### Cell culture and treatment

#### Astrocytes

Mouse primary astrocytes were cultured following an established protocol with minor modifications.^83^ Briefly, astrocytes were isolated from the cortices or hippocampus obtained from mouse pups of mixed sexes at postnatal day 0 (P0). P0 mouse pups were decapitated, and brains were isolated and kept in ice-cold Dulbecco’s modified Eagle medium (DMEM). After removing the meninges and capillaries, the cortices or hippocampus were minced and digested with 0.125% Trypsin-EDTA at 37°C for 10 min. Digestion was halted with DMEM containing 10% fetal bovine serum (FBS). Large tissue aggregates were removed by filtering through a 40-μm mesh. The cells were pelleted by centrifugation at 300 × *g* for 10 min and resuspended in DMEM with 10% FBS. Following a differential adhesion period of 20 min, the culture medium was replaced with a serum-free base medium. The culture medium consisted of a mixture of 50% Neurobasal and 50% DMEM, supplemented with 100 U/mL penicillin, 100 μg/mL streptomycin, 1 mM sodium pyruvate, 292 μg/mL glutamine, 5 ng/mL HBEGF, and 5 μg/mL N-acetylcysteine. Cells were seeded on T75 culture flasks pre-coated with poly-L-lysine. After 4 d, the medium was replaced with DMEM supplemented with 10% FBS.

For the collection of astrocyte-conditioned medium (ACM), astrocyte cultures were randomly chosen as control or reactive. Following 8 d in culture, the flask was agitated at 260 rpm for 8 h at 37°C to eliminate microglial cells and then harvested by trypsinization using 0.25% Trypsin-EDTA. Neurotoxic astrocytes were generated *in vitro* by culturing purified astrocytes for 8 d, followed by a 48-h exposure to activated microglia conditioned medium (AMCM). The conditioned medium from both control (ACM) and neurotoxic (aACM) astrocytes was harvested and concentrated using ultrafiltration concentration tubes with a 10 kDa retention capacity until it reached approximately a tenfold volume concentration for subsequent neuronal toxicity testing or exosome isolation.

Primary human astrocytes (HA, ScienCell Research Laboratories, Cat# HA) were cultured in poly-L-lysine-coated (2 μg/cm²) 24-well plates. Prior to cell seeding, plates were pre-coated with poly-L-lysine (10 mg/ml, Cat# 0413) diluted in sterile water (2 μg/cm²) and incubated overnight at 37°C. Cells were thawed and expanded in T-75 flasks following the manufacturer’s protocol, then subcultured into 24-well plates at a density of 5,000 cells/cm² in complete Astrocyte Medium (AM, Cat# 1801) supplemented with AGC and FBS. Cultures were maintained at 37°C in a 5% CO₂ incubator until reaching 70% confluency (typically 3-4 days post-seeding). For exosome stimulation, cells were gently washed twice with pre-warmed DPBS (Cat# 0303) to remove serum and growth factors. Serum- and AGC-free Astrocyte Medium (500 μL/well) was added, followed by exosomes resuspended in the same medium at a final concentration of 100 μg/mL. Control wells received vehicle solution (equal volume of PBS or exosome-free medium). The plate was incubated under standard conditions (37°C, 5% CO₂) for 24 h.

#### Microglia

Mouse primary microglia were cultured following an established protocol with minor modifications.^84^ Briefly, microglia were isolated from the cortices obtained from mouse pups of mixed sexes at postnatal day 0 (P0). P0 mouse pups were decapitated, and brains were isolated and kept in ice-cold DMEM. After removing the meninges and capillaries, the cortices were minced and digested with 0.125% trypsin-EDTA at 37°C for a duration of 15 min. The digestion process was halted with DMEM containing 10% FBS. Large tissue aggregates were removed by filtering through a 40 μm mesh. The cells were pelleted at 600× *g* for 10 min and resuspended in DMEM with 10% FBS. Following a differential adhesion period of 1 h, the culture medium was replaced with a serum-free base medium. The medium was changed the next day and changed every 4 d for the next 12 d. When mixed glial cultures were completely confluent, microglia were harvested by shake-off at 200 rpm at 37°C for 6 h. Microglia were counted and plated in 24-well culture plates at a density of 1.5×10^6^ cells/ml. The culture purity was >99%, verified by immunofluorescence with GFAP (for astrocytes), NeuN (for neurons) and CD11b (for microglia).

#### Neurons

Primary cortical neurons from mouse embryos at embryonic day 14-18 were isolated and cultured in neurobasal medium following a previously established protocol with minor modifications.^85^ Briefly, following a differential adhesion period of 4 h in DMEM containing 10% FBS, the culture medium was replaced with neurobasal medium supplemented with 1% B-27, 0.5 mM glutamine and 1% penicillin-streptomycin. Following 7 d in culture, neurotoxicity assay was conducted *in vitro*.

#### BV2 cells

Immortalized mouse BV2 microglial cells were cultured in DMEM medium with 10% FBS at 37°C in a humidified atmosphere containing 95% air and 5% CO₂. The BV2 cells were stimulated with 0.1 μg/mL lipopolysaccharide (LPS) in Opti-MEM for 4 hours. After LPS stimulation, 5 μM Aβ_1-42_ oligomers were added to the culture and incubated for an additional 20 h. The activated culture medium of BV2 cells (AMCM) was then collected and concentrated to approximately one-tenth of its original volume using 30 kDa ultrafiltration tubes for generating neurotoxic astrocytes.

## METHOD DETAILS

### Single-cell transcriptome analysis using open snRNA-seq data

Basic processing and image plotting of the snRNA-seq data were performed with Seurat (v.4.3.0), ggplot2 (v.3.4.4) and ggpurb (v.0.4.0), run on RStudio (v.1.1.442), using R (v. 4.1.1). Cells with the number of genes (nFeature_RNA) less than 200 and more than 10,000, as well as the percentage of mitochondrial genes larger than 5%were discarded. The expression matrix of snRNA-seq were converted into a standard input file. Following clustering, astrocytes were distinguished from different clusters using CellMarker (http://xteam.xbio.top/CellMarker/) and the Human Protein Atlas (https://www.proteinatlas.org/). Subsequently, the comparation of astrocytic *GSDME* expression (disease group versus healthy group) was identified using the FindMarkers function. Analyses of datasets from various databases were conducted separately. Identification of astrocytes using conventional snRNA seq analysis methods. Further subgroup analysis of astrocytes with a resolution ≥ 0.1. Within each cluster, we assessed the average log_2_ (fold change) of *Gfap*. Clusters with an avg_log_2_(fold change) > 0 were categorized as the reactive astrocyte group, while the remaining clusters were classified as the non-reactive astrocyte group. The cells were discarded with zero expression levels of GSDME. The analysis of astrocytic *GSDME* expression across various subtypes in Figure 6A was conducted post-clustering. Each subtype of astrocytes (ASTs) was subsequently paired with its counterpart to identify ASTs based on marker genes^67^. The correlations between *GSDME* and different astrocytic marker genes across different ASTs were analyzed using the Spearman method.

### RNA extraction and RNA-seq analysis

Total RNA was isolated from mouse cortical tissues using Trizol Reagent (Invitrogen Life Technologies), followed by direct lysis of cell samples in 24-well culture plates. RNA concentration, purity, and integrity were assessed via NanoDrop spectrophotometry (Thermo Scientific) and agarose gel electrophoresis. For library construction, 3 μg of RNA was subjected to mRNA enrichment with poly-T oligo-attached magnetic beads, followed by fragmentation in Illumina buffer using divalent cations (94°C, 5 min). First-strand cDNA was synthesized with SuperScript II reverse transcriptase and random hexamers, and second-strand cDNA was generated using DNA Polymerase I/RNase H. Blunt-end repair, 3′-adenylation, and Illumina adapter ligation were sequentially performed. Library fragments (400–500 bp) were size-selected with AMPure XP beads (Beckman Coulter, USA), amplified by 15-cycle PCR, and quantified via Agilent Bioanalyzer 2100 (Agilent Technologies). Final libraries were sequenced on an Illumina NovaSeq 6000 platform (Shanghai Personal Biotechnology Co., Ltd).

RNA sequencing was performed as follows: Raw sequencing data (FASTQ files) were quality-controlled using Cutadapt (v1.15) to remove adapters and low-quality reads, generating clean data. HISAT2 (v2.0.5) aligned filtered reads to the reference genome. Gene expression quantification was calculated via HTSeq (0.9.1) with FPKM normalization. Differential expression analysis employed DESeq (v1.39.0) with thresholds set at |log_2_FoldChange| >1.5 (tissue samples) or >2 (cellular samples) and adjusted *p*-value <0.05. Bidirectional clustering of differentially expressed genes (DEGs) was visualized using the pheatmap R package (v1.0.8), with Euclidean distance and complete linkage algorithms. Functional enrichment analysis encompassed GO term enrichment via topGO (v2.40.0) using a hypergeometric test (*p*<0.05), KEGG pathway annotation using clusterProfiler (v3.16.1) (*p*<0.05), and multi-database Gene Set Enrichment Analysis (GSEA) performed with clusterProfiler (v3.16.1). For GSEA, pre-ranked gene lists (sorted by log_2_FoldChange) were tested against gene sets from MSigDB Hallmark, GO Biological Process, and KEGG pathways. Significance for GSEA was defined as FDR <0.25 and |Normalized Enrichment Score (NES)| >1.

### Isolation of astrocytic exosomes

The ACM of WT-MCM, GSDME KO-MCM, WT-AMCM, or GSDME KO-AMCM astrocytes were harvested 48 h post-stimulation. The astrocytic exosomes were then isolated through differential ultracentrifugation, following the method previously described.^86^ Briefly, the collected medium underwent centrifugation at 300× *g* for 10 min and 2,000× *g* for 10 min at 4°C to eliminate dead cells and cellular debris. Subsequently, the supernatant was centrifuged at 10,000× *g* for 30 min at 4°C to remove larger microvesicles and apoptotic bodies. The remaining supernatant was subjected to ultracentrifugation at 100,000× *g* for 70 min at 4°C to yield the astrocytic exosome pellet. This pellet was washed once with PBS to ensure the removal of non-specific proteins and debris.

### Nanoparticle tracking analysis (NTA)

The size and quantity of isolated astrocytic exosomes were assessed using a ZetaView Nanoparticle Tracker and Particle Metrix software. Exosome samples were diluted with 1× PBS buffer for size and concentration measurements. The NTA measurements were taken at 11 different positions, and the ZetaView system was calibrated with 250 nm polystyrene particles. The temperature during the measurements was controlled at approximately 23°C and 30°C.

### Synaptosome engulfment assay

Synaptosomes were isolated from mouse brain according to the procedure described in a previous study^87^ and labeled with pHrodo Red succinimidyl ester. Excess pHrodo Red was removed by centrifugation, and the pHrodo-labeled synaptosomes were then resuspended in PBS containing 5% dimethyl sulfoxide (DMSO) before being stored at -80°C. Primary mouse astrocytes were seeded on Poly-L-lysine precoated 24-well plates at a density of 1×10^5^ cells per well for 24 h and exposed to MCM or aMCM for an additional 48 h. Subsequently, the cells were incubated with pHrodo-labeled synaptosomes (5 μL) for 24 h, followed by co-staining with Calcein AM to visualize the uptake of synaptosomes by astrocytes. Images were captured with a confocal microscope (IXplore SpinSR).

### Neurotoxicity assay

Mouse primary cortical neurons were isolated and cultured as mentioned before. The cultured cortical neurons were incubated with the concentrated ACM or purified exosomes collected from astrocytes in neurobasal medium for 24 h. Neuronal viability was measured using the Calcein AM/PI method with a viability/cytotoxicity assay kit (Beyotime, Cat#C2015M).

### Scanning electron microscope (SEM)

Astrocytes growing on cover slips were stimulated by MCM, AMCM or overexpressed EGFP-tagged GSDME-NT. Subsequently, the cells were rinsed thrice with PBS, followed by fixation with 2.5% glutaraldehyde at 4°C for 24 h. The slips were rinsed twice with PBS after glutaraldehyde removed. The morphology of astrocytes on glass slides was examined using a scanning electron microscope (ThermoFisher Quattro S) following treatment with osmium tetroxide, dehydration, and gold coating. For the exosome samples, 20 μL of exosome suspension droplets were pipetted into the copper mesh grid and retained on the copper mesh for a few moments (more than 1 min). Negative staining was performed by dripping 2% phosphotungstic acid onto the grid for 1-10 min, and dry naturally at room temperature after the filter paper is blotted dry. Samples were observed and photographed under a biological transmission electron microscope at 120 kV accelerating voltage.

### Transmission electron microscopy (TEM)

Astrocytes were treated with 0.25% trypsin at 24 h post-stimulation. Subsequently, the cells were pelleted by centrifugation at 600 rpm for 3 min, and resuspended in 3% glutaraldehyde. The resuspended cells were then centrifuged at 5000 rpm for 10 min to form tight cell clusters, after which the supernatant was carefully removed. Fresh glutaraldehyde was slowly added to the tubes, which were then stored at 4°C for 24 h. The cell clusters underwent osmium acid treatment, dehydration, and embedding, and were sectioned to a thickness of 70 nm. Finally, the sections were stained with lead citrate and uranyl acetate for observation under an electron microscope (HITACHI, HT-7800).

### Immuno-electron microscope (IEM)

Astrocytes stimulated with microglia conditioned medium for 24 h were collected for TEM samples, avoiding osmium-acid treatment. The cell clusters were cut into ultrathin 70 nm sections and placed on a nickel mesh with 200-300 mesh holes. Sections were soaked in 1% H_2_O_2_ for 10 min and then rinsed in double-distilled water for 10 min. Blocking was performed with 1% PBSA (1% BSA in PBS buffer) for 30-60 min at room temperature. 1:50 Rabbit cleaved GSDME primary antibody (Cell Signaling Technologies, E8G4U) was prepared in 1% PBSA, preincubated for 1 h at room temperature, and then incubated overnight at 4°C. The next day, sections were rinsed with PBS for 5 min, three times. Sections were then incubated with 1% PBSA for 5 min, followed by incubation with 1:40 colloidal gold-labeled secondary antibody (Abcam, ab39601) solution in 1% PBSA for 1 h at room temperature. Sections were washed three times with double-distilled water for 8 min each, stained with uranyl acetate (prepared in double-distilled water) for 12 min, and then washed with double-distilled water for 10 min. Finally, sections were stained with lead citrate for 6 min and washed with double-distilled water for 10 min. Residual water around the sections was carefully drained using filter paper, and images were taken using an electron microscope.

### RT-qPCR

Total RNA of both cell and exosomes (collected from 4×10^7^ cells) was extracted with Trizol (Invitrogen) using a standard protocol. The synthesis of cDNA was accomplished using either the TransScript® One-Step gDNA Removal and cDNA Synthesis SuperMix (TransGen Biotech) for standard RNA reverse transcription, or the miRNA 1st Strand cDNA Synthesis Kit (Vazyme Biotech) for the reverse transcription of microRNA. Quantitative RT-PCR was run using 1 μL cDNA and SYBR green chemistry (Monad Biotech) using the supplier’s protocol and a cycling program of 2 min at 95°C followed by 40 cycles of 95°C for 3 s and 60°C for 30 s on QuantStudio 3 Real-Time PCR Systems (Applied Biosystems). The primer sequences were shown in supplementary table 2. Either GAPDH or U6 snRNA was served as internal controls. The 2^−ΔΔ*C*t^ method was used for the calculation of relative fold expression of target genes.

### Western blotting

Protein samples were collected in PBS at 4°C and lysed using RIPA buffer supplemented with 1 mM phenylmethylsulfonyl fluoride (PMSF). The total protein concentration was determined using the bicinchoninic acid (BCA) assay, and equal amounts of total protein were loaded onto 12.5% SDS-polyacrylamide gel electrophoresis (PAGE) gels from Epizyme Biotech. After electrophoresis at 100 V for 60 min, the proteins were transferred to polyvinylidene difluoride (PVDF) membranes from Millipore. The membranes were blocked with 5% milk for 2 h at room temperature, followed by overnight incubation at 4°C with primary antibodies in the blocking buffer. Subsequently, the blots were washed three times with phosphate-buffered saline with Tween (PBST) and then incubated with horseradish peroxidase (HRP)-conjugated secondary antibodies for 2 h at room temperature. The protein bands were visualized using the Omni-ECL Femto Light Chemiluminescence Kit, and imaging was performed with a Chemi-Image System from Tanon.

### Immunostaining

Mice were anesthetized using 50 mg/kg Pentobarbital sodium and subsequently perfused transcardially with PBS following behavioural testing. The left hemisphere of the brain was excised and preserved in 4% paraformaldehyde at 4°C for a duration of 48 h, prior to dehydration with 20% and 30% sucrose solutions. The brains were then sectioned coronally at 30 μm using a frozen section machine (Leica Camera, Inc.). The sections underwent an overnight incubation with primary anti-mouse antibodies. Fluorescent-conjugated secondary antibodies were used to reveal the staining. The sections were then imaged using a confocal microscope (Zeiss LSM800).

For primary cell staining, 1.5×10^4^ astrocytes or 1×10^4^ neurons were cultured on poly-D-lysine coated glass slides in each well of a 24-well plate, using serum-free base medium for 24 h. The slides were then fixed in 4% paraformaldehyde for 30 min at room temperature, followed by permeabilization with PBS containing 0.1% Triton X-100 for another 30 min at room temperature. Subsequently, the cells were blocked with 3% BSA (PBS) mixture for 2 h at room temperature. An overnight incubation at 4°C was then conducted using primary anti-mouse antibodies. Fluorescein-conjugated secondary antibodies were applied to the slides for 1 h in light-protected conditions at room temperature. Washing with PB (0.1M; Na_2_HPO_4_ : NaH_2_PO_4_ = 1:1, mol/mol) was performed three times between each step. Images were captured using a fluorescence microscope after a 5 min incubation with DAPI at room temperature.

### Characterization of subcellular Ca^2+^

To obtain Ca^2+^ relative concentration of interested subcellular area, we constructed lentiviral expression vectors expressing genetically encoded Ca^2+^ indicators (GECIs). The plasmid pLVX-puro_G-CEPIA1er was purchased from HonorGene (Plasmid #164590). Plasmids for cytoplasm and mitochondria Ca^2+^ detection were obtained by replacing G-CEPIA1er with CEPIA4mt of pCMV CEPIA4mt (FENGHUISHENGWU, Plasmid #58220) and GCaMP6s of pGP-CMV-GCaMP6s (FENGHUISHENGWU, Plasmid #40753) respectively.^48,49^ The functionality of these indicators was validated by capturing subcellular Ca^2+^ dynamics using a confocal microscope (IXplore SpinSR). This allowed for the characterization of real-time fluorescence intensity changes by Image J observed 10 min post the application of 50 µM ATP (NEB, Cat# P0756S) or PBS. To explore astrocyte subcellular Ca^2+^ dynamics induced by aMCM or MCM, the real-time fluorescence intensity changes were recorded for 8 h after the stimulation using a High-content Imaging Analysis System (ImageXpress Micro Confocal).

### Saturated lipid analysis

Lipids were extracted from astrocytic exosomes according to the modified Bligh-Dyer method. Briefly, the same quantity of exosomes (80-100 μg) from indicated groups added with a deuterated lipid internal standard (Stearic acid 18,18,18-D3, #61163-39-7, Macklin Inc.) were incubated at 4°C and 1500 rpm for 1 h in a chloroform extract with 1% butylated hydroxytoluene (BHT) in methanol (1:2, v/v). Following the incubation, 350 µL of ice-cold Milli-Q water and 250 µL of ice-cold chloroform was added to induce phase separation. The mixture was then centrifuged at 16, 260*g* for 5 min at 4°C to transfer the lower organic phase containing lipids and oxidized sterols to a new tube. Subsequently, another 450 µL of ice-cold chloroform was added to the remaining aqueous phase for a second extraction. The extracts were combined and dried under organic mode in a SpeedVac. The lipid extract was re-suspended in 200 µL of chloroform:methanol (1:1, v/v). The exosomes lipid extract was utilized for targeted lipidomics analysis by a high-performance liquid chromatography-mass spectrometry (LC-MS, Thermo Scientific™ Q Exactive Plus™).

### Intracerebroventricular viral delivery in neonatal mice

Neonatal mice (postnatal days 0–3, P0–P3) were anesthetized by hypothermia on a pre-chilled ice pad for 2–3 min until the loss of reflex response. The pup’s head was disinfected with 70% ethanol and stabilized in a lateral recumbent position under a stereotaxic apparatus (Model details, Manufacturer). The skull was rotated to align the target injection site (lateral ventricle: anteroposterior −0.5 mm, mediolateral ±1.0 mm relative to bregma) upward, and the coordinates were microscopically verified (Microscope model, Manufacturer). A 34-gauge microinjection needle (Hamilton, USA) preloaded with 10 µL of AAV suspension (1×10¹² vg/mL) was vertically inserted into the lateral ventricle at a depth of 1.35 mm. Viral solution (1-2 µL) was infused at a rate of 0.2 µL/min using a microprocessor-controlled pump (Model details, Manufacturer). The needle was retained in situ for 2 min post-injection to prevent backflow and gradually withdrawn, followed by gentle pressure application with a sterile cotton swab to seal the puncture site. After allowing 5 min for wound closure, the contralateral hemisphere was injected using identical parameters. Pups were transferred to a heating pad (37°C) for 30 min to restore normothermia, monitored for respiratory stability and motor activity, and returned to the dam’s nest. Postoperative survival and maternal care were assessed for 24 h prior to subsequent experiments.

AAV viral vectors were generated by PackGene Biotech (Guangzhou, China). Briefly, HEK293 cells were transiently transfected with a GOI plasmid (pAAV-mir30shRNA.WPRE.SV40p), an adenovirus helper plasmid (pHelper), and an AAV RC plasmid (pAAV-php.eb). 72 h post-transfection, recombinant AAV were collected by lysing the cells to release the virus particles into the supernatant. The crude viral lysate was then purified via iodixanol-gradient centrifugation. Viral titers, reported in genome copies per milliliter (gc/mL), were measured using real-time PCR with Universal SYBR Green Mix (BIO-RAD, USA) and primers designed to target the inverted terminal repeat (ITR) sequence of the AAV vector. Vectors that met quality control standards were aliquoted and stored at -80°C until further use.

### Behavior test

#### Open field test

The open-field arena consisted of a white polypropylene chamber (50 × 50 × 40 cm). The central zone was defined as a 20 × 20 cm square area in the center of the arena, with the remaining area designated as the periphery. Illumination was adjusted to 60 lux using a calibrated light meter. Behavioral sessions were recorded and analyzed using Noldus Ethovision XT 11.5 software (Noldus Information Technology, Netherlands). Mice were transported to the behavioral testing room in covered cages (to minimize visual stress) and allowed to acclimate to the environment for 1 hour prior to testing. At the start of each trial, mice were gently grasped by the base of the tail and positioned head-first toward the corner of the arena, ensuring abdominal contact with the wall. Mice were allowed to freely explore the arena for 10 min while being video-recorded. After the trial, mice were returned to their home cages, and individual animals were identified by tail markings corresponding to their assigned arena and video codes. The arena and surrounding surfaces were thoroughly cleaned with 70% ethanol between trials to eliminate residual odors. The tracking software automatically quantified two parameters. First, center zone exploration, defined as the percentage of time spent in the central zone (20 × 20 cm). Second, total distance traveled (cm), representing cumulative locomotor activity across the entire arena.

#### Y-Maze

The Y-maze comprised three identical opaque acrylic arms (length 50 cm × width 18 cm × height 35 cm) arranged at 120° angles. Distinct geometric patterns (triangle, square, and circle) were affixed to each arm’s distal end as visual-spatial cues. Ambient illumination was maintained at 40 lux, verified using a calibrated light meter. Behavioral sessions were recorded and analyzed with Noldus Ethovision XT 11.5 software (Noldus Information Technology, Netherlands). Mice were transported to the behavioral room in covered cages and acclimated for 1 h. Each mouse was placed at the distal end of a randomly selected arm, facing away from the center, and allowed 8 minutes of free exploration. An arm entry required all four paws to enter an arm. Three parameters were quantified: total arm entries (complete entries during the session), spontaneous alternation (sequential entries into three distinct arms without repetition), and alternation score (calculated as [number of alternations / maximum alternations] × 100%).

#### Novel object recognition test

The test was conducted in the same open-field arena (50 cm × 50 cm × 40 cm) used for baseline behavioral assessments. Two identical objects (e.g., cylindrical batteries; 5 cm height × 2 cm diameter) were placed symmetrically in the arena during training phases. A novel object (distinct in shape/texture but matched for size) replaced one familiar object in the final test phase. Behavioral data were recorded using Noldus Ethovision XT 11.5 software (Noldus Information Technology, Netherlands).

Prior to testing, mice were transported to the behavioral testing room in covered cages and allowed to acclimate for 1 hour. Twenty-four hours before formal testing, each mouse was placed in the empty arena for a 5-min habituation session to reduce environmental novelty. After 24 hours, two identical objects were symmetrically positioned 15 cm from the arena walls, and mice were allowed to freely explore the objects for 10 min. Twelve hours later, the same two objects were reintroduced, and exploration behavior was recorded for 10 min to reinforce object familiarity. Six hours after the second training session, one familiar object was replaced with a novel object (position counterbalanced across subjects), and mice were tested for 10 min. Between trials, the arena and objects were cleaned with 70% ethanol to eliminate residual odors. Object exploration was operationally defined as direct sniffing or tactile contact with the object, measured when the snout was within 2 cm of it. Novel object preference index (PI) was calculated according to two distinct metrics. Specifically, time-based PI represented the ratio of time spent exploring the novel object to total exploration time (both objects), expressed as a percentage [(Novel object exploration time / Total exploration time) × 100%]. Frequency-based PI reflected the proportion of contacts made with the novel object relative to total contacts with both objects, calculated as [(Number of novel object contacts / Total contacts) × 100%].

#### siRNA transfection

Primary astrocytes were transfected with siRNA using Lipofectamine LTX&PLUS Reagent (Thermo Fisher Scientific, Cat#15338100) following the manufacturer’s protocol. Briefly, 1 hour prior to transfection, the culture medium was replaced with Opti-MEM Reduced-Serum Medium (Gibco, Cat#31985062) to minimize serum interference. For each well of a 24-well plate, a transfection complex was prepared by combining 50 nM siRNA (targeting *GSDME* or *Camk2a*; scrambled siRNA served as a negative control), 1 μL PLUS Reagent, and 1 μL Lipofectamine LTX Reagent in 50 μL Opti-MEM. The mixture was incubated at room temperature for 5 minutes to allow complex formation and then added dropwise to the cells.

#### Lentivirus preparation and infection

Lentiviral stocks were generated by co-transfecting HEK293T cells with the target plasmid (pLV-GSDME-NTD-G4S-eGFP, pLV-EMC10-GSDME-NTD-G4S-eGFP, pLV-EMC10-GSDME-NTD(Δ56)-G4S-eGFP, pLV-Emc10-eGFP, pLKO.1-shRNA-Scramble and pLKO.1-shRNA-Gsdme) and helper plasmids pCMV-Δ8.91, and pCMV-VSV-G using CarpTrans (OPM, Cat#AC501302) as recommended by the manufacturer. After 48 h post-transfection, lentivirus supernatant was filtered through 0.45 μm filter (Millipore) and concentrated using PEG8000 (Beyotime, Cat#ST43). The final lentivirus stocks were resuspended in DMEM (OPM, Cat#P081702-001) and stored at -80°C. For transfection, primary mouse astrocytes were incubated with lentivirus overnight and then replaced with fresh complete media. Cells were used for the assays at 120 h post transfection.

#### Peripheral effect of ADVs *in vivo*

Serum-derived ADVs were isolated from 200 μL of blood samples obtained from both AD patients and healthy controls. The concentrated ADVs (1 mg/mL, 100 μL per mice) were administered via tail vein injection to 8-week-old C57/BL6 mice. Whole blood samples (30 μL) were collected at 1-hour and 2-hour post-injection into 200 μL of EDTA anticoagulant buffer (0.25 mg/mL). Platelet and neutrophil counts were subsequently quantified using an automated hematology analyzer (XP-100; Sysmex, Kobe, Japan).

## Supplementary Figures

**Figure S1.**
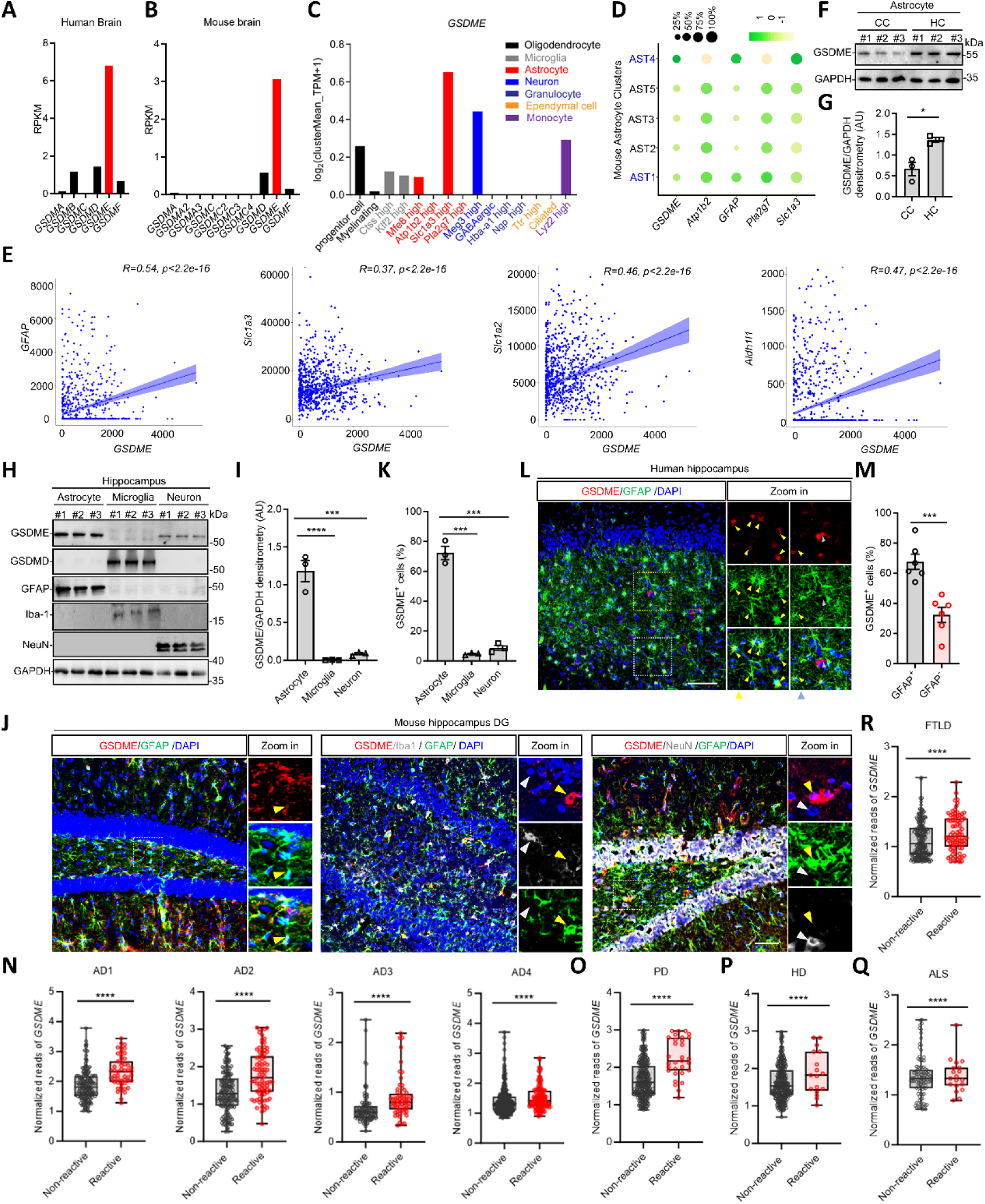
Spatiotemporal and Disease-Associated Heterogeneity of GSDME Expression in Astrocytes, related to. **Figure 1**. (A) Protein expression overview of gasdermin (GSDM) A-F in human brain according to the HPA RNA-seq normal tissues.^30^ (B) Protein expression overview of GSDMs in mouse brain according to the Mouse ENCODE transcriptome data.^31^ (C) Single-cell mRNA transcripts of GSDME in different cell subtypes of whole mouse brain were shown according to the published single-cell database (http://bis.zju.edu.cn/MCA/). (D) Spatial distribution of GSDME in different astrocyte subtypes was analyzed by open single-cell database.^33^ (E) Correlation between *GSDME* and *GFAP*, *Slc1a3*, *Slc1a2* and *Adh1l1* is calculated as Spearman correlation based on the single-cell RNA-seq expression data.^33^ (F and G) Immunoblot analysis of GSDME expression in primary astrocytes isolated from cerebral cortex (CC) or hippocampus (HC) of P1-3 mouse pups measured by ImageJ based on gray scale (*n* = 3 replicates per group). (H and I) Immunoblot analysis of GSDME expression in primary astrocytes (GFAP^+^), microglia (Iba1^+^) and neurons (NeuN^+^) isolated from hippocampus DG region of P1-3 mouse pups measured by ImageJ based on gray scale (*n* = 3 mice per group). (J and K) Representative IF images of mouse hippocampus DG region co-stained with GSDME, GFAP and either Iba1 or NeuN antibodies (J). Enlarged images of the boxed area are shown. For colocalization, each dot represents two coronal sections of one mouse (K). Scale bars, 40 μm. (L and M) Representative immunofluorescence (IF) images of hippocampus region from three human donors co-stained with anti-GSDME antibody and anti-GFAP antibody. The yellow arrows indicate GFAP^+^GSDME^+^ cells, while the blue arrows indicate GFAP^-^GSDME^+^ cells. Scale bars, 100 μm. Percentage of GSDME-positive cells within GFAP^+^ or GFAP*^-^* cell populations were quantified using Image J software. (N-R) Comparison of GSDME mRNA levels between non-reactive (NR) and reactive (R) astrocytes in snRNA-seq datasets of (N), AD (AD1^34^, *n*_NR_ = 165, *n*_R_ = 51; AD2^35^, *n*_NR_ = 177, *n*_R_ = 82; AD3^36^, *n*_NR_ = 103, *n*_R_ = 61; AD4^38^, *n*_NR_ = 367, *n*_R_ = 128) (D), PD^40^ (*n*_NR_ = 367, *n*_R_ = 31) (E), HD^39^ (*n*_NR_ = 367, *n*_R_ = 17) (F), ALS^39^ (*n*_NR_ = 106, *n*_R_ = 19) (G), FTLD^39^ (*n*_NR_ = 82, *n*_R_ = 253). Significance levels are determined using a two-sided Wilcoxon signed-rank test. All values are represented as mean ± SEM, \*\*\**p* < 0.001.

**Figure S2.**
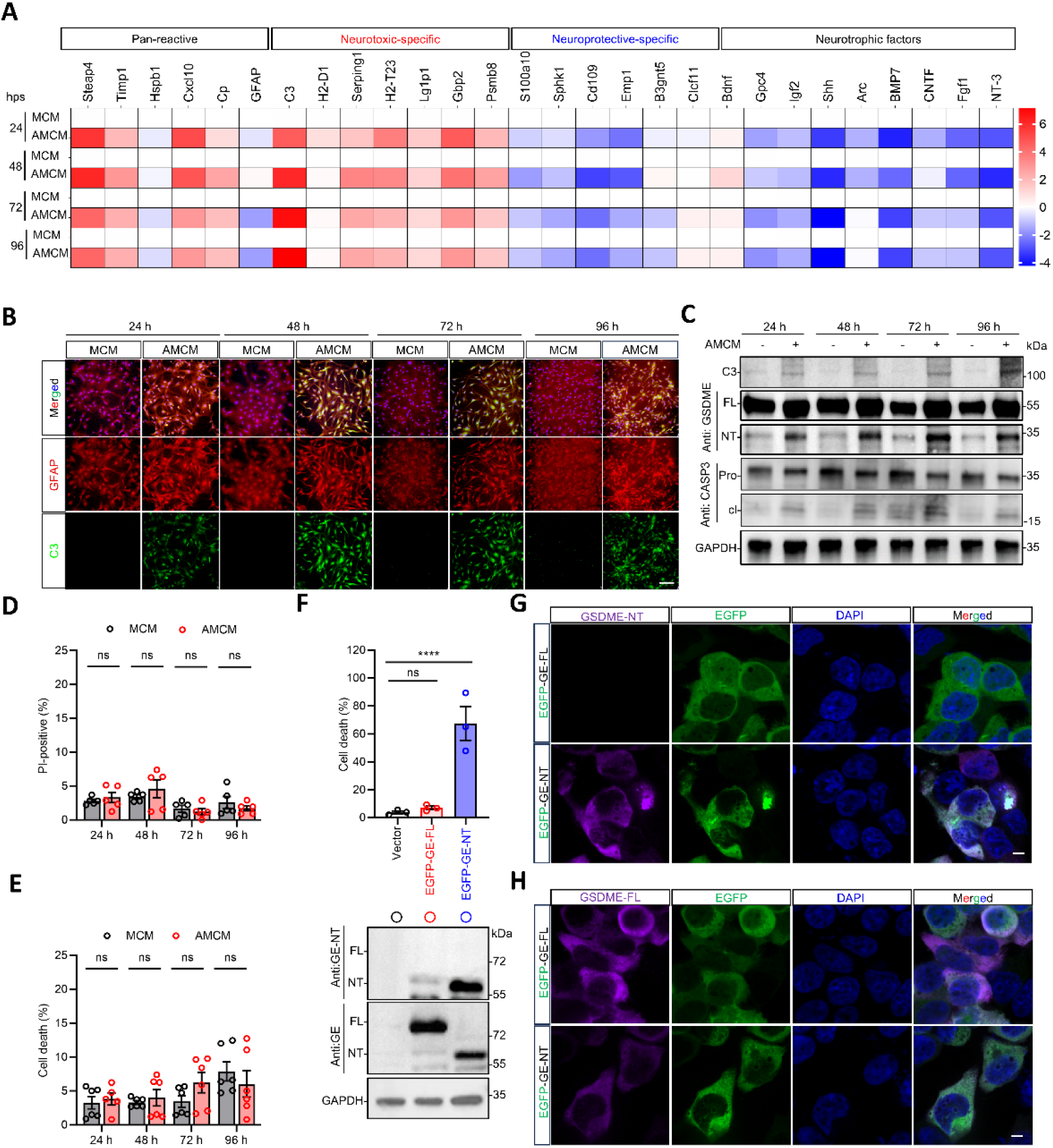
AMCM induces the neurotoxic phenotype and GSDME activation in primary astrocytes, related to Figure 2. (A) The mRNA expression of neurotoxic, neuroprotective and neurotrophic factors in astrocytes treated with MCM or AMCM (*n* = 3 replicates per group). (B) IF analysis of the induction of astrocytic reactivity by co-staining with C3 (Anti-C3, green) and GFAP (Anti-GFAP, red) at 24, 48, 72, 96 hps. Scale bars, 100 μm. (C) Quantification of the activation of caspase-3, GSDME in primary astrocytes as well as the secretion of C3 in the medium at 24, 48, 72, 96 hps. (D and E) Quantification of primary astrocytic cell death by PI (Prodium Iodide) staining (D) or LDH (lactate dehydrogenase) release assay (E) at 24, 48, 72, 96 hps. (F) Overexpression of EGFP-tagged GSDME (EGFP-GSDME-FL) or EGFP tagged GSDME-NT (EGFP-GSDME-NT) in HEK293T cells, and the cell death was determined by LDH release assay. The expression levels of EGFP-GSDME-FL, NT were further confirmed by immunoblotting using antibodies for either GSDME-FL or GSDME-NT special. (G and H) IF analysis of HEK293T cells in (F) with antibodies for either GSDME-NT special (G) or GSDME-FL (H) to validate the specificity of the antibody for GSDME NT. Scale bars, 10 μm. Statistical significance determined using a two-tailed Student’ s *t* test (D-E, *n* = 3 replicates). All values are represented as mean ± SEM, \**p* < 0.05; \*\**p* < 0.01; \*\*\**p* <0.001; \*\*\*\**p* < 0.0001.

**Figure S3.**
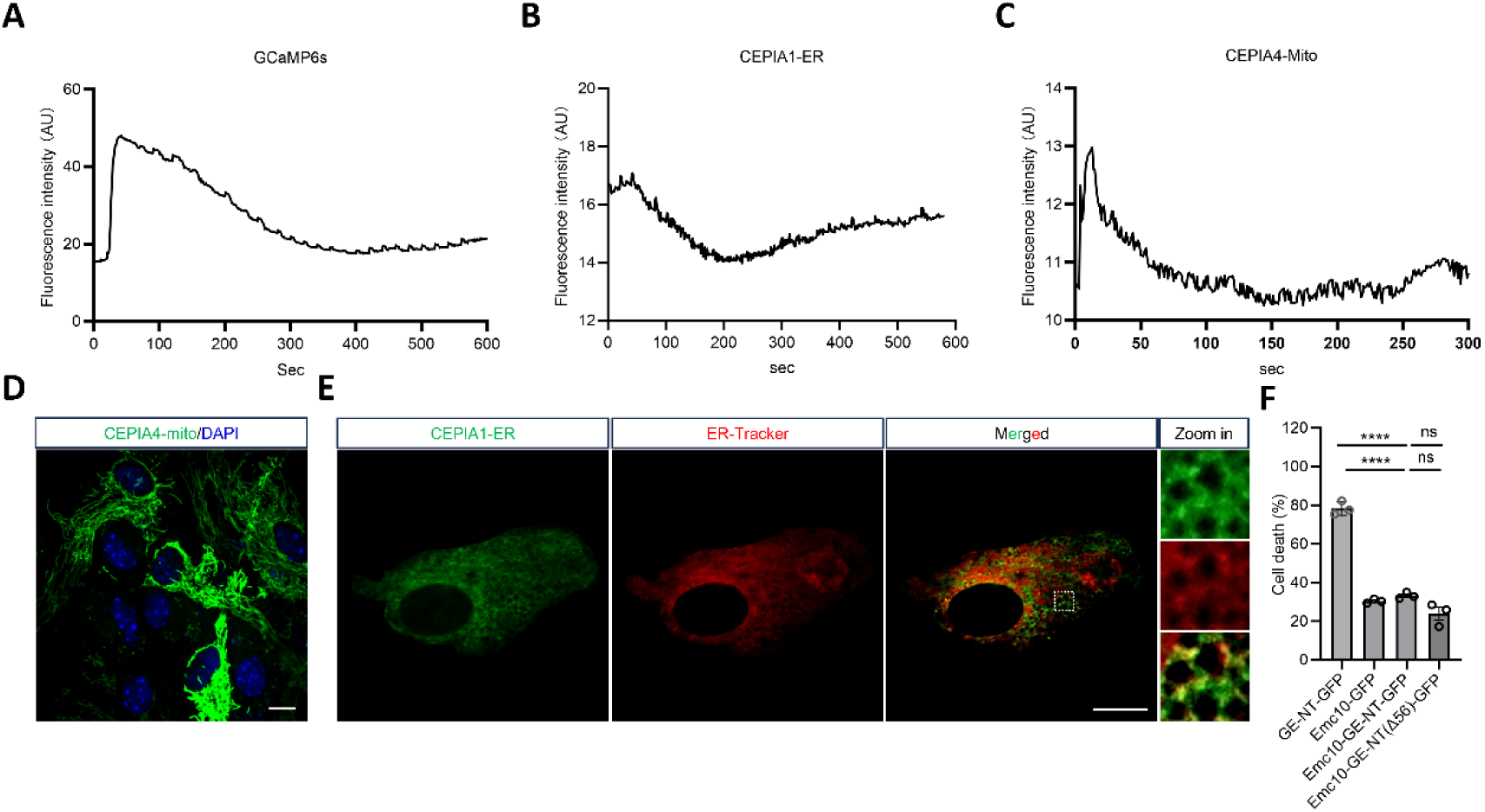
Detection of Ca^2+^ dynamics, related to Figure 3. (A-C) Quantifying of relative fluorescence intensity in HEK293T cells treated with 100 mM ATP or PBS within 10 min using the genetically encoded GCaMP6s for the cytosol (A), CEPIA1-ER for the ER (B), or CEPIA4-Mito for the mitochondria (C), to verify the response of these indicators. A representative curve is plotted based on data obtained from at least three independent experiments. (D) The location of CEPIA4-Mito was confirmed by IF staining with DAPI (blue). (E) The location of and CEPIA1-ER was confirmed by IF staining with ER-tracker (Red). Enlarged images of the boxed area are shown. Scale bars, 5 μm. (F) Quantification of primary astrocytic cell death by LDH (lactate dehydrogenase) release assay.

**Figure S4.**
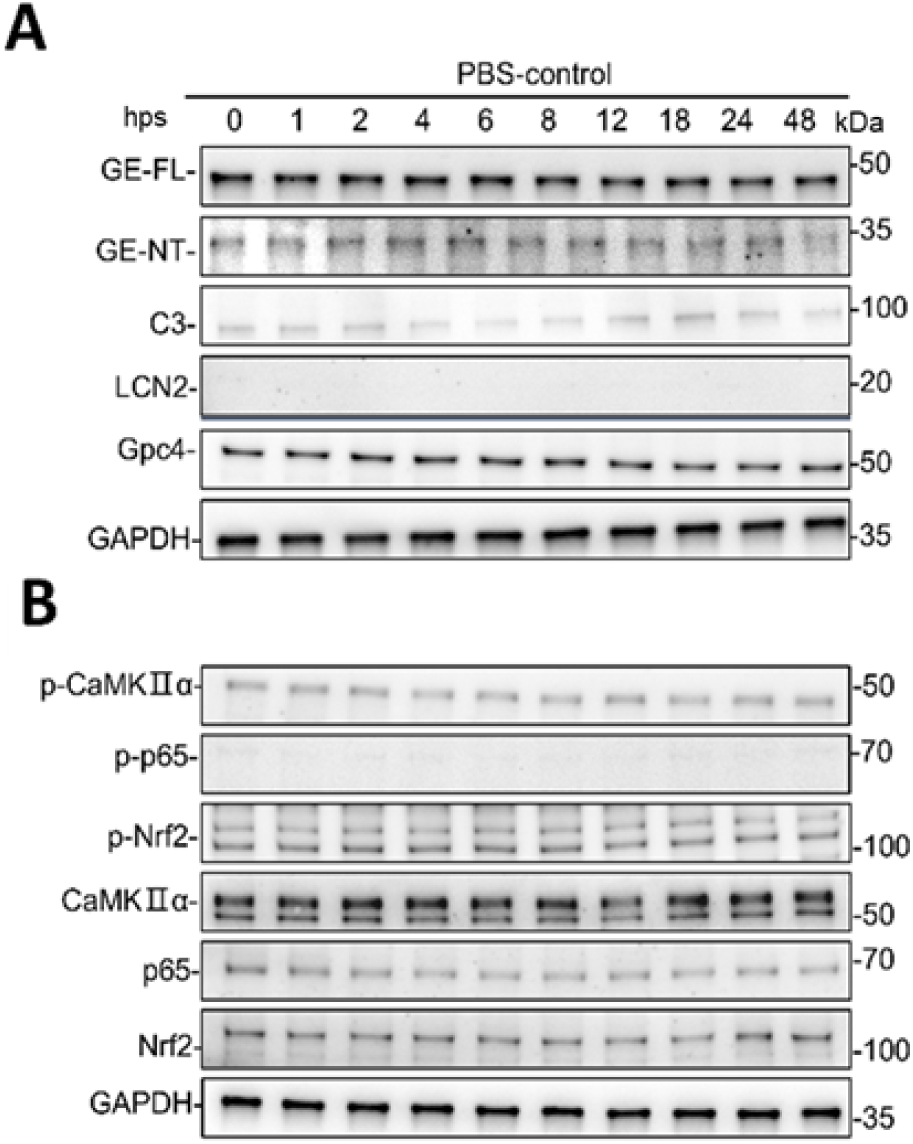
Temporal dynamics of GSDME cleavage and signaling pathways in PBS-treated astrocytes, related to Figure 4. (A) Time-dependent immunoblot analysis of GSDME full-length (GSDME-FL) and its N-terminal cleavage fragment (GSDME-NT), neurotoxic markers (C3, LCN2), and the neurotrophic factor GPC4 in astrocytes transfected with non-targeting scrambled siRNA (siNC) following the administration of PBS. (B) Phosphorylation kinetics of CaMKIIα, p65, and NRF2 in (A).

**Figure S5.**
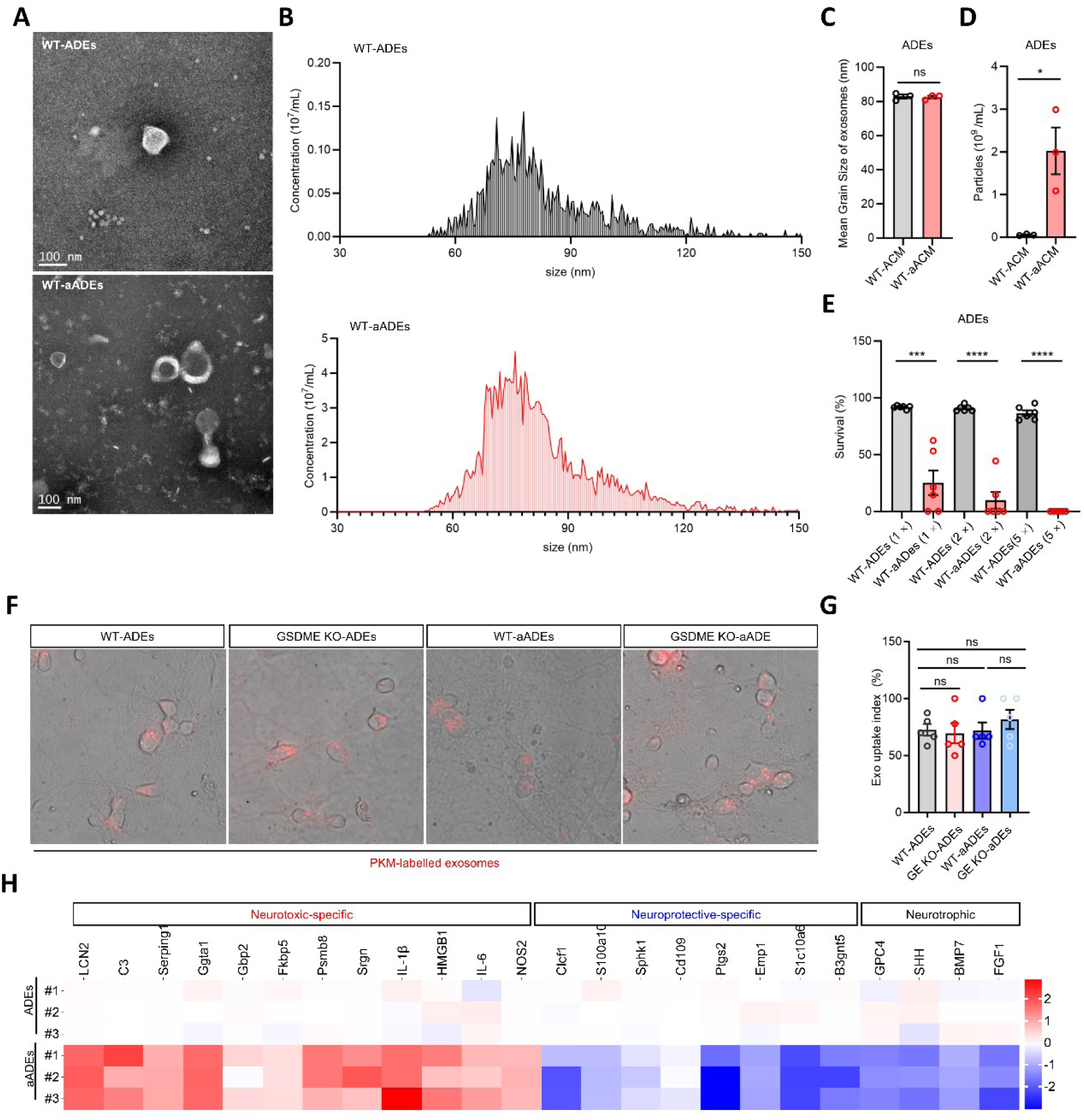
Uptake of astrocytic exosomes by neurons, related to Figure 5. (A) SEM images of wild type (WT) or reactive exosomes purified from WT-ACM or WT-aACM astrocytes, respectively. Scar bars, 100 nm. (B-D) Quantifying of the particle size of either WT or reactive exosomes by nanoparticle tracking analysis (B). The mean grain size (C) and number of exosome particles (D) were calculated (*n*= 3 replicates per group). (E) Quantification of the neuronal cell survival rate of mouse cortical neurons 24 h after the treatment with differently concentrated exosomes purified from equal volume of ACM or aACM, respectively. 1 × means that the isolated exomes was resuspended with equal volume of neuronal culture medium compared to that of ACM (*n*= 6 replicates per group). (F and G) Representative images of primary cortical neurons phagocyting PKM-labelled exosomes isolated from ACM of WT-ACM, GSDME KO-ACM, WT-aACM and GSDME KO-aACM astrocytes. Scale bars, 20 μm. The ratio of exosome (Exo) uptake by neurons was calculated by Image J (*n*= 5 replicates per group). (H) Heatmap depicting relative expression of astrocyte state genes in ADEs by transcriptional sequencing (*n* = 3 replicates per group). Statistical significance determined using a two-tailed Student’s *t* test. All values are represented as mean ± SEM, \**p* < 0.05; \*\**p* < 0.01; \*\*\**p* < 0.001; \*\*\*\**p* < 0.0001.

**Figure S6.**
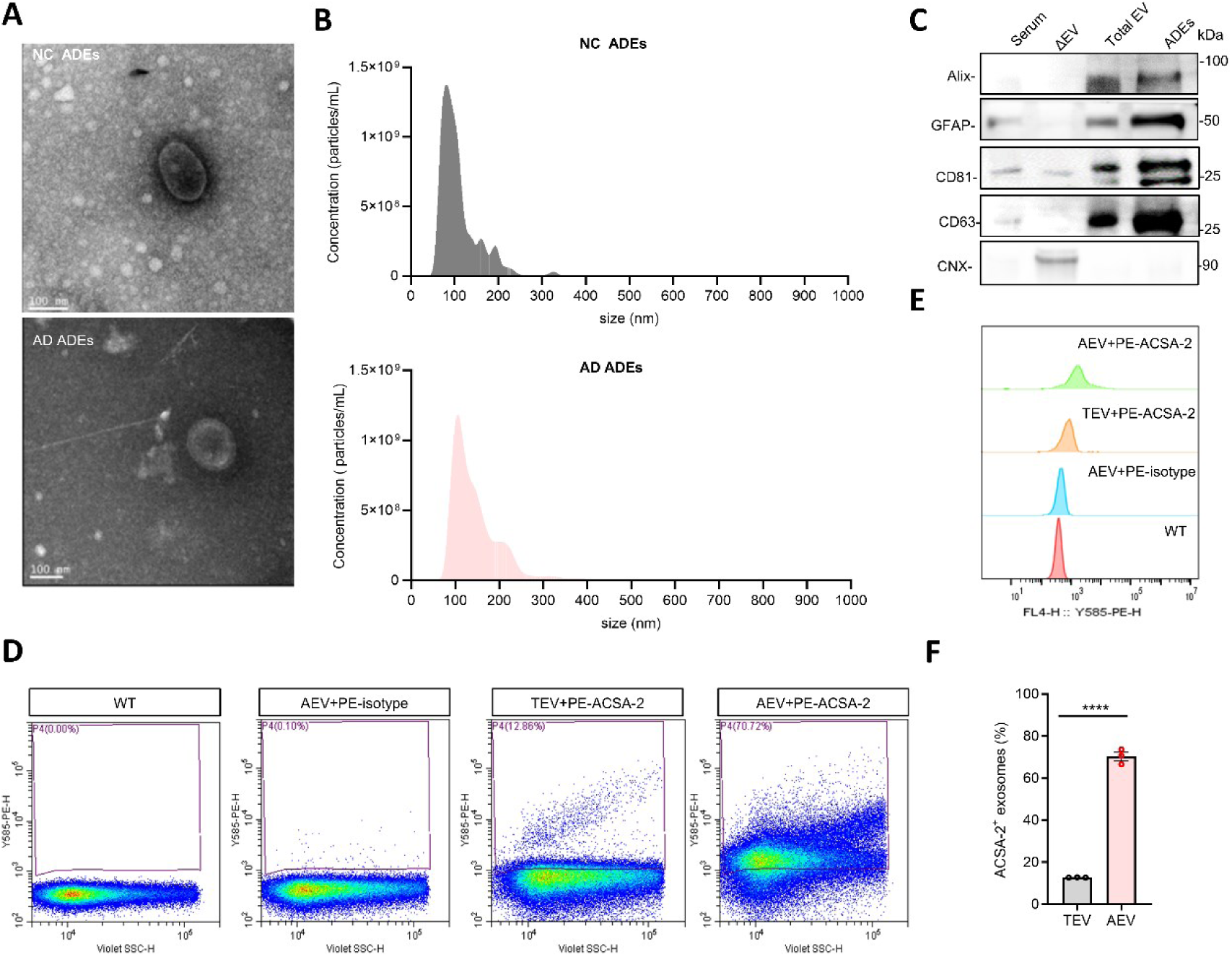
Isolation and identification of serum-derived astrocytic exosomes, related to Figure 7 and Figure 8. (A) Characterization of serum astrocyte-derived exosomes (ADEs) from healthy individuals and AD patients by TEM. scale bar, 100 nm. (B) NTA of ADEs isolated from human serum, showing size distribution and particle concentration. (C) Western blot analysis of protein markers in human serum ADEs. Compared to whole serum, ADEs exhibit robust expression of extracellular vesicle (EV)-specific markers (Alix, CD81, CD63) and the astrocytic marker GFAP, while lacking the endoplasmic reticulum marker calreticulin. (D-F) Representative flow cytometric analysis of ADEs. Purity validation using phycoerythrin (PE)-conjugated anti-astrocyte surface antigen-2 (ACSA-2) antibodies. ACSA-2-positive EVs account for ∼70% of ADEs, compared to ∼12% in total serum exosomes, confirming enrichment of astrocyte-specific ADEs. Statistical significance determined using a two-tailed Student’s *t* test.

**Table S1.**
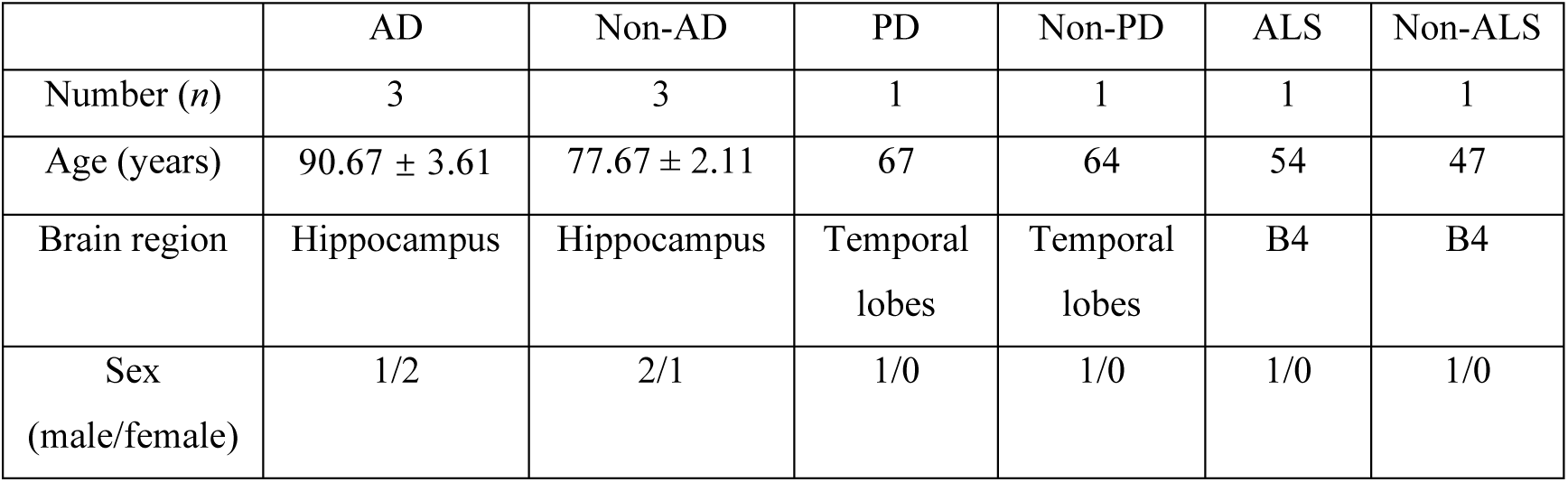
The characteristics of human donors, related to Figure 1.

|  | AD | Non-AD | PD | Non-PD | ALS | Non-ALS |
| --- | --- | --- | --- | --- | --- | --- |
| Number (*n*) | 3 | 3 | 1 | 1 | 1 | 1 |
| Age (years) | 90.67 ± 3.61 | 77.67 ± 2.11 | 67 | 64 | 54 | 47 |
| Brain region | Hippocampus | Hippocampus | Temporal lobes | Temporal lobes | B4 | B4 |
| Sex (male/female) | 1/2 | 2/1 | 1/0 | 1/0 | 1/0 | 1/0 |

**Table S2.**
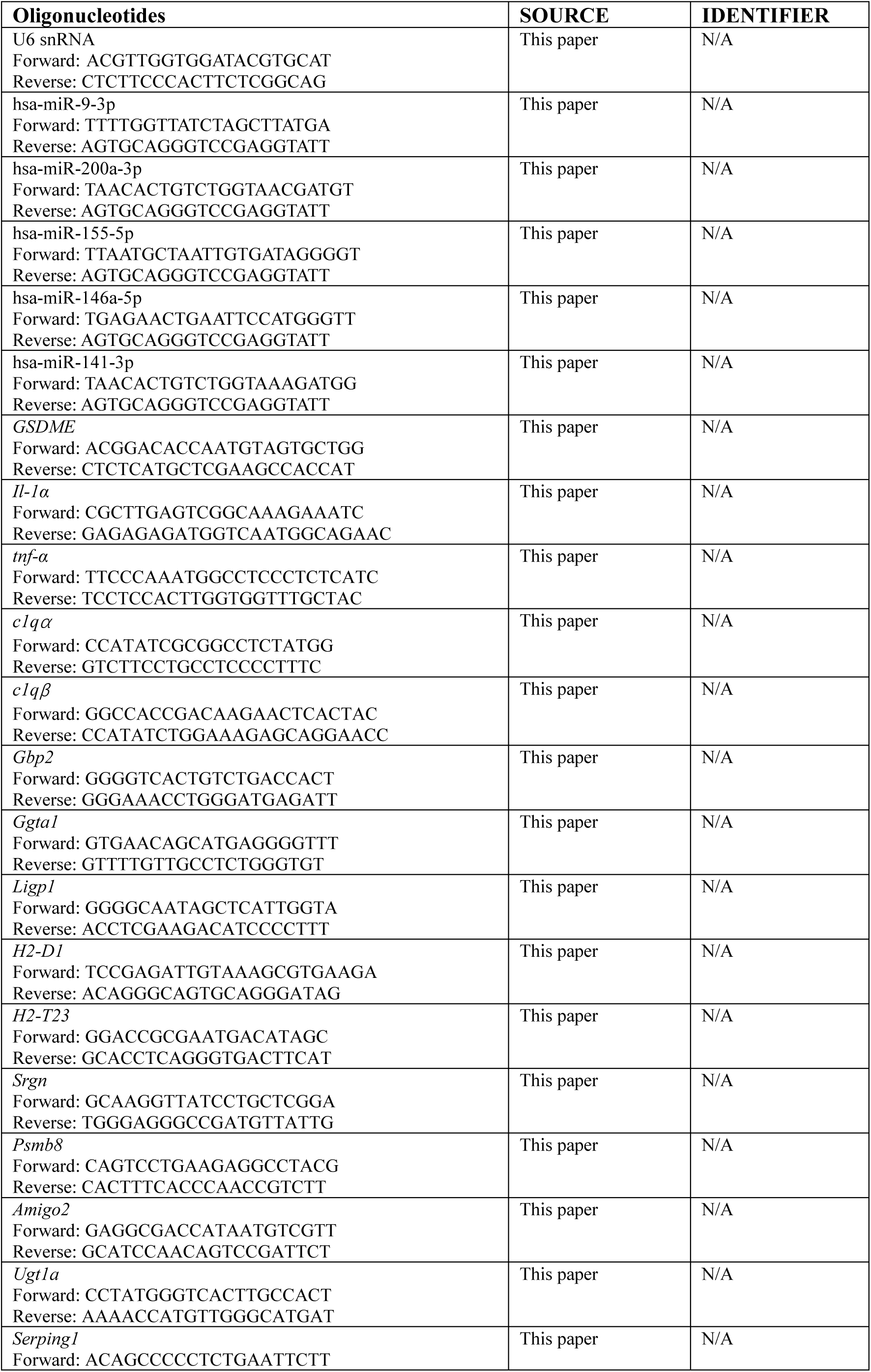
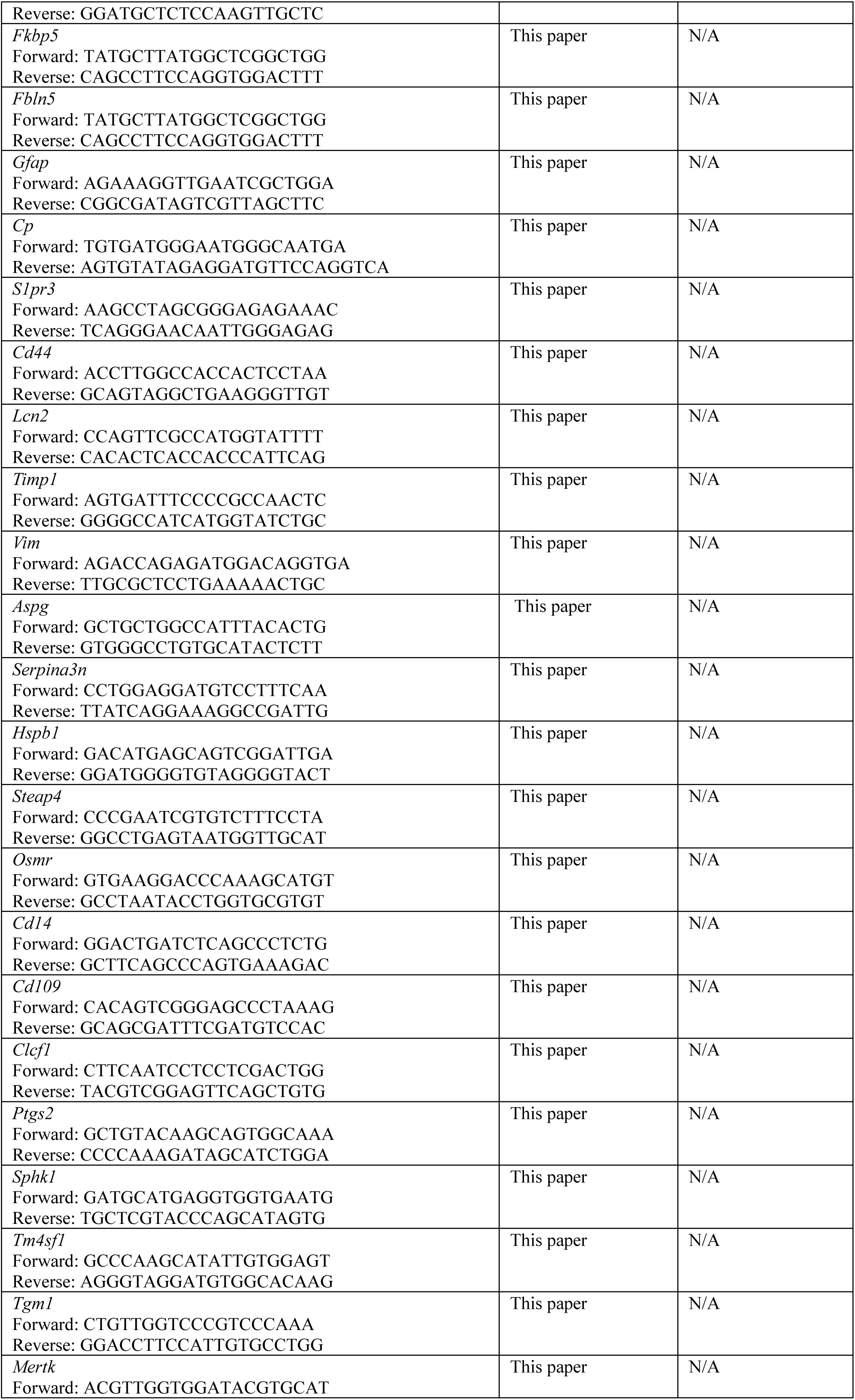
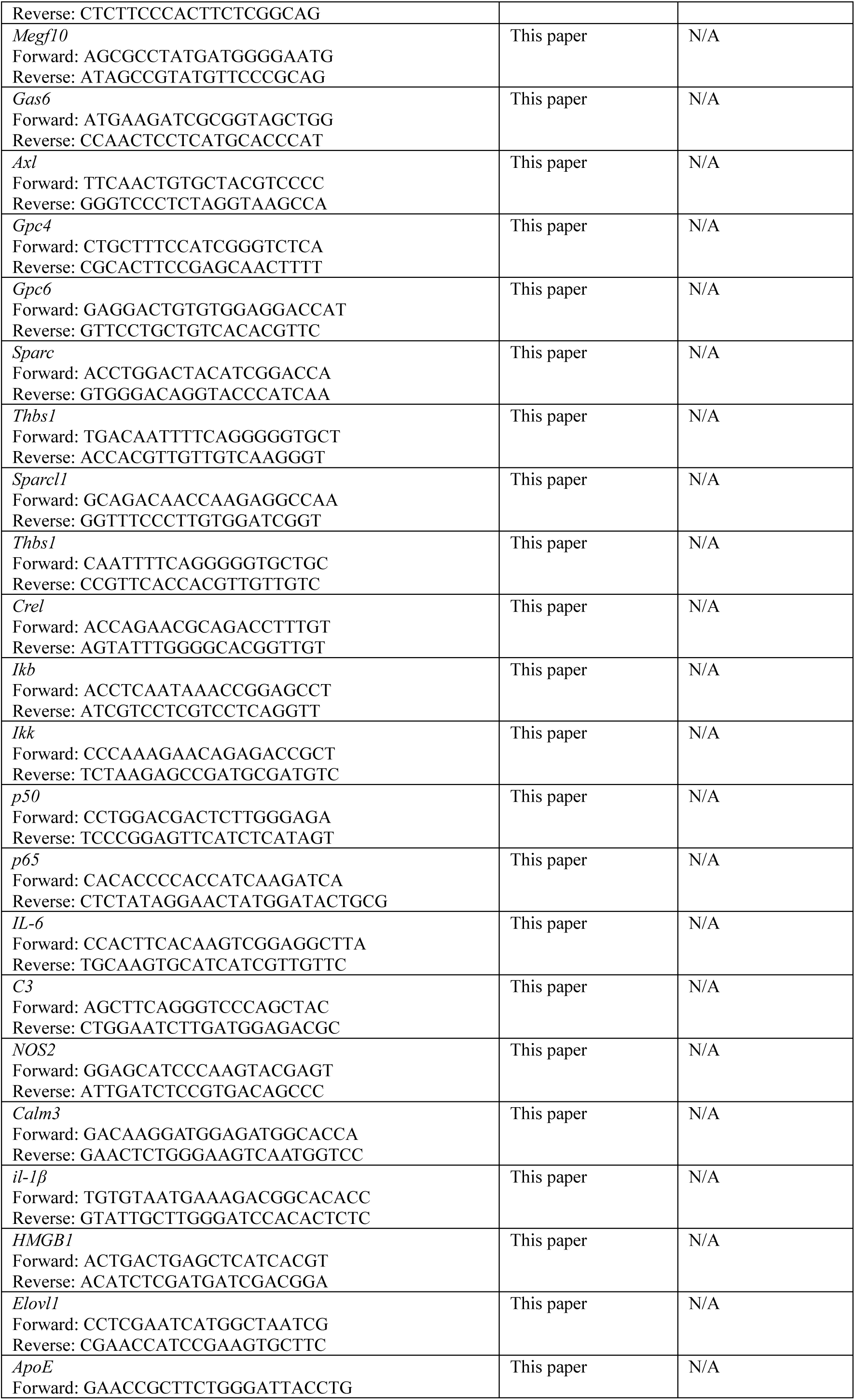
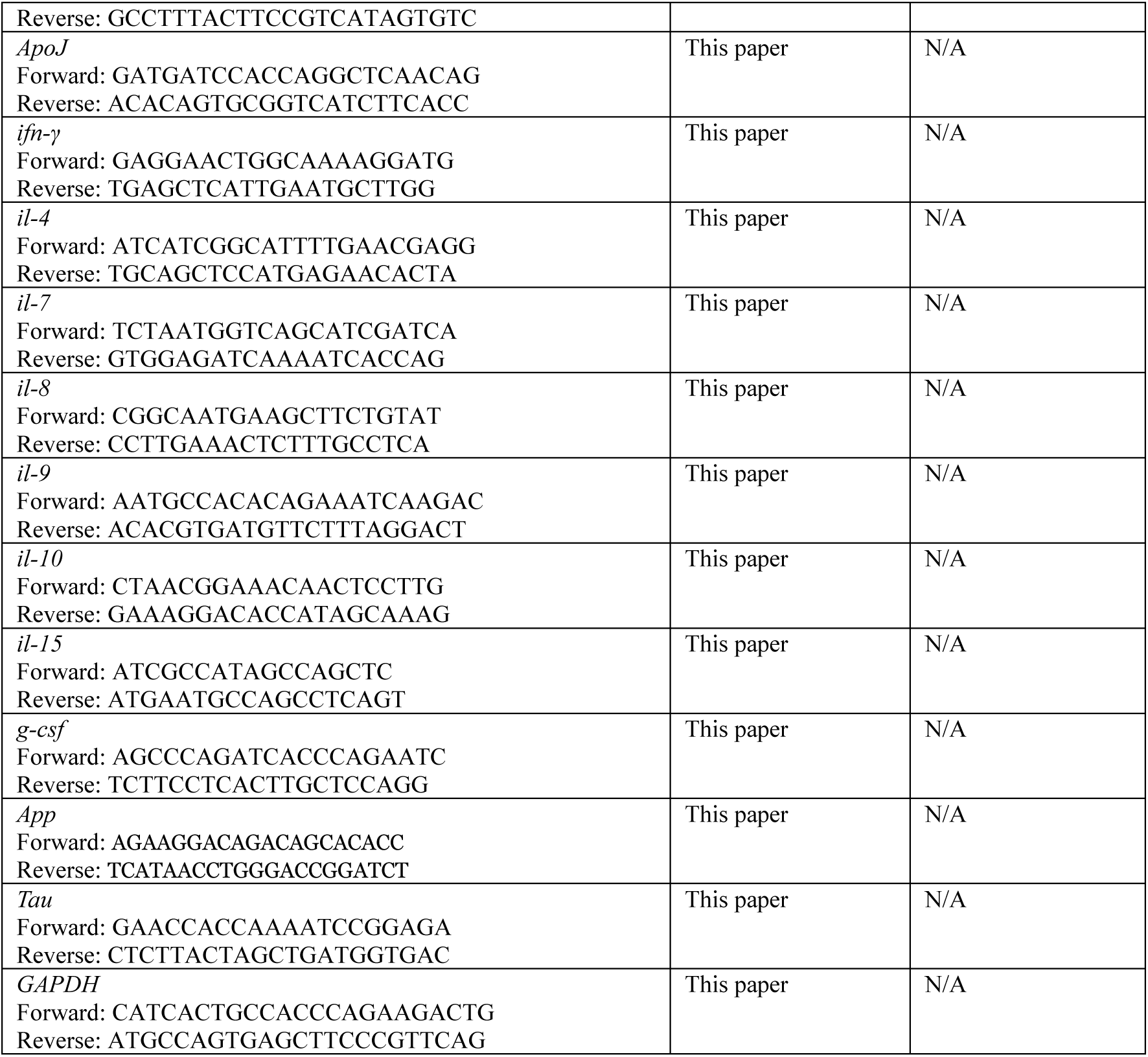
Primers used in this study.

## Notes

### Competing Interest Statement

The authors have declared no competing interest.

## REFERENCES

1. Busche, M.A., and Hyman, B.T. (2020). Synergy between amyloid-beta and tau in Alzheimer’s disease. Nat Neurosci 23, 1183–1193. 10.1038/s41593-020-0687-6.

2. Long, J.M., and Holtzman, D.M. (2019). Alzheimer Disease: An Update on Pathobiology and Treatment Strategies. Cell 179, 312–339. 10.1016/j.cell.2019.09.001.

3. Hardy, J.A., and Higgins, G.A. (1992). Alzheimer’s disease: the amyloid cascade hypothesis. Science 256, 184–185. 10.1126/science.1566067.

4. Braak, H., and Del Tredici, K. (2011). Alzheimer’s pathogenesis: is there neuron-to-neuron propagation? Acta Neuropathol 121, 589–595. 10.1007/s00401-011-0825-z.

5. Chetelat, G., La Joie, R., Villain, N., Perrotin, A., de La Sayette, V., Eustache, F., and Vandenberghe, R. (2013). Amyloid imaging in cognitively normal individuals, at-risk populations and preclinical Alzheimer’s disease. Neuroimage Clin 2, 356–365. 10.1016/j.nicl.2013.02.006.

6. Mattsson-Carlgren, N., Salvado, G., Ashton, N.J., Tideman, P., Stomrud, E., Zetterberg, H., Ossenkoppele, R., Betthauser, T.J., Cody, K.A., Jonaitis, E.M., et al. (2023). Prediction of Longitudinal Cognitive Decline in Preclinical Alzheimer Disease Using Plasma Biomarkers. JAMA Neurol 80, 360–369. 10.1001/jamaneurol.2022.5272.

7. Patani, R., Hardingham, G.E., and Liddelow, S.A. (2023). Functional roles of reactive astrocytes in neuroinflammation and neurodegeneration. Nat Rev Neurol 19, 395–409. 10.1038/s41582-023-00822-1.

8. Rothstein, J.D., Dykes-Hoberg, M., Pardo, C.A., Bristol, L.A., Jin, L., Kuncl, R.W., Kanai, Y., Hediger, M.A., Wang, Y., Schielke, J.P., and Welty, D.F. (1996). Knockout of glutamate transporters reveals a major role for astroglial transport in excitotoxicity and clearance of glutamate. Neuron 16, 675–686. 10.1016/s0896-6273(00)80086-0.

9. Bellaver, B., Povala, G., Ferreira, P.C.L., Ferrari-Souza, J.P., Leffa, D.T., Lussier, F.Z., Benedet, A.L., Ashton, N.J., Triana-Baltzer, G., Kolb, H.C., et al. (2023). Astrocyte reactivity influences amyloid-beta effects on tau pathology in preclinical Alzheimer’s disease. Nat Med 29, 1775–1781. 10.1038/s41591-023-02380-x.

10. Escartin, C., Galea, E., Lakatos, A., O’Callaghan, J.P., Petzold, G.C., Serrano-Pozo, A., Steinhauser, C., Volterra, A., Carmignoto, G., Agarwal, A., et al. (2021). Reactive astrocyte nomenclature, definitions, and future directions. Nat Neurosci 24, 312–325. 10.1038/s41593-020-00783-4.

11. Pereira, J.B., Janelidze, S., Smith, R., Mattsson-Carlgren, N., Palmqvist, S., Teunissen, C.E., Zetterberg, H., Stomrud, E., Ashton, N.J., Blennow, K., and Hansson, O. (2021). Plasma GFAP is an early marker of amyloid-beta but not tau pathology in Alzheimer’s disease. Brain 144, 3505–3516. 10.1093/brain/awab223.

12. De Strooper, B., and Karran, E. (2016). The Cellular Phase of Alzheimer’s Disease. Cell 164, 603–615. 10.1016/j.cell.2015.12.056.

13. van Dyck, C.H., Sabbagh, M., and Cohen, S. (2023). Lecanemab in Early Alzheimer’s Disease. Reply. N Engl J Med 388, 1631–1632. 10.1056/NEJMc2301380.

14. Henstridge, C.M., Hyman, B.T., and Spires-Jones, T.L. (2019). Beyond the neuron-cellular interactions early in Alzheimer disease pathogenesis. Nat Rev Neurosci 20, 94–108. 10.1038/s41583-018-0113-1.

15. Brandebura, A.N., Paumier, A., Onur, T.S., and Allen, N.J. (2023). Astrocyte contribution to dysfunction, risk and progression in neurodegenerative disorders. Nat Rev Neurosci 24, 23–39. 10.1038/s41583-022-00641-1.

16. Verkhratsky, A., and Nedergaard, M. (2018). Physiology of Astroglia. Physiol Rev 98, 239–389. 10.1152/physrev.00042.2016.

17. Shah, D., Gsell, W., Wahis, J., Luckett, E.S., Jamoulle, T., Vermaercke, B., Preman, P., Moechars, D., Hendrickx, V., Jaspers, T., et al. (2022). Astrocyte calcium dysfunction causes early network hyperactivity in Alzheimer’s disease. Cell Rep 40, 111280. 10.1016/j.celrep.2022.111280.

18. Takano, T., Tian, G.F., Peng, W., Lou, N., Libionka, W., Han, X., and Nedergaard, M. (2006). Astrocyte-mediated control of cerebral blood flow. Nat Neurosci 9, 260–267. 10.1038/nn1623.

19. Liddelow, S.A., Guttenplan, K.A., Clarke, L.E., Bennett, F.C., Bohlen, C.J., Schirmer, L., Bennett, M.L., Munch, A.E., Chung, W.S., Peterson, T.C., et al. (2017). Neurotoxic reactive astrocytes are induced by activated microglia. Nature 541, 481–487. 10.1038/nature21029.

20. Reichenbach, N., Delekate, A., Breithausen, B., Keppler, K., Poll, S., Schulte, T., Peter, J., Plescher, M., Hansen, J.N., Blank, N., et al. (2018). P2Y1 receptor blockade normalizes network dysfunction and cognition in an Alzheimer’s disease model. J Exp Med 215, 1649–1663. 10.1084/jem.20171487.

21. Hasel, P., Rose, I.V.L., Sadick, J.S., Kim, R.D., and Liddelow, S.A. (2021). Neuroinflammatory astrocyte subtypes in the mouse brain. Nat Neurosci 24, 1475–1487. 10.1038/s41593-021-00905-6.

22. Chen, W.T., Lu, A., Craessaerts, K., Pavie, B., Sala Frigerio, C., Corthout, N., Qian, X., Lalakova, J., Kuhnemund, M., Voytyuk, I., et al. (2020). Spatial Transcriptomics and In Situ Sequencing to Study Alzheimer’s Disease. Cell 182, 976–991 e919. 10.1016/j.cell.2020.06.038.

23. Zamanian, J.L., Xu, L., Foo, L.C., Nouri, N., Zhou, L., Giffard, R.G., and Barres, B.A. (2012). Genomic analysis of reactive astrogliosis. J Neurosci 32, 6391–6410. 10.1523/JNEUROSCI.6221-11.2012.

24. Habib, N., McCabe, C., Medina, S., Varshavsky, M., Kitsberg, D., Dvir-Szternfeld, R., Green, G., Dionne, D., Nguyen, L., Marshall, J.L., et al. (2020). Disease-associated astrocytes in Alzheimer’s disease and aging. Nat Neurosci 23, 701–706. 10.1038/s41593-020-0624-8.

25. Rothhammer, V., Borucki, D.M., Tjon, E.C., Takenaka, M.C., Chao, C.C., Ardura-Fabregat, A., de Lima, K.A., Gutierrez-Vazquez, C., Hewson, P., Staszewski, O., et al. (2018). Microglial control of astrocytes in response to microbial metabolites. Nature 557, 724–728. 10.1038/s41586-018-0119-x.

26. Wheeler, M.A., Clark, I.C., Tjon, E.C., Li, Z., Zandee, S.E.J., Couturier, C.P., Watson, B.R., Scalisi, G., Alkwai, S., Rothhammer, V., et al. (2020). MAFG-driven astrocytes promote CNS inflammation. Nature 578, 593–599. 10.1038/s41586-020-1999-0.

27. Fu, J., Schroder, K., and Wu, H. (2024). Mechanistic insights from inflammasome structures. Nat Rev Immunol 24, 518–535. 10.1038/s41577-024-00995-w.

28. Neel, D.V., Basu, H., Gunner, G., Bergstresser, M.D., Giadone, R.M., Chung, H., Miao, R., Chou, V., Brody, E., Jiang, X., et al. (2023). Gasdermin-E mediates mitochondrial damage in axons and neurodegeneration. Neuron 111, 1222–1240 e1229. 10.1016/j.neuron.2023.02.019.

29. Pollock, N.M., Fernandes, J.P., Woodfield, J., Moussa, E., Hlavay, B., Branton, W.G., Wuest, M., Mohammadzadeh, N., Schmitt, L., Plemel, J.R., et al. (2024). Gasdermin D activation in oligodendrocytes and microglia drives inflammatory demyelination in progressive multiple sclerosis. Brain Behav Immun 115, 374–393. 10.1016/j.bbi.2023.10.022.

30. Fagerberg, L., Hallstrom, B.M., Oksvold, P., Kampf, C., Djureinovic, D., Odeberg, J., Habuka, M., Tahmasebpoor, S., Danielsson, A., Edlund, K., et al. (2014). Analysis of the human tissue-specific expression by genome-wide integration of transcriptomics and antibody-based proteomics. Mol Cell Proteomics 13, 397–406. 10.1074/mcp.M113.035600.

31. Yue, F., Cheng, Y., Breschi, A., Vierstra, J., Wu, W., Ryba, T., Sandstrom, R., Ma, Z., Davis, C., Pope, B.D., et al. (2014). A comparative encyclopedia of DNA elements in the mouse genome. Nature 515, 355–364. 10.1038/nature13992.

32. Han, X., Wang, R., Zhou, Y., Fei, L., Sun, H., Lai, S., Saadatpour, A., Zhou, Z., Chen, H., Ye, F., et al. (2018). Mapping the Mouse Cell Atlas by Microwell-Seq. Cell 173, 1307. 10.1016/j.cell.2018.05.012.

33. Batiuk, M.Y., Martirosyan, A., Wahis, J., de Vin, F., Marneffe, C., Kusserow, C., Koeppen, J., Viana, J.F., Oliveira, J.F., Voet, T., et al. (2020). Identification of region-specific astrocyte subtypes at single cell resolution. Nat Commun 11, 1220. 10.1038/s41467-019-14198-8.

34. Yang, A.C., Vest, R.T., Kern, F., Lee, D.P., Agam, M., Maat, C.A., Losada, P.M., Chen, M.B., Schaum, N., Khoury, N., et al. (2022). A human brain vascular atlas reveals diverse mediators of Alzheimer’s risk. Nature 603, 885–892. 10.1038/s41586-021-04369-3.

35. Morabito, S., Miyoshi, E., Michael, N., Shahin, S., Martini, A.C., Head, E., Silva, J., Leavy, K., Perez-Rosendahl, M., and Swarup, V. (2021). Single-nucleus chromatin accessibility and transcriptomic characterization of Alzheimer’s disease. Nat Genet 53, 1143–1155. 10.1038/s41588-021-00894-z.

36. Lau, S.F., Cao, H., Fu, A.K.Y., and Ip, N.Y. (2020). Single-nucleus transcriptome analysis reveals dysregulation of angiogenic endothelial cells and neuroprotective glia in Alzheimer’s disease. Proc Natl Acad Sci U S A 117, 25800–25809. 10.1073/pnas.2008762117.

37. Garcia, F.J., Sun, N., Lee, H., Godlewski, B., Mathys, H., Galani, K., Zhou, B., Jiang, X., Ng, A.P., Mantero, J., et al. (2022). Single-cell dissection of the human brain vasculature. Nature 603, 893–899. 10.1038/s41586-022-04521-7.

38. Gerrits, E., Brouwer, N., Kooistra, S.M., Woodbury, M.E., Vermeiren, Y., Lambourne, M., Mulder, J., Kummer, M., Moller, T., Biber, K., et al. (2021). Distinct amyloid-beta and tau-associated microglia profiles in Alzheimer’s disease. Acta Neuropathol 141, 681–696. 10.1007/s00401-021-02263-w.

39. Pineda, S.S., Lee, H., Ulloa-Navas, M.J., Linville, R.M., Garcia, F.J., Galani, K., Engelberg-Cook, E., Castanedes, M.C., Fitzwalter, B.E., Pregent, L.J., et al. (2024). Single-cell dissection of the human motor and prefrontal cortices in ALS and FTLD. Cell 187, 1971–1989 e1916. 10.1016/j.cell.2024.02.031.

40. Wang, Q., Wang, M., Choi, I., Sarrafha, L., Liang, M., Ho, L., Farrell, K., Beaumont, K.G., Sebra, R., De Sanctis, C., et al. (2024). Molecular profiling of human substantia nigra identifies diverse neuron types associated with vulnerability in Parkinson’s disease. Sci Adv 10, eadi8287. 10.1126/sciadv.adi8287.

41. Rogers, C., Fernandes-Alnemri, T., Mayes, L., Alnemri, D., Cingolani, G., and Alnemri, E.S. (2017). Cleavage of DFNA5 by caspase-3 during apoptosis mediates progression to secondary necrotic/pyroptotic cell death. Nat Commun 8, 14128. 10.1038/ncomms14128.

42. Deng, Q., Wu, C., Parker, E., Liu, T.C., Duan, R., and Yang, L. (2024). Microglia and Astrocytes in Alzheimer’s Disease: Significance and Summary of Recent Advances. Aging Dis 15, 1537–1564. 10.14336/AD.2023.0907.

43. Kim, J., Yoo, I.D., Lim, J., and Moon, J.S. (2024). Pathological phenotypes of astrocytes in Alzheimer’s disease. Exp Mol Med 56, 95–99. 10.1038/s12276-023-01148-0.

44. Wieckowski, M.R., Giorgi, C., Lebiedzinska, M., Duszynski, J., and Pinton, P. (2009). Isolation of mitochondria-associated membranes and mitochondria from animal tissues and cells. Nat Protoc 4, 1582–1590. 10.1038/nprot.2009.151.

45. de Brito, O.M., and Scorrano, L. (2008). Mitofusin 2 tethers endoplasmic reticulum to mitochondria. Nature 456, 605–610. 10.1038/nature07534.

46. Brahimi-Horn, M.C., and Mazure, N.M. (2014). Hypoxic VDAC1: a potential mitochondrial marker for cancer therapy. Adv Exp Med Biol 772, 101–110. 10.1007/978-1-4614-5915-6_5.

47. Toutenhoofd, S.L., Foletti, D., Wicki, R., Rhyner, J.A., Garcia, F., Tolon, R., and Strehler, E.E. (1998). Characterization of the human CALM2 calmodulin gene and comparison of the transcriptional activity of CALM1, CALM2 and CALM3. Cell Calcium 23, 323–338. 10.1016/s0143-4160(98)90028-8.

48. Suzuki, J., Kanemaru, K., Ishii, K., Ohkura, M., Okubo, Y., and Iino, M. (2014). Imaging intraorganellar Ca2+ at subcellular resolution using CEPIA. Nat Commun 5, 4153. 10.1038/ncomms5153.

49. Chang-Graham, A.L., Perry, J.L., Strtak, A.C., Ramachandran, N.K., Criglar, J.M., Philip, A.A., Patton, J.T., Estes, M.K., and Hyser, J.M. (2019). Rotavirus Calcium Dysregulation Manifests as Dynamic Calcium Signaling in the Cytoplasm and Endoplasmic Reticulum. Sci Rep 9, 10822. 10.1038/s41598-019-46856-8.

50. Guna, A., Volkmar, N., Christianson, J.C., and Hegde, R.S. (2018). The ER membrane protein complex is a transmembrane domain insertase. Science 359, 470–473. 10.1126/science.aao3099.

51. Zhang, L., Xu, Z., Jia, Z., Cai, S., Wu, Q., Liu, X., Hu, X., Bai, T., Chen, Y., Li, T., et al. (2025). Modulating mTOR-dependent astrocyte substate transitions to alleviate neurodegeneration. Nat Aging 5, 468–485. 10.1038/s43587-024-00792-z.

52. Tian, Y., Shehata, M.A., Gauger, S.J., Ng, C.K.L., Solbak, S., Thiesen, L., Bruus-Jensen, J., Krall, J., Bundgaard, C., Gibson, K.M., et al. (2022). Discovery and Optimization of 5-Hydroxy-Diclofenac toward a New Class of Ligands with Nanomolar Affinity for the CaMKIIalpha Hub Domain. J Med Chem 65, 6656–6676. 10.1021/acs.jmedchem.1c02177.

53. Li, K.L., Huang, H.Y., Ren, H., and Yang, X.L. (2022). Role of exosomes in the pathogenesis of inflammation in Parkinson’s disease. Neural Regen Res 17, 1898–1906. 10.4103/1673-5374.335143.

54. Wang, J., Li, L., Zhang, Z., Zhang, X., Zhu, Y., Zhang, C., and Bi, Y. (2022). Extracellular vesicles mediate the communication of adipose tissue with brain and promote cognitive impairment associated with insulin resistance. Cell Metab 34, 1264–1279 e1268. 10.1016/j.cmet.2022.08.004.

55. Upadhya, R., Zingg, W., Shetty, S., and Shetty, A.K. (2020). Astrocyte-derived extracellular vesicles: Neuroreparative properties and role in the pathogenesis of neurodegenerative disorders. J Control Release 323, 225–239. 10.1016/j.jconrel.2020.04.017.

56. Guttenplan, K.A., Weigel, M.K., Prakash, P., Wijewardhane, P.R., Hasel, P., Rufen-Blanchette, U., Munch, A.E., Blum, J.A., Fine, J., Neal, M.C., et al. (2021). Neurotoxic reactive astrocytes induce cell death via saturated lipids. Nature 599, 102–107. 10.1038/s41586-021-03960-y.

57. Liu, Z., Zhang, H., Liu, S., Hou, Y., and Chi, G. (2023). The Dual Role of Astrocyte-Derived Exosomes and Their Contents in the Process of Alzheimer’s Disease. J Alzheimers Dis 91, 33–42. 10.3233/JAD-220698.

58. Nagai, J., Bellafard, A., Qu, Z., Yu, X., Ollivier, M., Gangwani, M.R., Diaz-Castro, B., Coppola, G., Schumacher, S.M., Golshani, P., et al. (2021). Specific and behaviorally consequential astrocyte G(q) GPCR signaling attenuation in vivo with ibetaARK. Neuron 109, 2256–2274 e2259. 10.1016/j.neuron.2021.05.023.

59. Li, M., Liu, Z., Wu, Y., Zheng, N., Liu, X., Cai, A., Zheng, D., Zhu, J., Wu, J., Xu, L., et al. (2024). In vivo imaging of astrocytes in the whole brain with engineered AAVs and diffusion-weighted magnetic resonance imaging. Mol Psychiatry 29, 545–552. 10.1038/s41380-022-01580-0.

60. Yasuda, R., Hayashi, Y., and Hell, J.W. (2022). CaMKII: a central molecular organizer of synaptic plasticity, learning and memory. Nat Rev Neurosci 23, 666–682. 10.1038/s41583-022-00624-2.

61. Yun, J., Shin, D., Lee, E.H., Kim, J.P., Ham, H., Gu, Y., Chun, M.Y., Kang, S.H., Kim, H.J., Na, D.L., et al. (2025). Temporal Dynamics and Biological Variability of Alzheimer Biomarkers. JAMA Neurol 82, 384–396. 10.1001/jamaneurol.2024.5263.

62. Lee, H.G., Lee, J.H., Flausino, L.E., and Quintana, F.J. (2023). Neuroinflammation: An astrocyte perspective. Sci Transl Med 15, eadi7828. 10.1126/scitranslmed.adi7828.

63. De Schutter, E., Roelandt, R., Riquet, F.B., Van Camp, G., Wullaert, A., and Vandenabeele, P. (2021). Punching Holes in Cellular Membranes: Biology and Evolution of Gasdermins. Trends Cell Biol 31, 500–513. 10.1016/j.tcb.2021.03.004.

64. Zhu, C., Xu, S., Jiang, R., Yu, Y., Bian, J., and Zou, Z. (2024). The gasdermin family: emerging therapeutic targets in diseases. Signal Transduct Target Ther 9, 87. 10.1038/s41392-024-01801-8.

65. Hu, J.J., Liu, X., Xia, S., Zhang, Z., Zhang, Y., Zhao, J., Ruan, J., Luo, X., Lou, X., Bai, Y., et al. (2020). FDA-approved disulfiram inhibits pyroptosis by blocking gasdermin D pore formation. Nat Immunol 21, 736–745. 10.1038/s41590-020-0669-6.

66. Miao, R., Jiang, C., Chang, W.Y., Zhang, H., An, J., Ho, F., Chen, P., Zhang, H., Junqueira, C., Amgalan, D., et al. (2023). Gasdermin D permeabilization of mitochondrial inner and outer membranes accelerates and enhances pyroptosis. Immunity 56, 2523–2541 e2528. 10.1016/j.immuni.2023.10.004.

67. Rizzuto, R., Marchi, S., Bonora, M., Aguiari, P., Bononi, A., De Stefani, D., Giorgi, C., Leo, S., Rimessi, A., Siviero, R., et al. (2009). Ca(2+) transfer from the ER to mitochondria: when, how and why. Biochim Biophys Acta 1787, 1342–1351. 10.1016/j.bbabio.2009.03.015.

68. Belosludtsev, K.N., Dubinin, M.V., Belosludtseva, N.V., and Mironova, G.D. (2019). Mitochondrial Ca2+ Transport: Mechanisms, Molecular Structures, and Role in Cells. Biochemistry (Mosc) 84, 593–607. 10.1134/S0006297919060026.

69. Liu, J., and Yang, J. (2022). Mitochondria-associated membranes: A hub for neurodegenerative diseases. Biomed Pharmacother 149, 112890. 10.1016/j.biopha.2022.112890.

70. Barazzuol, L., Giamogante, F., and Cali, T. (2021). Mitochondria Associated Membranes (MAMs): Architecture and physiopathological role. Cell Calcium 94, 102343. 10.1016/j.ceca.2020.102343.

71. Kuchibhotla, K.V., Lattarulo, C.R., Hyman, B.T., and Bacskai, B.J. (2009). Synchronous hyperactivity and intercellular calcium waves in astrocytes in Alzheimer mice. Science 323, 1211–1215. 10.1126/science.1169096.

72. Korte, N., Barkaway, A., Wells, J., Freitas, F., Sethi, H., Andrews, S.P., Skidmore, J., Stevens, B., and Attwell, D. (2024). Inhibiting Ca(2+) channels in Alzheimer’s disease model mice relaxes pericytes, improves cerebral blood flow and reduces immune cell stalling and hypoxia. Nat Neurosci 27, 2086–2100. 10.1038/s41593-024-01753-w.

73. Princen, K., Van Dooren, T., van Gorsel, M., Louros, N., Yang, X., Dumbacher, M., Bastiaens, I., Coupet, K., Dupont, S., Cuveliers, E., et al. (2024). Pharmacological modulation of septins restores calcium homeostasis and is neuroprotective in models of Alzheimer’s disease. Science 384, eadd6260. 10.1126/science.add6260.

74. Asai, H., Ikezu, S., Tsunoda, S., Medalla, M., Luebke, J., Haydar, T., Wolozin, B., Butovsky, O., Kugler, S., and Ikezu, T. (2015). Depletion of microglia and inhibition of exosome synthesis halt tau propagation. Nat Neurosci 18, 1584–1593. 10.1038/nn.4132.

75. Cheng, L., Vella, L.J., Barnham, K.J., McLean, C., Masters, C.L., and Hill, A.F. (2020). Small RNA fingerprinting of Alzheimer’s disease frontal cortex extracellular vesicles and their comparison with peripheral extracellular vesicles. J Extracell Vesicles 9, 1766822. 10.1080/20013078.2020.1766822.

76. Liu, X., Shen, L., Wan, M., Xie, H., and Wang, Z. (2024). Peripheral extracellular vesicles in neurodegeneration: pathogenic influencers and therapeutic vehicles. J Nanobiotechnology 22, 170. 10.1186/s12951-024-02428-1.

77. Sardar Sinha, M., Ansell-Schultz, A., Civitelli, L., Hildesjo, C., Larsson, M., Lannfelt, L., Ingelsson, M., and Hallbeck, M. (2018). Alzheimer’s disease pathology propagation by exosomes containing toxic amyloid-beta oligomers. Acta Neuropathol 136, 41–56. 10.1007/s00401-018-1868-1.

78. Rabah, Y., Berwick, J.P., Sagar, N., Pasquer, L., Placais, P.Y., and Preat, T. (2025). Astrocyte-to-neuron H(2)O(2) signalling supports long-term memory formation in Drosophila and is impaired in an Alzheimer’s disease model. Nat Metab 7, 321–335. 10.1038/s42255-024-01189-3.

79. Pons-Espinal, M., Blasco-Agell, L., Fernandez-Carasa, I., Andres-Benito, P., di Domenico, A., Richaud-Patin, Y., Baruffi, V., Marruecos, L., Espinosa, L., Garrido, A., et al. (2024). Blocking IL-6 signaling prevents astrocyte-induced neurodegeneration in an iPSC-based model of Parkinson’s disease. JCI Insight 9. 10.1172/jci.insight.163359.

80. Ameroso, D., Meng, A., Chen, S., Felsted, J., Dulla, C.G., and Rios, M. (2022). Astrocytic BDNF signaling within the ventromedial hypothalamus regulates energy homeostasis. Nat Metab 4, 627–643. 10.1038/s42255-022-00566-0.

81. Allen, N.J., Bennett, M.L., Foo, L.C., Wang, G.X., Chakraborty, C., Smith, S.J., and Barres, B.A. (2012). Astrocyte glypicans 4 and 6 promote formation of excitatory synapses via GluA1 AMPA receptors. Nature 486, 410–414. 10.1038/nature11059.

82. Jack, C.R., Jr., Bennett, D.A., Blennow, K., Carrillo, M.C., Dunn, B., Haeberlein, S.B., Holtzman, D.M., Jagust, W., Jessen, F., Karlawish, J., et al. (2018). NIA-AA Research Framework: Toward a biological definition of Alzheimer’s disease. Alzheimers Dement 14, 535–562. 10.1016/j.jalz.2018.02.018.

83. Wolfes, A.C., and Dean, C. (2018). Culturing In Vivo-like Murine Astrocytes Using the Fast, Simple, and Inexpensive AWESAM Protocol. J Vis Exp. 10.3791/56092.

84. Tamashiro, T.T., Dalgard, C.L., and Byrnes, K.R. (2012). Primary microglia isolation from mixed glial cell cultures of neonatal rat brain tissue. J Vis Exp, e3814. 10.3791/3814.

85. Beaudoin, G.M., 3rd, Lee, S.H., Singh, D., Yuan, Y., Ng, Y.G., Reichardt, L.F., and Arikkath, J. (2012). Culturing pyramidal neurons from the early postnatal mouse hippocampus and cortex. Nat Protoc 7, 1741–1754. 10.1038/nprot.2012.099.

86. Dickens, A.M., Tovar, Y.R.L.B., Yoo, S.W., Trout, A.L., Bae, M., Kanmogne, M., Megra, B., Williams, D.W., Witwer, K.W., Gacias, M., et al. (2017). Astrocyte-shed extracellular vesicles regulate the peripheral leukocyte response to inflammatory brain lesions. Sci Signal 10. 10.1126/scisignal.aai7696.

87. Nagy, A., and Delgado-Escueta, A.V. (1984). Rapid preparation of synaptosomes from mammalian brain using nontoxic isoosmotic gradient material (Percoll). J Neurochem 43, 1114–1123. 10.1111/j.1471-4159.1984.tb12851.x.

